# G3BP1 tethers the TSC complex to lysosomes and suppresses mTORC1 in the absence of stress granules

**DOI:** 10.1101/2020.04.16.044081

**Authors:** Mirja T. Prentzell, Ulrike Rehbein, Marti Cadena Sandoval, Ann-Sofie De Meulemeester, Ralf Baumeister, Laura Brohée, Bianca Berdel, Mathias Bockwoldt, Bernadette Carroll, Andreas von Deimling, Constantinos Demetriades, Gianluca Figlia, Alexander M. Heberle, Ines Heiland, Birgit Holzwarth, Lukas A. Huber, Jacek Jaworski, Katharina Kern, Andrii Kopach, Viktor I. Korolchuk, Ineke van ’t Land-Kuper, Matylda Macias, Mark Nellist, Stefan Pusch, Michele Reil, Anja Reintjes, Friederike Reuter, Chloë Scheldeman, Eduard Stefan, Aurelio Teleman, Omar Torres-Quesada, Saskia Trump, Peter de Witte, Teodor Yordanov, Christiane A. Opitz, Kathrin Thedieck

## Abstract

G3BP1 (Ras GTPase-activating protein-binding protein 1) is widely recognized as a core component of stress granules (SG), non-membranous RNA-protein-assemblies required for cellular survival under stress. We report that in the absence of SG, G3BP1 acts as lysosomal anchor of the Tuberous Sclerosis Complex (TSC) protein complex. By tethering the TSC complex to lysosomes, G3BP1 suppresses signaling through the metabolic master regulator mTORC1 (mechanistic target of rapamycin complex 1). Like the known TSC complex subunits, G3BP1 suppresses phenotypes related to mTORC1 hyperactivity in the context of tumors and neuronal dysfunction. Thus, G3BP1 is not only a core component of SG but also a key element of lysosomal TSC-mTORC1 signaling.

**Highlights:** The *bona fide* stress granule component G3BP1

- is a key element of the TSC-mTORC1 signaling axis.
- tethers the TSC complex to lysosomes.
- prevents mTORC1 hyperactivation by metabolic stimuli.
- suppresses mTORC1-driven cancer cell motility and epileptiform activity.

**Graphical Abstract:** 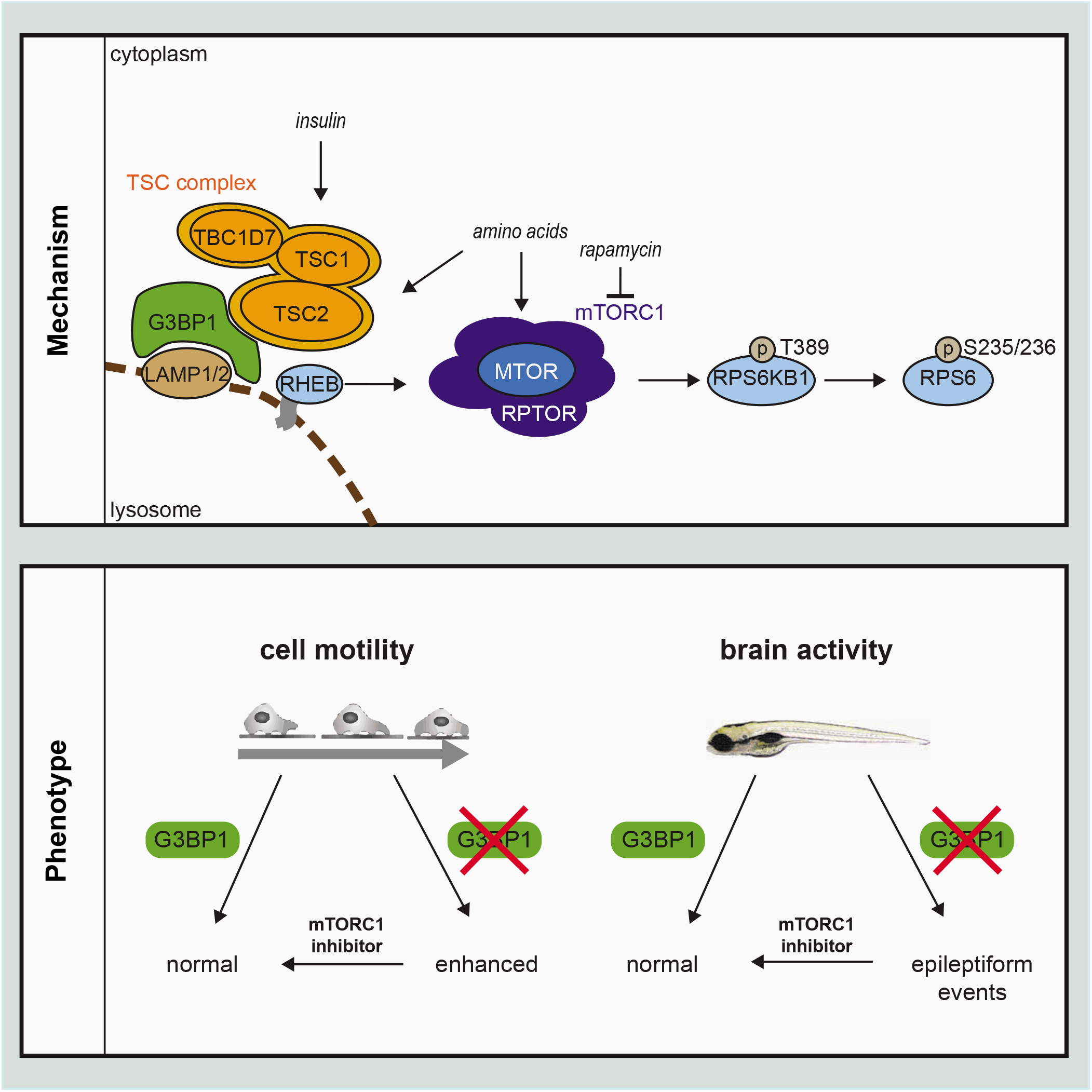

## Introduction

The TSC complex suppresses signaling through the mechanistic target of rapamycin complex 1 (MTOR complex 1, mTORC1), a multiprotein kinase complex that constitutes a metabolic master regulator (Kim and Guan, 2019; Liu and Sabatini, 2020; Tee, 2018). mTORC1 promotes virtually all anabolic processes (Hoxhaj and Manning, 2019; Mossmann et al., 2018), and its hyperactivity is associated with metabolic imbalance and human diseases related to cellular overgrowth, migration, and neuronal excitability (Condon and Sabatini, 2019). Consequently, mTORC1 is recognized as an important driver of tumorigenesis as well as epilepsy (Crino, 2016; LiCausi and Hartman, 2018; Tee et al., 2016). The cause of mTORC1 hyperactivity is often related to a disturbance of the TSC multiprotein complex, known to consist of the subunits TSC1 (hamartin), TSC2 (tuberin), and TBC1D7 (Dibble et al., 2012). The central role of the TSC complex as a tumor suppressor is highlighted by the fact that mutations in the *TSC1* and *TSC2* genes frequently occur in cancer (Huang and Manning, 2008; Kwiatkowski, 2003) and cause tuberous sclerosis complex (TSC), an autosomal dominant disorder, which leads to benign tumors in almost all organ systems and represents one of the most frequent genetic causes of epilepsy (Borkowska et al., 2011; Curatolo et al., 2008; Jozwiak et al., 2019; Marcotte and Crino, 2006; Orlova and Crino, 2010).

In healthy cells, nutritional inputs such as insulin (Menon et al., 2014) and amino acids (Carroll et al., 2016; Demetriades et al., 2014) inhibit the TSC complex, resulting in the derepression of mTORC1 (Kim and Guan, 2019). The TSC complex acts as a GTPase-activating protein (GAP) towards the small GTPase Ras homolog-mTORC1 binding (RHEB) (Garami et al., 2003; Inoki et al., 2003; Tee et al., 2003; Zhang et al., 2003). RHEB directly binds and activates mTORC1 at lysosomes (Avruch et al., 2006; Long et al., 2005; Sancak et al., 2010; Sancak et al., 2007). Thus, RHEB inactivation by the TSC complex restricts the activity of mTORC1 and its multiple anabolic outcomes (Condon and Sabatini, 2019; Kim and Guan, 2019; Rabanal-Ruiz and Korolchuk, 2018). Suppression of RHEB and mTORC1 by the TSC complex takes place at mTORC1’s central signaling platform – the lysosomes (Demetriades et al., 2014; Menon et al., 2014). Thus, recruitment to the lysosomal compartment is crucial for the TSC complex to act on RHEB and mTORC1. The molecular mechanism anchoring mTORC1 at the lysosomes via the LAMTOR-RAG GTPase complex is understood in much detail (Condon and Sabatini, 2019; Kim and Guan, 2019; Rabanal-Ruiz and Korolchuk, 2018). Furthermore, RHEB is known to directly associate with lysosomes via its farnesyl-moiety (Rabanal-Ruiz and Korolchuk, 2018). However, the TSC complex lacks a clear lipid-targeting signal (Kim and Guan, 2019) and it is not yet known how the TSC complex is recruited to lysosomes. Identifying the lysosomal anchor for the TSC complex is important to understand the molecular basis of mTORC1 suppression by the TSC complex. In addition, a tether of the TSC complex is likely to be of high biomedical relevance because of its possible involvement in diseases driven by TSC-mTORC1 dysregulation.

In this study, we identify G3BP1 as a lysosomal tether of the TSC complex. G3BP1 is primarily recognized as an RNA-binding protein that constitutes a core component of SG (Alam and Kennedy, 2019; Reineke and Neilson, 2019), cytoplasmic RNA-protein assemblies formed upon stresses that inhibit protein synthesis (Anderson and Kedersha, 2002; Buchan and Parker, 2009). They are sites of stress-induced mRNA triage that sort transcripts for maintenance or decay and adapt cellular signaling to stress (Anderson and Kedersha, 2008; Anderson et al., 2015). G3BP1 is best described as a SG nucleating protein (Alam and Kennedy, 2019; Kedersha et al., 2016; Mahboubi and Stochaj, 2017; Tourriere et al., 2003), and is widely used as a marker to monitor SG assembly (Kedersha et al., 2008; Moon et al., 2019). G3BP1’s function in SG has also been linked with its involvement in neurological diseases and cancer (Alam and Kennedy, 2019). Only few SG-independent functions of G3BP1 have been proposed. As a protein with RNA binding properties, G3BP1 was suggested to bind to mRNAs of oncogenes and tumor suppressors (Alam and Kennedy, 2019). In its initial report, G3BP1 was proposed to act as a Ras GTPase-activating protein (Ras GAP) binding protein (Gallouzi et al., 1998; Kennedy et al., 2001; Parker et al., 1996) and thus a protein binding property gave rise to its name, although this putative function has since been challenged (Annibaldi et al., 2011). Thus, at present we know little about potential protein binding properties of G3BP1 and putative functions that do not involve SG.

## Results

### G3BP1 inhibits mTORC1 in the absence of stress granules

In a proteomic analysis of the MTOR interactome (Schwarz et al., 2015), we discovered that G3BP1 was significantly enriched with high sequence coverage, along with MTOR and the mTORC1-specific scaffold protein regulatory-associated protein of MTOR complex 1 (RPTOR) (**Figure 1A, S1A, B**). We confirmed the mass spectrometry data by coimmunoprecipitation and found that G3BP1 interacts with MTOR and RPTOR in MCF-7 breast cancer cells (**Figure S1C, D**). G3BP1 is well known for its role in SG assembly (Alam and Kennedy, 2019; Reineke and Neilson, 2019), and SG inhibit mTORC1 (Thedieck et al., 2013; Wippich et al., 2013). To test whether G3BP1 inhibits mTORC1 under conditions that induce SG, we treated MCF-7 cells with arsenite, a frequently used inducer of SG (Anderson et al., 2015). After 30-minute exposure to arsenite, a cytoplasmic punctate pattern of the SG markers G3BP1 and eukaryotic translation initiation factor 3 subunit A (EIF3A) (Kedersha and Anderson, 2007) indicated the presence of SG (**Figure 1B**). Arsenite stress also enhanced the inhibitory phosphorylation of the eukaryotic translation initiation factor 2 alpha (EIF2S1) at Ser51 (**Figure 1C**), which serves as a marker for conditions that inhibit translation and induce SG (Anderson and Kedersha, 2002). In agreement with earlier reports (Heberle et al., 2019; Thedieck et al., 2013; Wang and Proud, 1997), arsenite exposure for 30 minutes enhanced the phosphorylation of the mTORC1 substrate ribosomal protein S6 kinase B1 (RPS6KB1) (Holz and Blenis, 2005) at T389 (RPS6KB1-pT389) (**Figure 1C, E**). G3BP1 knockdown by short hairpin RNA (shG3BP1, **Figure 1D, S1E**) reduced the G3BP1 protein levels, but did not alter RPS6KB1-T389 phosphorylation (**Figure 1C, E**). Also, upon arsenite exposure for various time periods up to 60 minutes, G3BP1 knockdown by shRNA or siRNA (**Figure S1E**) did not alter RPS6KB1-pT389 levels (**Figure S1F-K**). Therefore, we conclude that in the presence of SG, G3BP1 does not affect mTORC1 activity.

**Figure 1.**
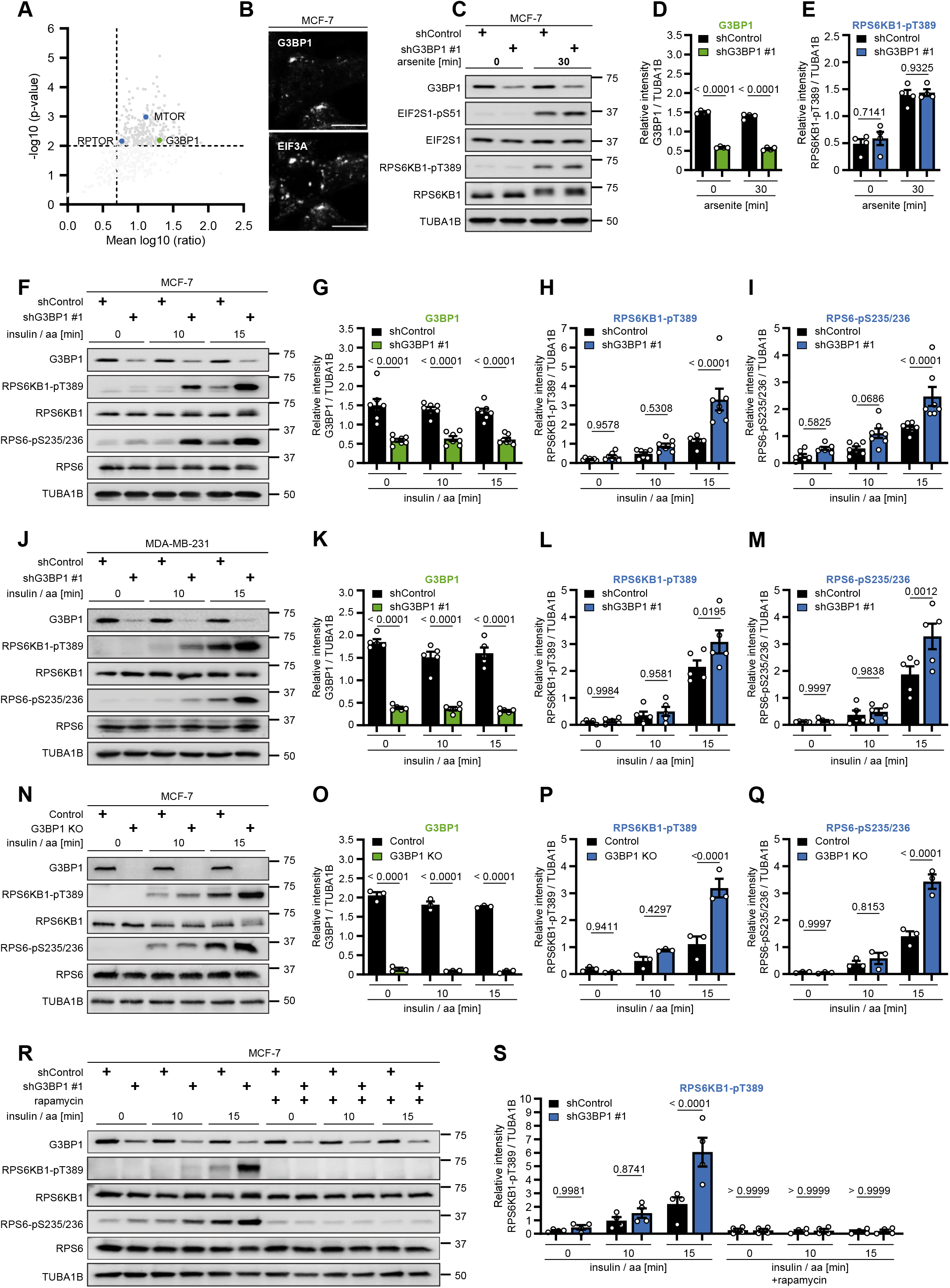
G3BP1 suppresses mTORC1 reactivation by insulin and nutrients. **(A)** Re-analysis of the MTOR interactome data reported by Schwarz et al. (2015). Volcano plot showing the mean log_10_ ratios of proteins detected by tandem mass spectrometry in MTOR *versus* mock immunoprecipitation (IP) experiments. Proteins quantified in at least two out of three biological replicates were plotted against the negative log_10_ p-value (Student’s t-test). Proteins with a mean ratio > 5 and a p-value < 0.01 (sector highlighted in dark gray) were considered significantly enriched. G3BP1 is marked in green, the mTORC1 core components MTOR and RPTOR are marked in blue. **(B)** Immunofluorescence (IF) analysis of MCF-7 cells, serum starved, and treated with 500 μM arsenite for 30 minutes. Cells were stained with G3BP1 and EIF3A antibodies. Scale bar 10 μm. Representative images shown for n = 3 biological replicates. **(C)** MCF-7 cells stably transduced with shG3BP1 #1 or shControl were serum starved and treated with 500 μM arsenite for 30 minutes. Data shown are representative of n = 4 biological replicates. **(D)** Quantitation of G3BP1 immunoblot data shown in (**C**). Data are shown as the mean ± SEM and overlaid with the single data points represented as dot plots. G3BP1 levels (black and green bars) were compared between shControl and shG3BP1 #1 cells, using a one-way ANOVA followed by a Sidak’s multiple comparisons test across n = 4 biological replicates. p-values of the Sidak’s multiple comparisons test are presented above the bar graphs. **(E)** Quantitation of RPS6KB1-pT389 immunoblot data shown in (**C**). RPS6KB1-pT389 levels (black and blue bars) are represented and compared between shControl and shG3BP1 #1 cells as described in (**D**). **(F)** shG3BP1 #1 or shControl MCF-7 cells were serum and amino acid starved, and stimulated with 100 nM insulin and amino acids (insulin / aa) for the indicated time periods. Data shown are representative of n = 7 biological replicates. **(G)** Quantitation of G3BP1 immunoblot data shown in (**F**). Data are shown as the mean ± SEM and overlaid with the single data points represented as dot plots. G3BP1 levels (black and green bars) were compared between shControl and shG3BP1 #1 cells, using a one-way ANOVA followed by a Sidak’s multiple comparisons test across n = 7 biological replicates. p-values of the Sidak’s multiple comparisons test are presented above the corresponding bar graphs. **(H)** Quantitation of RPS6KB1-pT389 immunoblot data shown in (**F**). RPS6KB1-pT389 levels (black and blue bars) are represented and compared between shControl and shG3BP1 #1 cells as described in (**G**). **(I)** Quantitation of RPS6-pS235/236 immunoblot data shown in (**F**). RPS6-pS235/236 levels (black and blue bars) are represented and compared between shControl and shG3BP1 #1 cells as described in (**G**). **(J)** shG3BP1 #1 or shControl MDA-MB-231 cells were serum and amino acid starved, and stimulated with 100 nM insulin / aa for the indicated time periods. Data shown are representative of n = 5 biological replicates. **(K)** Quantitation of G3BP1 immunoblot data shown in (**J**). Data are shown as the mean ± SEM and overlaid with the single data points represented as dot plots. G3BP1 levels (black and green bars) were compared between shControl and shG3BP1 #1 cells, using a one-way ANOVA followed by a Sidak’s multiple comparisons test across n = 5 biological replicates. p-values of the Sidak’s multiple comparisons test are presented above the corresponding bar graphs. **(L)** Quantitation of RPS6KB1-pT389 immunoblot data shown in (**J**). RPS6KB1-pT389 levels (black and blue bars) are represented and compared between shControl and shG3BP1 #1 cells as described in (**K**). **(M)** Quantitation of RPS6-pS235/236 immunoblot data shown in (**J**). RPS6-pS235/236 levels (black and blue bars) are represented and compared between shControl and shG3BP1 #1 cells as described in (**K**). **(N)** G3BP1 CRISPR/Cas9 KO or control MCF-7 cells were serum and amino acid starved, and stimulated with 100 nM insulin / aa for the indicated time periods. Data shown are representative of n = 3 biological replicates. **(O)** Quantitation of G3BP1 immunoblot data shown in (**N**). Data are shown as the mean ± SEM and overlaid with the single data points represented as dot plots. G3BP1 levels (black and green bars) were compared between control and G3BP1 KO cells, using a one-way ANOVA followed by a Sidak’s multiple comparisons test across n = 3 biological replicates. p-values of the Sidak’s multiple comparisons test are presented above the corresponding bar graphs. **(P)** Quantitation of RPS6KB1-pT389 immunoblot data shown in (**N**). RPS6KB1-pT389 levels (black and blue bars) are represented and compared between control and G3BP1 KO cells as described in (**O**). **(Q)** Quantitation of RPS6-pS235/236 immunoblot data shown in (**N**). RPS6-pS235/236 levels (black and blue bars) are represented and compared between control and G3BP1 KO cells as described in (**O**). **(R)** shG3BP1 #1 or shControl MCF-7 cells were serum and amino acid starved, and stimulated with 100 nM insulin / aa for the indicated time periods. The rapamycin treatment started 30 minutes before insulin / aa stimulation. Data shown are representative of n = 4 biological replicates. **(S)** Quantitation of RPS6KB1-pT389 immunoblot data shown in (**R**). Data are shown as the mean ± SEM and overlaid with the single data points represented as dot plots. RPS6KB1-pT389 (black and blue bars) was compared between shControl and shG3BP1 #1 cells, using a one-way ANOVA followed by a Sidak’s multiple comparisons test across n = 4 biological replicates. p-values of the Sidak’s multiple comparisons test are presented above the corresponding bar graphs.

We next tested whether G3BP1 influences mTORC1 activity under conditions that are not associated with the formation of SG. For this purpose, we starved MCF-7 cells and then restimulated them with insulin and amino acids to activate metabolic signaling through mTORC1. G3BP1 was targeted by two different shRNA sequences (**Figure S1E**). Insulin and amino acids enhanced phosphorylation of RPS6KB1-T389 and of its substrate ribosomal protein S6 (RPS6-pS235/236) (Pende et al., 2004), indicative of mTORC1 activation (**Figure 1F, H, I and S2A, C, D**). Of note, G3BP1 knockdown led to a further increase in RPS6KB1-pT389 and RPS6-pS235/236 (**Figure 1F-I and S2A-D**). In triple negative MDA-MB-231 cells (Neve et al., 2006), shG3BP1-mediated knockdown enhanced RPS6KB1-pT389 and RPS6-pS235/236 as well (**Figure 1J-M and S2E-H**). Targeting G3BP1 by siRNA knockdown (**Figure S2I-L**) or CRISPR/Cas9 knockout (**Figure 1N-Q and S2M**) also resulted in RPS6KB1-T389 and RPS6-S235/236 hyperphosphorylation. To test whether enhanced RPS6KB1-pT389 and RPS6-pS235/236 in G3BP1-deficient cells is mediated by mTORC1, we used the allosteric mTORC1 inhibitor rapamycin, which potently inhibited RPS6KB1-T389 and RPS6-S235/236 phosphorylation in G3BP1-deficient cells (**Figure 1R, S**). Thus, we conclude that G3BP1 restricts mTORC1 activation by amino acids and insulin.

As G3BP1 is a core component of SG (Alam and Kennedy, 2019; Reineke and Neilson, 2019), which are known to inhibit mTORC1 under stress (Thedieck et al., 2013; Wippich et al., 2013), we wondered whether SG were also present in metabolically stimulated cells. To test this, we performed immunofluorescence (IF) experiments in which we analysed the distribution patterns of endogenous G3BP1 and EIF3A in cells stimulated with insulin and amino acids, or upon arsenite stress as a positive control (**Figure S2N, O**). G3BP1 knockdown reduced G3BP1 levels, as expected, but SG remained present in the arsenite treated cells (further discussed below). While arsenite induced SG, no puncta indicative of SG became visible in insulin and amino acid stimulated cells, and G3BP1 and EIF3A were distributed throughout the cytoplasm. Thus, mTORC1 inhibition by G3BP1 occurs in the absence of SG.

### G3BP1 resides at lysosomes

To identify the subcellular compartment where G3BP1 acts in the absence of SG, we fractionated lysates of starved cells by sucrose density gradient centrifugation (**Figure 2A**). The TSC complex components TSC1, TSC2, and TBC1D7 were predominantly detected in the fractions containing the lysosome associated membrane proteins 1 and 2 (LAMP1, LAMP2) (Eskelinen, 2006). This is in line with earlier biochemical and IF-based studies demonstrating that the TSC complex inhibits mTORC1 at lysosomes when cells lack amino acids or growth factors (Carroll et al., 2016; Demetriades et al., 2014; Menon et al., 2014). In the absence of SG inducers, G3BP1 exhibits a ubiquitous cytoplasmic localization (**Figure S2N**) (Irvine et al., 2004), but so far no specific sub-cellular enrichment has been identified. We found that G3BP1 resides in the lysosomal fractions (**Figure 2A**). Thus, in the absence of SG, G3BP1 co-fractionates with the TSC complex and lysosomal proteins. We demonstrated the lysosomal association of G3BP1 further *in situ* by proximity ligation assays (PLA) of G3BP1 with LAMP1 (**Figure 2B, C**). Thus, we propose that G3BP1 localizes to lysosomes, in close proximity to LAMP1.

**Figure 2.**
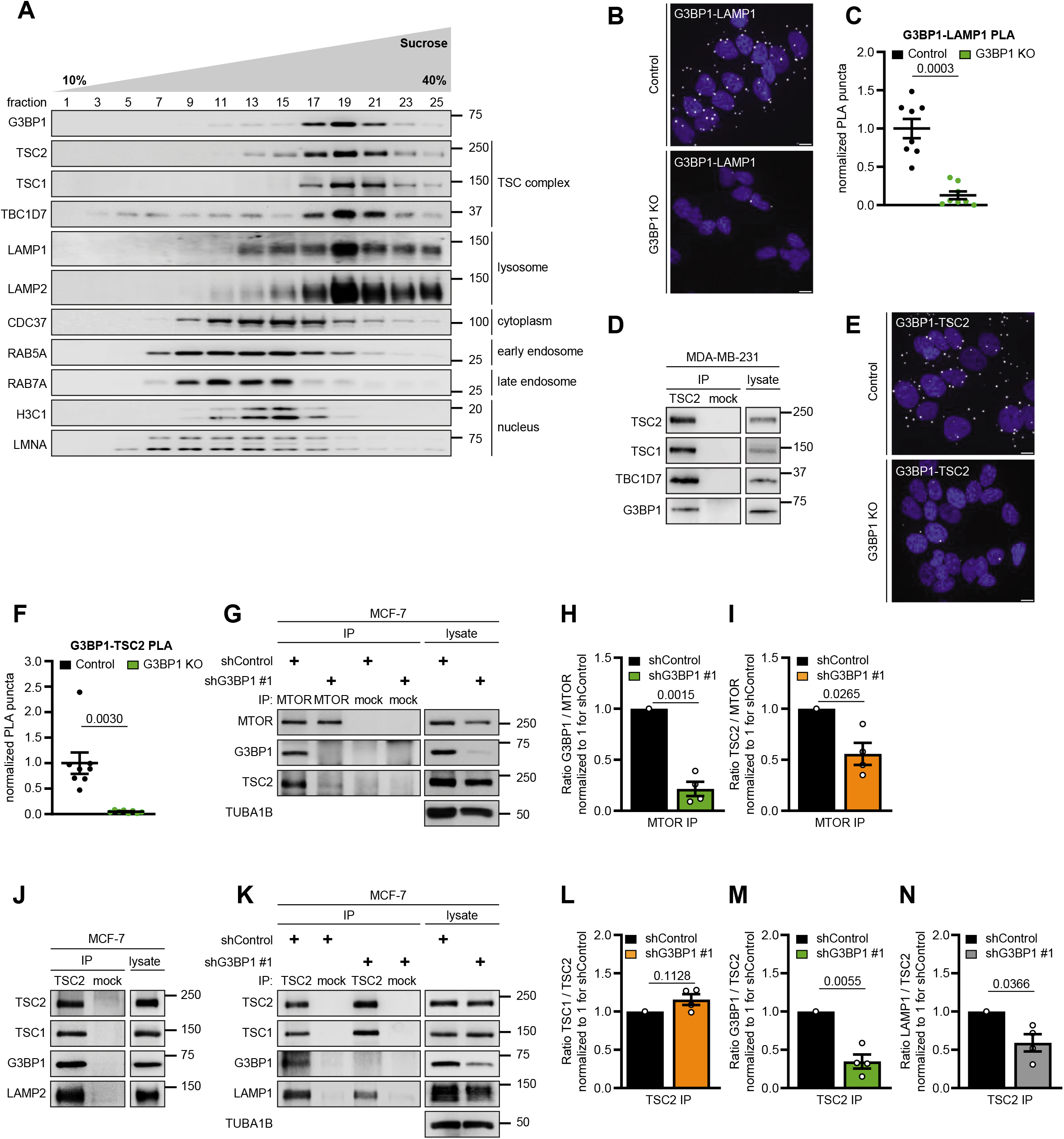
G3BP1 resides at lysosomes. **(A)** Separation of MCF-7 cell lysates by sucrose density gradient. Cells were serum and amino acid starved. Samples were separated in a 10 to 40% sucrose gradient and analyzed by immunoblot. TSC2, TSC1 and TBC1D7, TSC complex; LAMP1 and LAMP2, lysosomal proteins; CDC37, cytoplasmic marker; RAB5A and RAB7A, early and late endosomal marker proteins, respectively; Histone H3 and LMNA, nuclear markers. Data shown are representative of n = 3 biological replicates. **(B)** PLA analysis of G3BP1-LAMP1 association in serum and amino acid starved MCF-7 G3BP1 CRISPR/Cas9 KO and control cells. Data shown are representative of n = 3 biological replicates. PLA puncta, white dots; nuclei, blue (DAPI). Scale bar 10 μm. **(C)** Quantitation of data shown in (**B**). Data are shown as the mean ± SEM and overlaid with the single data points represented as dot plots. The number of PLA puncta per cell was normalized to 1 for the mean of control cells. Control and G3BP1 KO cells were compared using a paired two-tailed Student’s t-test across n = 8 technical replicates. The p-value is presented above the graph. Data shown are representative of n = 3 biological replicates. **(D)** IPs from MDA-MB-231 cells with antibodies against TSC2 (TSC2 #1) or mock (mouse IgG). Data shown are representative of n = 3 biological replicates. **(E)** PLA analysis of G3BP1-TSC2 association in serum and amino acid starved MCF-7 G3BP1 CRISPR/Cas9 KO and control cells. Data shown are representative of n = 4 biological replicates. PLA puncta, white dots; nuclei, blue (DAPI). Scale bar 10 μm. **(F)** Quantitation of data shown in (**E**). Data are shown as the mean ± SEM and overlaid with the single data points represented as dot plots. The number of PLA puncta per cell was normalized to 1 for the mean of control cells. Control and G3BP1 KO cells were compared using a paired two-tailed Student’s t-test across n = 8 technical replicates. The p-value is presented above the graph. Data shown are representative of n = 4 biological replicates. **(G)** IPs from MCF-7 cells with antibodies against MTOR or mock (rat IgG). shG3BP1 #1 or shControl cells were serum and amino acid starved, and stimulated with 100 nM insulin / aa for 15 minutes. Data shown are representative of n = 4 biological replicates. **(H)** Quantitation of G3BP1 immunoblot data shown in (**G**). The ratios of G3BP1/ MTOR (black and green bars) are shown as the mean ± SEM and overlaid with the single data points represented as dot plots. All data were normalized to 1 for shControl. shControl and shG3BP1 #1 cells were compared using a paired two-tailed Student’s t-test across n = 4 biological replicates. p-values are presented above the corresponding bar graphs. **(I)** Quantitation of TSC2 immunoblot data shown in (**G**). The ratios of TSC2/ MTOR (black and orange bars) are represented and compared between shControl and shG3BP1 #1 cells as described in (**H**). **(J)** IPs from MCF-7 cells with antibodies against TSC2 (TSC2 #2 or #3) or mock (rabbit IgG). Data shown are representative of n = 3 biological replicates. **(K)** IPs from MCF-7 cells with antibodies against TSC2 (TSC2 #2) or mock (rabbit IgG). shG3BP1 #1 or shControl cells were serum and amino acid starved, and stimulated with 100 nM insulin / aa for 15 minutes. Data shown are representative of n = 4 biological replicates. **(L)** Quantitation of TSC1 immunoblot data shown in (**K**). The ratios of TSC1/ TSC2 (black and orange bars) are shown as the mean ± SEM and overlaid with the single data points represented as dot plots. All data were normalized to 1 for shControl. shControl and shG3BP1 #1 cells were compared using a paired two-tailed Student’s t-test across n = 4 biological replicates. p-values are presented above the corresponding graphs. **(M)** Quantitation of G3BP1 immunoblot data shown in (**K**). The ratios of G3BP1/ MTOR (black and green bars) are represented and compared between shControl and shG3BP1 #1 cells as described in (**L**). **(N)** Quantitation of LAMP1 immunoblot data shown in (**K**). The ratios of LAMP1/ TSC2 (black and grey bars) are represented and compared between shControl and shG3BP1 #1 cells as described in (**L**).

### G3BP1 tethers the TSC complex to lysosomes

G3BP1 co-fractionates with the TSC complex (**Figure 2A**), and we investigated whether they physically interact. Indeed, as TSC1 and TBC1D7, G3BP1 co-immunoprecipitated with TSC2 (**Figure 2D**). PLA supported the association of G3BP1 with TSC2 *in situ* (**Figure 2E, F**), indicative of a distance between the two proteins of less than 40 nm (Debaize et al., 2017). Thus, G3BP1 is a novel interactor of the TSC complex.

Interestingly, TSC2 and G3BP1 both co-immunoprecipitated with MTOR (**Figure 2G-I**). This physical interaction likely reflects the lysosomal localization of G3BP1, the TSC complex, and mTORC1. G3BP1 deficiency significantly reduced TSC2-MTOR association (**Figure 2G-I**), suggesting that G3BP1 is required for the TSC complex to act on MTOR. As a likely scenario, we hypothesized that G3BP1 might inhibit mTORC1 by mediating the localization of the TSC complex to lysosomes. We first tested this assumption in IPs of TSC2, which co-immunoprecipitated not only TSC1 and G3BP1 but also the lysosomal proteins LAMP1 and 2 (**Figure 2J, K**). G3BP1 deficiency significantly reduced the physical interaction of TSC2 with LAMP1 (**Figure 2K-N**), indicative of a role of G3BP1 as a lysosomal tether for the TSC complex.

To further address the requirement of G3BP1 for the lysosomal localization of the TSC complex, we analyzed TSC2-LAMP2 association *in situ* by PLA in G3BP1-proficient and -deficient cells (**Figure 3A, B**). As reported earlier (Carroll et al., 2016; Demetriades et al., 2014; Demetriades et al., 2016; Menon et al., 2014), TSC2-LAMP2 association was highest in starved cells and decreased upon stimulation with amino acids and insulin. In starved cells, G3BP1 knockdown significantly reduced TSC2-LAMP2 association, to a similar level as observed upon insulin and amino acid stimulation. This result was corroborated by IF analysis of TSC2 and LAMP1 co-localization in G3BP1 CRISPR/Cas9 KO cells (**Figure 3C, D**). G3BP1 KO reduced TSC2-LAMP1 co-localization in starved cells to the same extent as metabolic stimulation with insulin and amino acids. Thus, G3BP1 mediates lysosomal localization of the TSC complex in cells deprived of insulin and nutrients. In agreement with this, we observed a significant induction of RPS6KB1 and RPS6 phosphorylation not only in metabolically stimulated cells, but also when inhibiting G3BP1 in starved cells (**Figure 3E-H**). The signals under starvation had been quenched in earlier experiments by the much stronger signals upon metabolic stimulation (**Figure 1F-I**). Thus, we propose that in G3BP1 deficient cells, impaired lysosomal recruitment of the TSC complex under starvation enhances mTORC1 activity. This results in faster phosphorylation of mTORC1 substrates upon metabolic stimuli.

**Figure 3.**
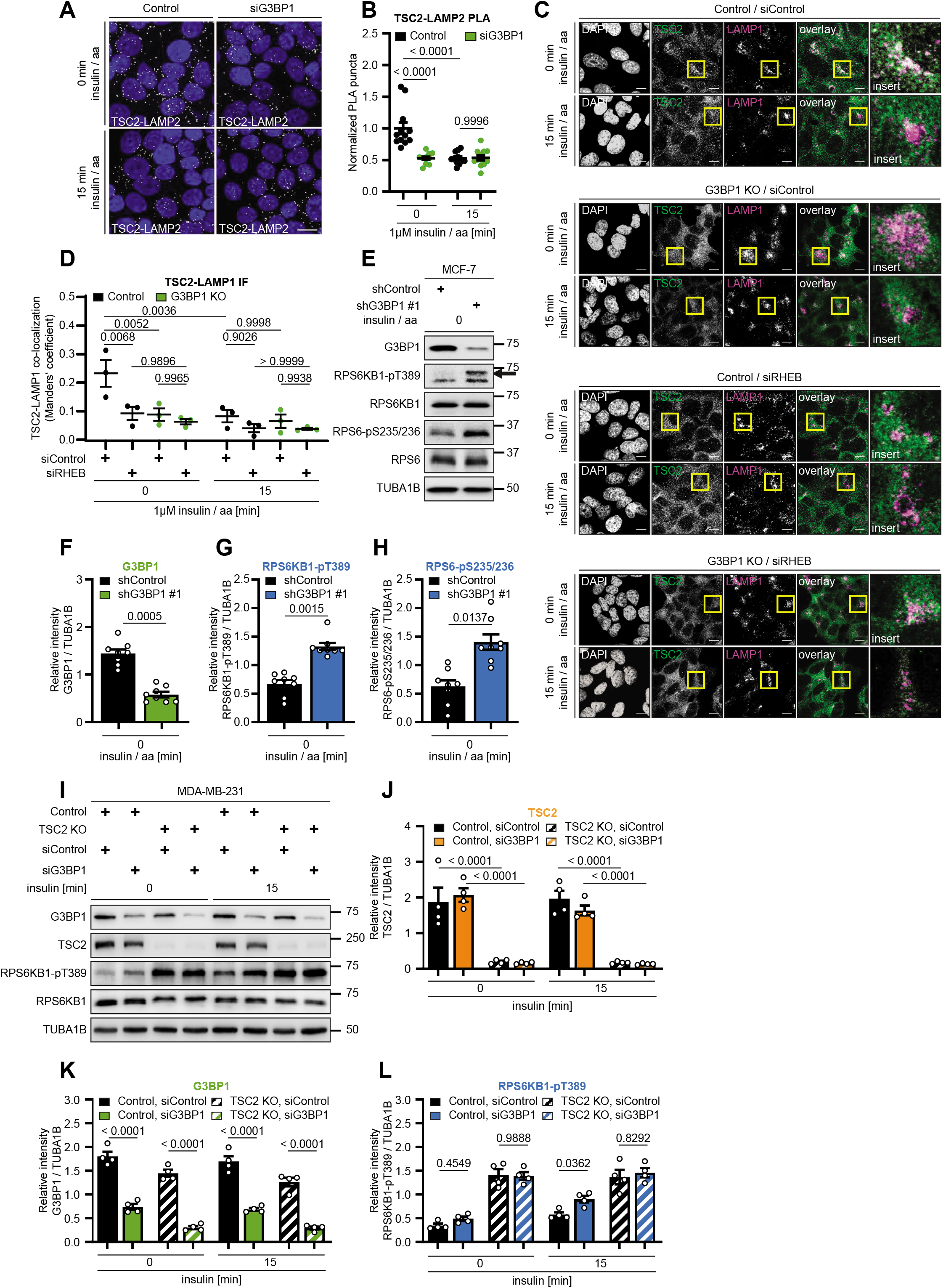
G3BP1 tethers the TSC to lysosomes. **(A)** PLA analysis of TSC2-LAMP2 association in si*Renilla* luciferase (Control) or siG3BP1 transfected MCF-7 cells. Cells were serum and amino acid starved, and stimulated with 1 μM insulin / aa for 15 minutes. Data shown are representative of n = 4 biological replicates. PLA puncta, white dots; nuclei, blue (DAPI). Scale bar 100 μm. **(B)** Quantitation of data shown in (**A**). Data are shown as the mean ± SEM and overlaid with the single data points represented as dot plots. The number of PLA puncta per field was normalized to the number of DAPI-positive nuclei, and the mean of serum and amino acid starved control cells was set to 1. Control and siG3BP1 cells were compared using a one-way ANOVA followed by a Sidak’s multiple comparisons test across n = 12 technical replicates. p-values are presented above the graphs. Data shown are representative of n = 4 biological replicates. **(C)** IF analysis of LAMP1-TSC2 co-localization in MCF-7 G3BP1 CRISPR/Cas9 KO and control cells. Cells transfected with either siControl or siRHEB were serum and amino acid starved, and stimulated with 1 μM insulin / aa for 15 minutes. Scale bar 10 μm. White regions in overlay, co-localization of LAMP1 and TSC2. Insert, magnification of the area in the yellow square. Nuclei were stained with DAPI. Images are representative of n = 4-5 distinct fields of view/ replicate and n = 3 biological replicates. **(D)** Quantitation of data shown in (**C**). The Manders’ correlation coefficient for TSC2 and LAMP1 is represented as mean ± SEM, which was calculated across n = 3 biological replicates with 4-5 distinct fields of view in each. The single data points are overlaid as dot plots. The differences among all conditions were assessed by a one-way ANOVA followed by a Sidak’s multiple comparisons test. p-values are presented above the graphs. **(E)** shG3BP1 #1 or shControl MCF-7 cells were serum and amino acid starved. The arrow indicates the specific RPS6KB1-pT389 signal. Data shown are representative of n = 8 biological replicates. **(F)** Quantitation of G3BP1 immunoblot data shown in (**E**). Data are shown as the mean ± SEM and overlaid with the single data points represented as dot plots. G3BP1 levels (black and green bars) were compared between shControl and shG3BP1 #1 cells, using a paired two-tailed Student’s t-test across n = 8 biological replicates. p-values are presented above the corresponding bar graphs. **(G)** Quantitation of RPS6KB1-pT389 immunoblot data shown in (**E**). RPS6KB1-pT389 levels (black and blue bars) are represented and compared between shControl and shG3BP1 #1 cells as described in (**F**). **(H)** Quantitation of RPS6-pS235/236 immunoblot data shown in (**E**). RPS6-pS235/236 levels (black and blue bars) are represented and compared between shControl and shG3BP1 #1 cells as described in (**F**). **(I)** Control or TSC2 CRISPR/Cas9 KO MDA-MB-231 cells, transfected with either siControl or siG3BP1 were serum starved, and stimulated with 100 nM insulin for 15 minutes. Data shown are representative of n = 4 biological replicates. **(J)** Quantitation of TSC2 immunoblot data shown in (**I**). Data are shown as the mean ± SEM and overlaid with the single data points represented as dot plots. TSC2 levels (black and orange bars) were compared between control and TSC2 KO cells using a one-way ANOVA followed by a Sidak’s multiple comparisons test across n = 4 biological replicates. p-values are presented above the corresponding bar graphs. **(K)** Quantitation of G3BP1 immunoblot data shown in (**I**). Data are shown as the mean ± SEM and overlaid with the single data points represented as dot plots. G3BP1 levels (black and green bars) were compared between siControl and siG3BP1 in control or TSC2 KO cells, using a one-way ANOVA followed by a Sidak’s multiple comparisons test across n = 4 biological replicates. p-values are presented above the corresponding bar graphs. **(Q)** Quantitation of RPS6KB1-pT389 immunoblot data shown in (**I**). RPS6KB1-pT389 levels (black and blue bars) are represented and compared between siControl and siG3BP1 in control or TSC2 KO cells as described in (**K**).

The TSC complex acts as a GAP for RHEB, and their interaction contributes to the lysosomal localization of the TSC complex (Carroll et al., 2016; Menon et al., 2014). A similar function has been suggested for RAG GTPases upon depletion of amino acids (Demetriades et al., 2014). To test whether the mechanisms via which G3BP1 and RHEB target the TSC complex to lysosomes are interdependent, we compared the effects of RHEB and G3BP1 inhibition on TSC2-LAMP1 co-localization (**Figure 3C, D**). We found that G3BP1 KO and RHEB knockdown reduced TSC2-LAMP1 co-localization to a similar extent, and they did not exert an additive effect. Thus, G3BP1 and RHEB are both necessary for the lysosomal recruitment of the TSC. In other words, the association with its target GTPase is not sufficient for the lysosomal localization of the TSC complex as it requires G3BP1 as an additional tether.

### G3BP1 suppresses mTORC1 via the TSC complex

Our data so far showed that G3BP1 recruits the TSC complex to lysosomes and inhibits mTORC1. We tested next if G3BP1’s function as an mTORC1 suppressor depends on the TSC complex. For this purpose, we conducted an epistasis experiment in which we analyzed the effect of G3BP1 inhibition on mTORC1 activity in the presence or absence of TSC2 (**Figure 3I-L**). We had previously stimulated cells with insulin and amino acids, as they both signal through the TSC complex (Carroll et al., 2016; Demetriades et al., 2014; Demetriades et al., 2016). Amino acids also signal to mTORC1 via TSC complex-independent routes (Liu and Sabatini, 2020; Rabanal-Ruiz and Korolchuk, 2018). Thus, for the epistasis experiment, we opted to stimulate the cells exclusively with insulin to only assess mTORC1 inactivation via the TSC complex. As expected, RPS6KB1-T389 was hyperphosphorylated to a similar extent in starved or insulin-stimulated TSC2 CRISPR/Cas9 KO cells, as the TSC complex was absent. G3BP1 inhibition induced RPS6KB1-T389 hyperphosphorylation in starved control cells, and this effect was further enhanced by insulin. However, G3BP1 inhibition did not further enhance RPS6KB1-pT389 in TSC2 KO cells (**Figure 3I, L**). Thus, we propose that G3BP1 and the TSC complex act in the same signaling pathway to suppress mTORC1.

### TSC2 mediates the formation of the G3BP1-TSC complex

To further understand the molecular makeup of the TSC-G3BP1 complex, we next determined which of the known subunits mediates G3BP1 binding. For this purpose, we analyzed G3BP1 binding to TSC1 in TSC2 KO or control cells (**Figure 4A**). TSC2 KO resulted in a complete loss of G3BP1 from the TSC1-TBC1D7 complex, indicating that G3BP1 binds TSC2. We next aimed to detOf note, overexpression of Cermine the TSC2-binding domain of G3BP1. A C-terminal fragment of G3BP1, consisting of amino acids 333-466, co-immunoprecipitated with GFP-TSC2 to a similar extent as full-length G3BP1 (**Figure 4B, C**). This indicates that G3BP1 binds TSC2 mainly via its C-terminus, harboring RNA recognition motifs (RRM) and arginine-glycine-glycine repeats (RGG) (Tourriere et al., 2003) (**Figure S1A**). In contrast, the middle part (amino acids 183-332; containing the proline rich domain) and the N-terminal region (amino acids 1-182; harboring the NTF2-like domain) of G3BP1 exhibited faint or no interaction with TSC2, respectively. Thus, we conclude that the G3BP1-TSC2 interaction is mainly mediated by G3BP1’s C-terminus. Of note, overexpression of C-terminal G3BP1 (lacking the NTF2-like domain) cannot induce SG (Reineke and Lloyd, 2015; Takahashi et al., 2013; Tourriere et al., 2003; Zhang et al., 2019). This further supports that C-terminal G3BP1 interacts with TSC2 in a SG-independent manner. We propose that the C-terminal region of G3BP1 has a dual function in mediating the interaction with RNA in SG (Reineke and Neilson, 2019), and with the TSC complex under non-stress conditions.

**Figure 4.**
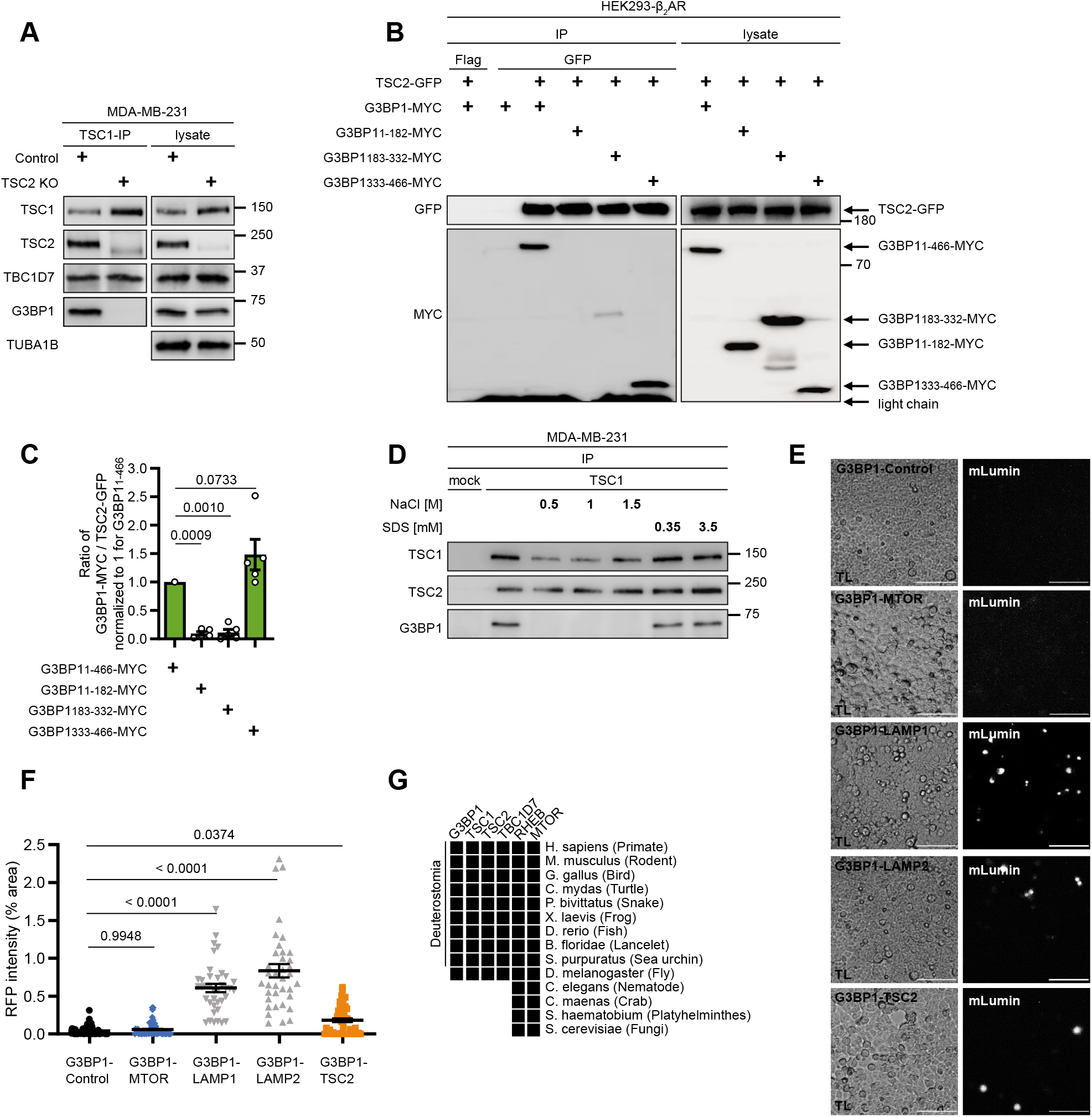
Properties of the TSC2-G3BP1 interaction. **(A)** IPs from TSC2 KO or control MDA-MB-231 cells with antibodies against TSC1 (TSC1 #1) or mock (rabbit IgG). Data shown are representative of n = 3 biological replicates. **(B)** IPs with antibodies against GFP or Flag from HEK293-β2AR cells co-transfected with TSC2-GFP and full length G3BP1_1-466_-MYC or truncated G3BP1-MYC versions (G3BP1_1-182_, G3BP1_183-332_, G3BP1_333-466_). Data shown are representative of n = 5 biological replicates. **(C)** Quantitation of G3BP1-myc immunoblot data shown in (**B**). The ratios of G3BP1-myc/ TSC2-GFP are shown. All data were normalized to 1 for G3BP1_1-466_. Data are shown as the mean ± SEM and overlaid with the single data points represented as dot plots. The ratios were compared between full length G3BP1_1-466_ and the truncated versions (G3BP1_1-182_, G3BP1_183-332_, G3BP1_333-466_), using a one-way ANOVA followed by a Sidak’s multiple comparisons test across n = 5 biological replicates. p-values are presented above the corresponding bar graphs. **(D)** Resistance of the TSC-G3BP1 complex against high salt or detergent. IPs from MDA-MB-231 cells with antibodies against TSC1 (TSC1 #2) or mock (mouse IgG) were incubated with the indicated concentrations of NaCl and SDS. Data shown are representative of n = 3 biological replicates. **(E)** Bimolecular fluorescence complementation (BiFC) analysis of HEK293T cells transfected with plasmids carrying G3BP1 fused to a C-terminal mLumin fragment, together with an N-terminal mLumin fragment only (Control), or an N-terminal mLumin fragment fused to MTOR, LAMP1, LAMP2 or TSC2. Scale bar 100 μm. One representative image of each channel is shown for at least n = 3 biological replicates. A scheme depicting the fusion constructs is shown in **Figure S3A**. **(F)** Quantitation of data shown in (**E**). Data are shown as the mean ± SEM and overlaid with the single data points represented as dot plots. The percentages of mLumin fluorescence intensity (RFP) / picture were compared between G3BP1-Control and the different plasmid combinations (G3BP1-MTOR, G3BP1-LAMP1, G3BP1-LAMP2, G3BP1-TSC2), using a oneway ANOVA followed by a Sidak’s multiple comparisons test across at least 22 biological fields of view from at least n = 3 biological replicates. p-values are presented above the corresponding bar graphs. **(G)** Excerpt of a phylogenetic Blast analysis of G3BP1, TSC1, TSC2, TBC1D7, RHEB, and MTOR. A black square depicts the presence of the protein in the respective species, based on blastp+ search against NCBI nr protein database (e-value < 1e-30; for details see materials and methods).

The known members of the TSC complex are resistant to high salt and detergent conditions, indicative of their high binding affinity (Dibble et al., 2012; Nellist et al., 1999). The complex formed by TSC1, TSC2, and TBC1D7 remains stable at 1.5 M NaCl and 0.1% (3.5 mM) sodium dodecyl sulfate (SDS) (Dibble et al., 2012). To obtain information about the affinity of the TSC2-G3BP1 interaction, we incubated TSC1 IPs with up to 1.5 M NaCl or up to 3.5 mM SDS (**Figure 4D**). While the TSC1-TSC2 interaction was resistant to 1.5 M NaCl, the binding to G3BP1 was lost at 0.5 M NaCl. This salt sensitivity suggests that the complex is formed via electrostatic interactions. In line with this, the G3BP1 C-terminus harbors an intrinsically disordered region (IDR) (Panas et al., 2019), which – as is typical for IDRs (Forman-Kay and Mittag, 2013) – contains a high density of positively charged arginine residues that mediate electrostatic interactions. Importantly, the interaction of TSC2 with G3BP1 was highly stable against denaturation by SDS that preferentially disrupts hydrophobic interactions at the concentration used in this experiment (3.5 mM) (Hojgaard et al., 2018). Thus, upon SDS exposure, G3BP1 exhibits high affinity to the TSC complex, which is in a similar range as that between TSC1 and TSC2 (Dibble et al., 2012). We conclude that the TSC complex and G3BP1 form a highly stable complex that requires electrostatic interactions.

### G3BP1 bridges TSC2 to LAMP1/2

We next assessed the proximity of the G3BP1 association with TSC2, the LAMP1/2 proteins, and MTOR. Bimolecular fluorescence complementation (BiFC) assays detect protein-protein interactions in living cells with a maximum distance of 10 nm (Hu et al., 2002) (**Figure 4E, F** and **S3A**), and are thus indicative of close, likely direct contact between proteins. While all BiFC fusion proteins were expressed (**Figure S3B**), no BiFC signal was observed for cells in which G3BP1 was co-expressed with MTOR (**Figure 4E, F**). Thus, their interaction detected in IPs may not be direct, but is possibly mediated by their common association with lysosomes. In contrast, we did detect BiFC signals for G3BP1 with LAMP1, LAMP2, and TSC2, indicative of a close interaction between them. Based on this, and on our findings that G3BP1 knockdown impedes TSC2-LAMP1/2 binding (**Figure 2K-N and 3A, B**) and TSC2 KO prevents G3BP1 binding to TSC1-TBC1D7 (**Figure 4A**), we propose that G3BP1 bridges TSC2 to the lysosomal proteins LAMP1 and LAMP2, thereby mediating the lysosomal localization of the TSC complex.

### G3BP1 co-appears with the TSC complex during evolution

As our analyses established G3BP1 as a key component of mammalian TSC-mTORC1 signaling, we asked whether G3BP1 appeared during evolution together with the other subunits of the TSC complex and its targets. Therefore, we analyzed the phylogenetic distribution of G3BP1, TSC1, TSC2, TBC1D7, RHEB, and MTOR (**Figure 4G**). While MTOR and RHEB are present in the yeast *S. cerevisiae,* G3BP1 appears together with the other TSC complex components in the clade of Deuterostomia. Although G3BP1 orthologues have been proposed in *S. cerevisiae* (Yang et al., 2014) and in the nematode *C. elegans* (Jedrusik-Bode et al., 2013), evidence for their functional homology with G3BP1 is scarce. Our sequence similarity analyses (BLASTP, NCBI NR database, BLOSUM45 matrix; 19.02.2020) showed that the human protein with the highest similarity to the proposed G3BP1 orthologue Bre5 (UniProt ID P53741) in *S. cerevisiae* is a *C. elegans* UNC-80 like protein that is functionally unrelated to G3BP1. And although the *C. elegans* protein GTBP-1 (UniProt ID Q21351) exhibits the highest sequence similarities to human G3BP1 and 2, the similarities are low (evalues 4-e7 and 0.12) and are restricted to the NTF2 and RRM domains of which they cover only 23%, thus not matching the thresholds for our phylogenetic analysis. In summary, while SG existed already in low eukaryotes, including *S. cerevisiae* (Hoyle et al., 2007), we propose that a functional G3BP1 orthologue emerged later together with the TSC complex.

### G3BP2 is a functional paralogue of G3BP1 in mTORC1 signaling

G3BP2 exhibits high identity and similarity with G3BP1 (**Figure S4A, B**) (Kennedy et al., 2001), and can substitute for G3BP1 in SG assembly (Kedersha et al., 2016; Matsuki et al., 2013). Thus, G3BP1 and 2 might be redundant for many functions, and we asked whether G3BP2 might also compensate for G3BP1 in mTORC1 signaling. Indeed, phylogenetic analysis suggests that G3BP2 emerged together with G3BP1 indicating that they both evolved from a common ancestor gene as functional components of the TSC-mTORC1 axis (**Figure 5A**). Like G3BP1, G3BP2 co-immunoprecipitated with the TSC complex and MTOR (**Figure 5B, Figure S4C**). G3BP2 co-fractionated with G3BP1 and lysosomal proteins in sucrose gradients, identifying the lysosome as their primary localization site when SG are absent (**Figure 5C**). G3BP2 gave rise to BiFC signals with LAMP1, LAMP2, and TSC2 (**Figure 5D, E and S4D, E**), suggesting that G3BP2 binds to TSC2 and the LAMP1/2 proteins directly. G3BP2 knockdown enhanced RPS6KB1-T389 and RPS6-S235/236 phosphorylation, indicative of mTORC1 hyperactivity (**Figure 5F-I**). In agreement with previous data (Kedersha et al., 2016), G3BP2 expression was enhanced in G3BP1 KO cells (**Figure 5J, K**) and less so upon G3BP1 knockdown (**Figure 5L, M**). This suggests that indeed G3BP2 induction may partially compensate for G3BP1 KO, highlighting the strength of the effect of G3BP1 on mTORC1 activity (**Figure 1N-Q**). Thus, we conclude that G3BP2 is a functional paralogue of G3BP1 in TSC-mTORC1 signaling.

**Figure 5.**
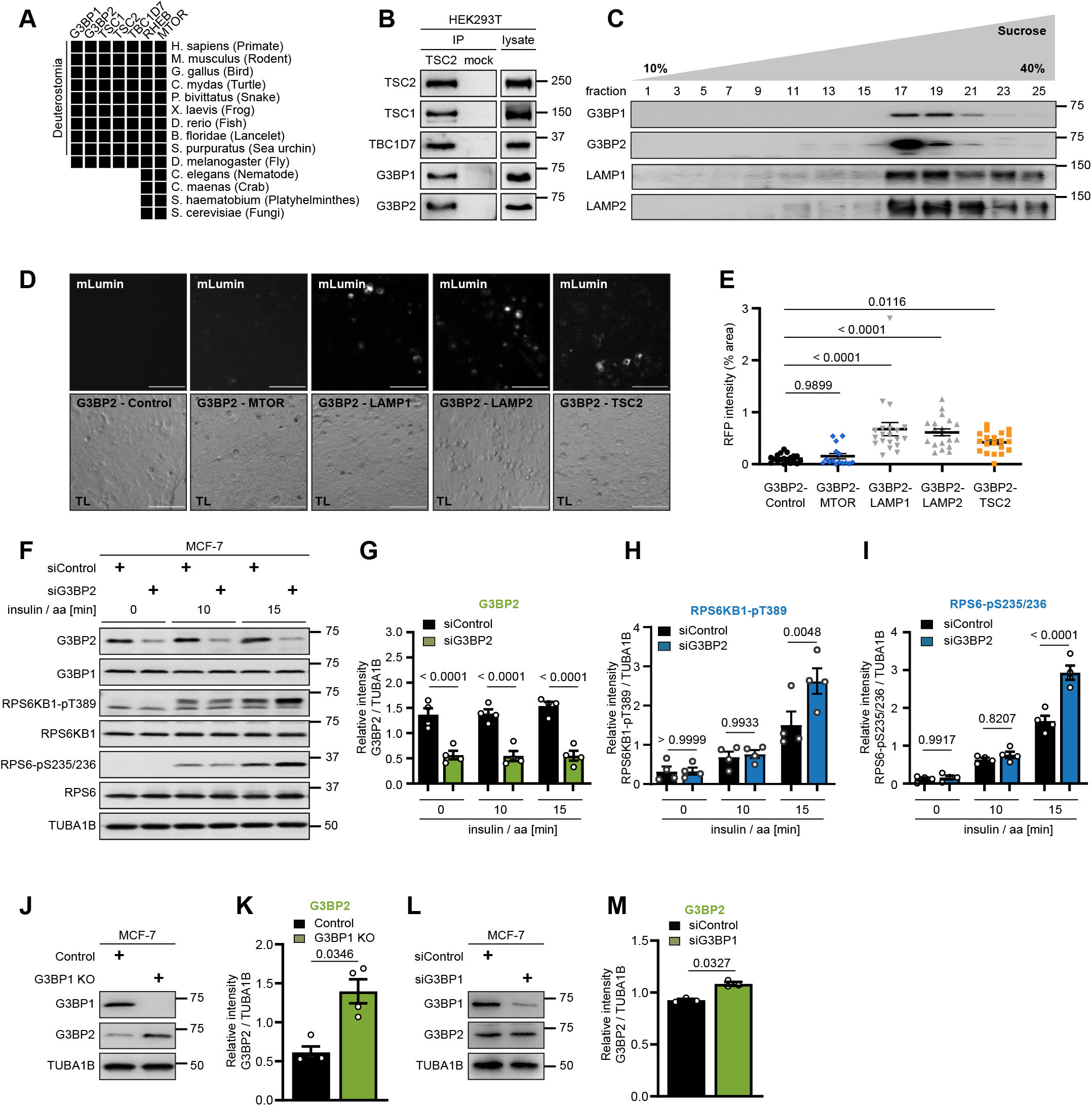
G3BP2 shares the function of G3BP1 in the TSC-mTORC1 axis. **(A)** Reanalysis of phylogenetic Blast analysis presented in **Figure 4G** including G3BP2 in addition. **(B)** IPs from HEK293T cells with antibodies against TSC2 (TSC2 #1) or mock (mouse IgG). Data shown are representative of n = 3 biological replicates. **(C)** Separation of MCF-7 cell lysates by a 10 to 40% sucrose density gradient. Cells were serum and amino acid starved. Data shown are representative of n = 3 biological replicates. **(D)** BiFC analysis of HEK293T cells transfected with plasmids carrying G3BP2 fused to a C-terminal mLumin fragment, together with an N-terminal mLumin fragment only (Control), or an N-terminal mLumin fragment fused to MTOR, LAMP1, LAMP2 or TSC2. Scale bar 100 μm. One representative image of each channel is shown for n = 4 biological replicates. A scheme depicting the fusion constructs is shown in **Figure S4D**. **(E)** Quantitation of data shown in (**D**). Data are shown as the mean ± SEM and overlaid with the single data points represented as dot plots. The percentages of mLumin fluorescence intensity (RFP)/ picture were compared between G3BP2-Control and the different plasmid combinations (G3BP2-MTOR, G3BP2-LAMP1, G3BP2-LAMP2, G3BP2-TSC2), using a oneway ANOVA followed by a Sidak’s multiple comparisons test across at least 15 biological fields of view from n=4 biological replicates. p-values are presented above the corresponding bar graphs. **(F)** MCF-7 cells transfected with siControl or siG3BP2 were serum and amino acid starved, and stimulated with 100 nM insulin / aa for the indicated time periods. Data shown are representative of n = 4 biological replicates. **(G)** Quantitation of G3BP2 immunoblot data shown in (**F**). Data are shown as the mean ± SEM and overlaid with the single data points represented as dot plots. G3BP2 levels (black and green bars) were compared between siControl and siG3BP2 cells, using a one-way ANOVA followed by a Sidak’s multiple comparisons test across n = 4 biological replicates. p-values are presented above the corresponding bar graphs. **(H)** Quantitation of RPS6KB1-pT389 immunoblot data shown in (**F**). RPS6KB1-pT389 levels (black and blue bars) are represented and compared between siControl and siG3BP2 cells as described in (**G**). **(I)** Quantitation of RPS6-pS235/236 immunoblot data shown in (**F**). RPS6-pS235/236 levels (black and blue bars) are represented and compared between siControl and siG3BP2 cells as described in (**G**). **(J)** G3BP1 CRISPR/Cas9 KO or control MCF-7 cells were serum and amino acid starved. Data shown are representative of n = 4 biological replicates. **(K)** Quantitation of data shown in (**J**). Data are shown as the mean ± SEM and overlaid with the single data points represented as dot plots. G3BP2 levels (black and green bars) were compared between control and G3BP1 KO cells, using a paired two-tailed Student’s t-test across n = 4 biological replicates. The p-value is presented above the bar graph. **(L)** MCF-7 cells, transfected with siControl or siG3BP1 were serum and amino acid starved. Data shown are representative of n = 3 biological replicates. **(M)** Quantitation of data shown in (L). Data are shown as the mean ± SEM and overlaid with the single data points represented as dot plots. G3BP2 levels (black and green bars) were compared between siControl and siG3BP1 cells, using a paired two-tailed Student’s t-test across n = 3 biological replicates. The p-value is presented above the bar graph.

### G3BP1 suppresses mTORC1-driven migration in breast cancer cells

We next investigated the consequences of G3BP1-mediated mTORC1 suppression in the context of cancer. In migration assays, G3BP1 deficiency resulted in faster wound closure, which was abrogated by rapamycin (**Figure 6A, B**). This suggests that G3BP1 restricts mTORC1-driven cell motility. As changes in proliferation might confound cell motility assays, we analyzed proliferation by real-time cell analysis (RTCA). In line with previous findings (Winslow et al., 2013), G3BP1-deficiency reduced cell proliferation (**Figure 6C, D**), indicating that mTORC1-driven cell motility in G3BP1-deficient cells was not a result of enhanced proliferation. Analysis of RNASeq data from invasive breast cancer revealed G3BP1 mRNA expression levels to be similar in the four breast cancer subtypes defined by the PAM50 classification (Koboldt et al., 2012) (**Figure 6E**). Analysis across all subtypes showed that patients with *G3BP1* mRNA or protein expression below the median exhibited significantly shorter relapse free survival (RFS) than those with expression above the median (**Figure 6F, G**). Our observations phenocopied the shorter RFS in patients with low TSC1 or TSC2 levels (**Figure 6H, I**). This suggests that G3BP1 and the two core TSC complex components could be used as subtype-independent prognostic markers in breast cancer patients and indicators of mTORC1 activity and cancer cell motility.

**Figure 6.**
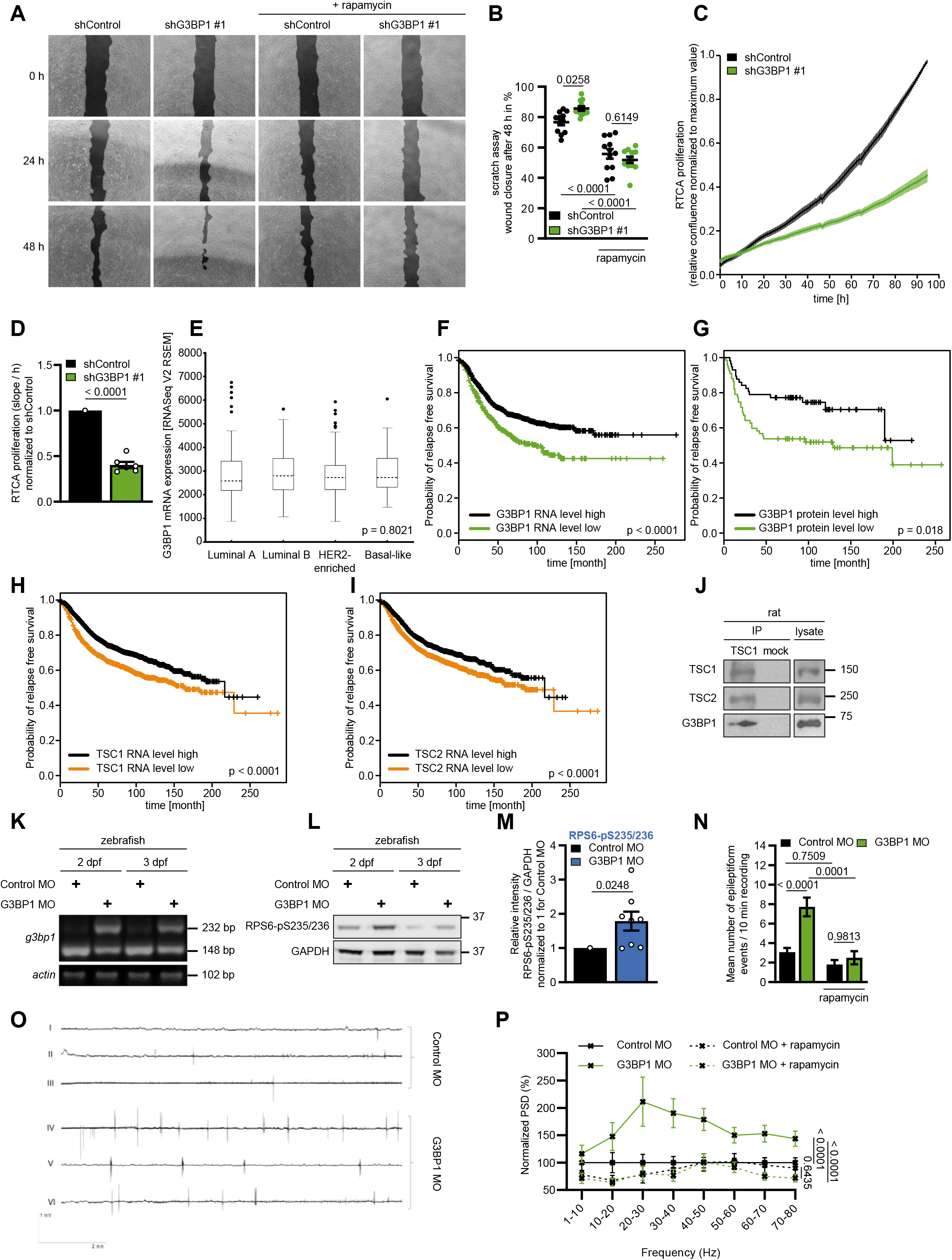
G3BP1 inhibits mTORC1-driven cancer cell motility and epileptogenic events. **(A)** Scratch assay in shG3BP1 #1 or shControl MCF-7 cultures. Pictures were taken at 0, 24, and 48 hours. Rapamycin was added 24 hours prior to the 0 h time point. The scratch was highlighted using the TScratch software (Geback et al., 2009). A representative image for each condition is shown. Data shown are representative of n = 3 biological replicates. **(B)** Quantitation of data shown in (**A**). Data are shown as the mean ± SEM and overlaid with the single data points represented as dot plots. Percentage of wound closure at 48 h was normalized to the initial wound area (0 h), and compared between shControl and shG3BP1 #1 cells, using a one-way ANOVA followed by a Sidak’s multiple comparisons test across n = 12 scratches from n = 3 biological replicates. p-values are presented above or below the corresponding bar graphs. **(C)** RTCA proliferation analysis of shG3BP1 #1 or shControl MCF-7 cells. The impedance was measured every 30 minutes for 5 days. Displayed is the relative confluence of cells normalized to 1 for the maximum value. Data are shown as the mean ± SEM for n = 6 biological replicates. **(D)** Quantitation of data shown in (**C**). The proliferation (slope/ hour) was compared between shControl and shG3BP1 #1 cells using a paired two-tailed Student’s t-test across n = 6 biological replicates. Data were normalized to the shControl condition, which was set to 1. Data are shown as the mean ± SEM with the corresponding dot plots overlaid. p-values are presented above the corresponding bar graphs. **(E)** *G3BP1* mRNA expression analysis. RNA seq V2 RSEM values from TCGA invasive breast cancer (TCGA, provisional) were classified according to PAM50 and analysed regarding *G3BP1* mRNA expression. Expression of *G3BP1* in luminal A (n = 231), luminal B (n = 127), HER2-enriched (n = 58) and basal-like (n = 97) breast cancer samples was analysed using a Kruskal-Wallis ANOVA by ranks. Data are shown as boxplots, representing the median with 25th and 75th percentiles as boxes and 5th and 95th percentiles as whiskers. The p-value of the Kruskal-Wallis ANOVA by ranks is shown. **(F)** Relapse-free survival of breast cancer patients based on *G3BP1* mRNA expression (probeID: 225007_at). Patients with high G3BP1 mRNA expression (n=1224) were compared to patients with low expression (n=409). Breast cancer patients were divided based on the best performing threshold. The survival period was assessed using the log-rank test and the p-value is presented. **(G)** Relapse-free survival comparing patients with high G3BP1 protein levels (n=57, probeID: Q13283) to those with low (n=67) G3BP1 protein expression. Breast cancer patients were divided based on the best performing threshold. The survival period was assessed using the log-rank test and the p-value is presented. **(H)** Relapse-free survival of breast cancer patients based on *TSC1* mRNA expression (probeID:209390_at). Patients were split into those with high expression (n= 2541) and low expression levels (n=1030). Breast cancer patients were divided based on the best performing threshold. The survival period was assessed using the log-rank test and the p-value is presented. **(I)** Relapse-free survival of breast cancer patients based on *TSC2* mRNA expression (probeID: 215735_s_a). Patients were split into those with high expression (n=1712) and low expression levels (n= 1859). Breast cancer patients were divided based on the best performing threshold. The survival period was assessed using the log-rank test and the p-value is presented. **(J)** IPs from brain tissue of rats with antibodies against TSC1 (TSC1 #3) or mock (rabbit IgG). Data shown are representative of n = 2 biological replicates. **(K)** PCR of control (control MO) and G3BP1 (G3BP1 MO) morpholino-injected zebrafish larvae at 2 and 3 days post fertilization (dpf). 10 larvae per condition were pooled. Data shown are representative of n = 3 biological replicates. **(L)** Zebrafish larvae, injected with control MO or G3BP1 MO for 2 or 3 days were analyzed by immunoblot. Data shown are representative of n = 4 biological replicates. **(M)** Quantitation of RPS6-pS235/236 immunoblot data shown in (**L**). Data are shown as the mean ± SEM and overlaid with the single data points represented as dot plots. Protein levels were normalized to the loading control GAPDH and then to the intensity of the control MO. The normalized RPS6-pS235/236 values were pooled for day 2 and 3. Control and G3BP1 MO (black and blue bars) were compared using a paired two-tailed Student’s t-test across n = 4 biological replicates. The p-value is presented above the bar graph. **(N)** Control and G3BP1 MO injected zebrafish larvae were treated on 3 dpf for 24 h with rapamycin or left untreated. Non-invasive local field potentials were recorded for 10 minutes from larval optic tecta at 4 dpf. Epileptiform events are represented as the mean ± SEM, and were compared between control and G3BP1 MO using a one-way ANOVA followed by a Sidak’s multiple comparisons test across 20 larvae per condition. p-values are presented above the corresponding bar graphs. **(O)** Non-invasive local field potentials in control and G3BP1 MO (quantified and described in **(N)**). Three representative 10 minutes recordings are shown for control and G3BP1 MO. **(P)** Power spectral density (PSD) estimation for data shown in **(N)**. Data are represented as mean ± SEM. The PSD was compared, using a two-way ANOVA followed by a Sidak’s multiple comparison test across 20 larvae per condition. p-values are presented for the comparisons between control MO versus G3BP1 MO, G3BP1 MO versus G3BP1 MO + rapamycin, and control MO + rapamycin versus G3BP1 MO + rapamycin.

### Brain G3BP1 suppresses mTORC1-driven epileptogenic events

Next to its importance as a tumor suppressor, the TSC complex has crucial neuronal functions and epilepsy is a hallmark of TSC. Therefore, G3BP1 may play a similar role in the brain. To test this, we conducted TSC1 IPs from rat brain lysates (**Figure 6J**). Together with TSC2, G3BP1 co-immunoprecipitated with TSC1, indicating that G3BP1 binds the TSC complex in the brain. To explore the impact of G3BP1 in epilepsy, we used a zebrafish model in which *tsc2* KO elicits pronounced epileptiform events and which is thus suitable to recapitulate the human TSC disease (Scheldeman et al., 2017). The zebrafish G3BP1 orthologue exhibits 67.8% sequence identity with the human protein (**Figure S5A**). We targeted zebrafish *g3bp1* with morpholino oligonucleotides (G3BP1 MO) (**Figure S5B**). Efficient *g3bp1* knockdown was evaluated by RT-PCR (**Figure 6K**). In agreement with our observations in human cell lines, *g3bp1* inhibition enhanced mTORC1 activity, as determined by RPS6-pS235/236 levels, in the zebrafish larvae (**Figure 6L, M**). Recordings of non-invasive local field potentials (LFP) from larval optic tecta (**Figure 6N, O** and **S5C, D**) revealed that *g3bp1* deficiency elicits epileptiform events. We tested whether the increased number of epileptiform events was due to hyperactive mTORC1. To reduce mTORC1 hyperactivity, we treated control and G3BP1 MO injected larvae with rapamycin prior to brain activity recordings. Rapamycin fully suppressed the epileptiform events in *g3bp1*-deficient larvae to the level in control animals (**Figure 6N**). We confirmed this result by power spectral density (PSD) analysis (**Figure 6P**), an automated method to quantify the spectral power across multiple LFP recordings (Hunyadi et al., 2017). We found that *g3bp1* deficiency enhanced the LFP power in the frequency range between 20-80 Hz, an effect that was fully rescued by rapamycin (**Figure 6P**). Taken together, we conclude that *g3bp1* deficiency elicits mTORC1-driven epileptiform events. Thus, *g3bp1* inhibition phenocopies the effect of a *tsc2* KO (Scheldeman et al., 2017), highlighting the importance of *g3bp1* as a suppressor of neuronal mTORC1 *in vivo.*

## Discussion

In this study, we demonstrate that G3BP1 acts outside of SG as a lysosomal tether of the TSC complex (**Graphical Abstract**). G3BP1 directly interacts with TSC2 and LAMP1/2, thus securing the TSC complex to lysosomes. Similar to the known TSC complex subunits, G3BP1 suppresses mTORC1. TSC2 and G3BP1 do not exert additive effects on mTORC1 activity in insulin-stimulated cells, highlighting that they act together in the insulin-mTORC1 axis. G3BP1 deficiency leads to mTORC1-driven phenotypes in both cancer and neuronal dysfunction. Thus, we propose that G3BP1 is not only a core SG component but also a key element of mTORC1 signaling on lysosomes.

G3BP1 was identified over two decades ago as a RasGAP binding protein, and thus a role of G3BP1 in the RAS pathway was proposed (Gallouzi et al., 1998; Kennedy et al., 2001; Parker et al., 1996). However, this hypothesis has been questioned (Annibaldi et al., 2011) and present research primarily focuses on the role of G3BP1 in SG formation and RNA metabolism (Alam and Kennedy, 2019; Reineke and Neilson, 2019). In line with the initial reports, we demonstrate that G3BP1’s identification as a GAP-binding protein was correct – although for a different GAP – as it exerts this role by binding TSC2, the GAP component of the TSC complex (Inoki et al., 2003). It therefore may be rewarding to revisit whether G3BP1 also binds to other RAS-related GAPs. Our data indicate that, at least in the insulin-mTORC1 axis, G3BP1 exerts its suppressor function through the TSC complex. However, this does not exclude involvement in other signaling pathways such as RAS (Parker et al., 1996), NFKB1 (Prigent et al., 2000), WNT (Bikkavilli and Malbon, 2011), and TGFB (Zhang et al., 2015). As they all crosstalk with mTORC1 via the TSC complex (Ghosh et al., 2006; Inoki et al., 2006; Ma et al., 2005; Thien et al., 2015), the observations implicating G3BP1 in these pathways might in fact result from its function within the TSC complex; which will be an intriguing direction for future research.

Why does G3BP1 inhibit mTORC1 upon metabolic starvation and restimulation, but not under stress conditions that promote SG formation? It is well documented that arsenite and other SG-inducing stressors enhance TSC2 degradation (Heberle et al., 2019; Huang and Manning, 2008; Orlova and Crino, 2010; Thedieck et al., 2013). Without TSC2, G3BP1 cannot bind to the TSC complex (**Figure 4A**) and thus cannot inhibit mTORC1. Another mechanism by which G3BP1 might inhibit mTORC1 under stress is through its role as a nucleator of SG, which restrict mTORC1 activity (Thedieck et al., 2013; Wippich et al., 2013). However, previous studies (Bley et al., 2015; Kedersha et al., 2016; Matsuki et al., 2013) and our own results (**Figure S2N, O**) show that SG are present in G3BP1-deficient cells. SG formation in the absence of G3BP1 is mediated by other SG factors such as T cell internal antigen 1 (TIA1) (Kedersha et al., 2016) or the G3BP1-paralogue G3BP2 (Kedersha et al., 2016; Kennedy et al., 2001; Matsuki et al., 2013), and thus SG remain to inhibit mTORC1. Hence, the absence of G3BP1’s inhibitory effect on mTORC1 in arsenite-stressed cells is likely due to (i) the degradation of TSC2 and (ii) the presence of SG in the absence of G3BP1.

By means of biochemical approaches, we identify the lysosome as the primary site of G3BP1 localization when SG are absent (**Figure 2A** and **5C**). This is in agreement with the major function of the TSC complex and mTORC1 at lysosomes, and this view is further supported by the appearance of G3BP1 in a recently published study on the lysosomal proteome (Wyant et al., 2018). Interestingly, SG have recently also been reported to hitchhike on lysosomes with annexin A11 (ANXA11) acting as a tether (Liao et al., 2019). The proximity of SG to lysosomes might allow G3BP1 shuttling, enabling rapid switching between its two functions. Despite the strong biochemical evidence for its lysosomal localization, we do not exclude that G3BP1 controls signaling at other subcellular sites. IF data show a ubiquitous cytoplasmic distribution of G3BP1 in the absence of SG (**Figure S2N**) (Irvine et al., 2004), reminiscent of the IF patterns for the TSC complex (Carroll et al., 2016; Demetriades et al., 2014) and MTOR (Betz and Hall, 2013). Indeed, next to lysosomes, MTOR has been proposed to localize to multiple subcellular sites (Betz and Hall, 2013), and accumulating evidence suggests that both RHEB and the TSC complex can reside at sites other than lysosomes (Hao et al., 2018; Zhang et al., 2013). Thus, both biochemical data and imaging results correlate with our suggestion of a functional connection between G3BP1, the TSC complex and mTORC1 at lysosomes and, likely, other subcellular loci (Kim and Guan, 2019).

The proposed function of G3BP1 and the TSC complex in the same pathway would suggest that deficiency of either factor affects mTORC1-driven phenotypes in a similar way. Ablation of the *TSC1* or *TSC2* tumor suppressor genes results in increased cancer cell motility and metastasis (Astrinidis et al., 2002; Goncharova et al., 2006). Similarly, G3BP1 deficiency enhances cancer cell motility in an mTORC1-dependent manner (**Figure 6A, B**), and low G3BP1 mRNA and protein levels correlate with a poor outcome in breast cancer (**Figure 6F, G**). Conflicting observations on the effect of G3BP1 on cell motility (Alam and Kennedy, 2019) may arise from the growth defect, which we (**Figure 6C, D**) and others observe upon G3BP1 inhibition (Alam and Kennedy, 2019; Dou et al., 2016; Huang et al., 2016; Wang et al., 2018). This growth defect has been attributed to the de-repression of cell cycle arrest factors whose mRNAs are bound and inhibited by G3BP1 (Alam and Kennedy, 2019). Such cell cycle defects can mask G3BP1’s inhibitory effect on migration, depending on the cell context and type of assay.

The opposite effects of G3BP1 on migration and proliferation may also limit its potential as a therapeutic target in cancer. In addition, the dual roles of G3BP1 in oncogenic mTORC1 signaling versus SG formation argue against G3BP1 as an anti-tumor target, as G3BP1 inhibition alone is not sufficient to inhibit SG (**Figure S2N, O**; and (Kedersha et al., 2016)), but results in mTORC1 hyperactivation. G3BP1 may, however, be a promising marker to guide drug therapies targeting mTORC1 and its upstream cues. Such compounds have been approved for several tumor entities including metastatic ER-positive breast cancer (Baselga et al., 2012; Paplomata and O’Regan, 2014), but their clinical success so far remained limited (Friend and Royce, 2016). At first glance, our finding that low G3BP1 levels correlate with a shorter progression-free survival in breast cancer seems at odds with reports on sarcoma (Somasekharan et al., 2015), colon (Zhang et al., 2012), and gastric cancer (Min et al., 2015), in which high G3BP1 expression positively correlates with tumor size, invasion, and metastasis. Yet, SG were found to be critical for G3BP1-mediated oncogenicity in these entities, suggesting that the function of G3BP1 as a SG nucleator may dominate in these cases. This effect likely is less important in tumors addicted to hyperactive mTORC1, in which G3BP1 may act as a tumor suppressor. This suggests that G3BP1 is a poor prognostic marker across different cancer entities as both high and low levels can be oncogenic. However, low G3BP1 levels are likely a good indicator of mTORC1 hyperactivity, which correlates with tumor sensitivity to mTORC1 inhibitors (Grabiner et al., 2014; Kwiatkowski and Wagle, 2015; Meric-Bernstam et al., 2012; Wagle et al., 2014). Therefore, low G3BP1 levels might enable the stratification of patients to clinical inhibitors of mTORC1 and its upstream cues.

Also neuronal G3BP1 phenotypes deserve evaluation as to whether they are mediated by the TSC-mTORC1 axis. G3BP1 deficiency impairs synaptic transmission (Martin et al., 2013; Zekri et al., 2005) and there is evidence for a linkage with early-onset epilepsy in humans (Appenzeller et al., 2014; Heyne et al., 2018). Our finding that *g3bp1* inhibition elicits epileptogenic events in zebrafish (**Figure 6N, O**) supports a link between G3BP1 deficiency and epilepsy. G3BP1 down-regulation inactivates the TSC complex, and *TSC1* and *TSC2* mutations – leading to de-repression of mTORC1 – frequently cause epilepsy (Curatolo et al., 2015; Jozwiak et al., 2019; Roach and Kwiatkowski, 2016). Consistent with a common mechanism, rapamycin suppresses the epileptogenic events in *g3bp1* deficient zebrafish larvae (**Figure 6N**). G3BP1’s function via the TSC complex, the insulin responsive GAP of RHEB, is mirrored by the KICSTOR complex (Peng et al., 2017; Wolfson et al., 2017). The KICSTOR complex is the lysosomal tether for the GATOR1 subcomplex, which is the GAP for the RAG GTPases that activate mTORC1 in response to amino acids. Like mutations in the genes encoding the components of the TSC complex, mutations in genes encoding the KICSTOR complex (Wolfson et al., 2017) and GATOR1 subcomplex (Baldassari et al., 2016) components have been associated with neuronal malformation and epilepsy, referred to as “mTORopathies” (Crino, 2015; Wong and Crino, 2012). mTORC1 inhibitors show encouraging results for the treatment of TSC-related epilepsy (van der Poest Clement et al., 2020) and have been proposed to benefit epilepsy patients with alterations in KICSTOR or GATOR1 (Baulac, 2016; Crino, 2015; Sadowski et al., 2015). Our findings suggest that also epilepsy patients with G3BP1 alterations may benefit from treatment with mTORC1 inhibitors, which will add G3BP1 to the family of genes whose mutations cause mTORopathies.

In conclusion, we identify G3BP1 as an essential lysosomal tether of the TSC complex that suppresses mTORC1 at lysosomes. Future research will reveal whether this dual role in nutrient signaling and SG formation is specific to G3BP1, or whether also other SG components have non-granule functions to orchestrate cellular responses to environmental signals.

## Supporting information

Supplemental Figures and Key Resource Table

## Acknowledgements

We thank Manuela Brom and Felix Bestvater from the Light Microscopy Facility (German Cancer Research Center, Heidelberg) for excellent wide-field microscopy resources and their support in image acquisition, and Damir Krunic, Fabian Tetzlaff and Gergely M. Solecki for their help with ImageJ (ImageJ version 1.50b and Fiji version 1.49v). We thank Ursula Klingmüller and Jochen Utikal for kindly providing access to their camera facilities. We thank the FACS & Imaging Core Facility at the Max Planck Institute for Biology of Ageing for support. We thank Jan Maes for help with zebrafish microinjections. We thank Michael N. Hall (Biozentrum, University of Basel, Switzerland) for kindly sharing the polyclonal TSC1 and TSC2 antibodies (Molle, 2006). R777-E138 Hs.MTOR-nostop and R777-E356 Hs.TSC2-nostop were gifts from Dominic Esposito (Addgene plasmids # 70422 and # 70640). The BiFC plasmids bFos-myc-LC151 and bJun-HA-LN-151 were gifts from Qingming Luo (Huazhong University of Science and Technology (HUST), Wuhan, China). We thank Dyah L. Dewi, Ahmed Sadik, Luis F. Somarribas Patterson, Laura Corbett, Kathrin Breuker, and José Ramos Pittol for support and helpful discussions.

M.T.P. is recipient of the Research Award of the German Tuberous Sclerosis Foundation 2019 and was supported by the German Research Foundation (Excellence Initiative GSC-4, Spemann Graduate School). M.C.S. was supported by the Graduate School of Medical Sciences of the University of Groningen. K.T. is recipient of the Research Award of the German Tuberous Sclerosis Foundation 2017 and acknowledges support from the German TS Foundation and the Stichting TSC Fonds. K.T. acknowledges funding from the BMBF e:Med initiative MAPTor-NET (031A426B), a Rosalind-Franklin-Fellowship of the University of Groningen, the Ubbo Emmius Fund, the German Research Foundation (DFG, TH 1358/3-1), the PoLiMeR Innovative Training Network (Marie Skłodowska-Curie grant agreement No. 812616) which has received funding from the European Union’s Horizon 2020 research and innovation programme. C.A.O., K.T. and S.T. were supported by the BMBF e:Med initiative GlioPATH (01ZX1402). C.A.O., I.H., and K.T. acknowledge support from the MESI-STRAT project, which has received funding from the European Union’s Horizon 2020 research and innovation programme under grant agreement No. 754688. T.Y. and L.A.H were supported by the Austrian Science Fund (FWF DK W11 and P26682_B21) and the Molecular Cell Biology and Oncology PhD program (MCBO) at the Medical University of Innsbruck. R.B. is supported by the Deutsche Forschungsgemeinschaft DFG (German Research Foundation under Germany’s Excellence Strategy EXC 294 and EXC-2189-Projektnummer 390939984; CRC850 and CRC1381). A.S.M. is a PhD fellow of the Fund for O6260 Research Foundation-Flanders (FWO, 11F2919N). B.C. is funded by the British Skin Foundation and Vice-Chancellor’s Fellowship, University of Bristol. C.D. is funded by the European Research Council (ERC) under the European Union’s Horizon 2020 research and innovation programme (grant agreement No 757729), and by the Max Planck Society. G.F. is recipient of a Long Term EMBO Postdoctoral fellowship. M.N. was supported by the Stichting Michelle, TS Alliance and TS Association (UK). J.J. and A.K. are financed by the TEAM grant from the Foundation for Polish Science (POIR.04.04.00-00-5CBE/17-00).

## Author contributions

M.T.P., U.R. and M.C.S. planned and conducted experiments, analyzed the results, and wrote the manuscript. B.B., B.H., K.K., I.L.K., M.R., A.R., and F.R. supported the experimental work. S.P. performed cloning and supported BiFC analysis. A.v.D. enabled cloning and BiFC analysis. A.S.M., C.S., and P.W. conducted the zebrafish experiments. R.B. was involved in the initial phases of the project. L.B. and C.D. assisted with experiments, supported imaging analyses and data interpretation and provided strategic advice. M.B. and I.H. conducted the phylogenetic analyses. B.C. and V.K. conducted and analyzed IF experiments, and G.F. and A.T. conducted and analyzed PLAs to determine lysosomal localization of the TSC. A.M.H. supported the experimental work and manuscript writing. M.N., L.A.H., and T.Y. supported the generation of CRISPR cell lines. A.K. provided input on IDRs and charged residues in G3BP1. M.M. and J.J. conducted IPs from rat brain extracts. O.T.Q. and E.S. identified the G3BP1 region that interacts with TSC2. S.T. conducted the expression and survival analyses. C.A.O. and K.T. planned and guided the project and wrote the manuscript. All authors read and revised the manuscript. Apart from the first and last authors, all other author names are listed in alphabetical order.

## Star Methods

### Contact for Reagent and Resource Sharing

Further information and requests for resources and reagents should be directed to and will be fulfilled by the Lead Contact, Kathrin Thedieck.

### Method Details

#### Cell culture conditions and cell treatments

Experiments were performed in HeLa alpha Kyoto cells, MCF-7 cells (ACC115), MCF-7 cells expressing GFP-LC3 (MCF-7-LC3), MDA-MB-231, HEK293T, and HEK293-β_2_AR cells. All cells, except of HEK293-β_2_AR, were cultivated in Dulbecco’s modified Eagle’s medium (DMEM) with 4.5 g/L glucose, supplemented with 10% fetal bovine serum (FBS) and 3 mM L-glutamine (termed full DMEM medium) if not indicated otherwise. HEK293-β_2_AR were cultured in DMEM with 4.5 g/L glucose and 0.584 mM L-glutamine, supplemented with 10% FBS and 1% penicillin and streptomycin. All cell lines were maintained at 37°C in a 5% CO_2_ incubator and regularly tested for mycoplasma infection.

SG formation was induced with arsenite at a final concentration of 500 μM for the indicated time periods. Prior to arsenite stress, cells were washed with phosphate-buffered saline (PBS) and serum starved for 16 hours.

Metabolic stimulation experiments: for serum and amino acid starvation, cells were washed in PBS and cultured for 16 hours in Hank’s balanced salt solution (HBSS). For stimulation with insulin and amino acids (insulin / aa), the medium was exchanged to DMEM supplemented with 3 mM L-glutamine and 100 nM or 1 μM insulin, as indicated in the figure legends.

For serum starvation, cells were washed in PBS and cultured for 16 hours in DMEM with 4.5 g/L glucose, supplemented with 3 mM L-glutamine. For stimulation with insulin alone, insulin was directly added to the starvation media for the time periods indicated in the figure legends.

Lyophilized rapamycin was dissolved in methanol to a concentration of 1 nmol / μL and aliquoted to 5 μL per tube. 5 μL aliquots were dried with open lids under a sterile cell culture hood and deep frozen at −80° degrees. Aliquots were thawed immediately before an experiment and methanol-dried rapamycin was directly dissolved in HBSS or DMEM to a final concentration of 20 or 100 nM, as indicated. Hence, no carrier was used in experiments with rapamycin.

#### RNA knockdown experiments

siRNA knockdown of G3BP1, G3BP2 and RHEB was induced for two days using ON-TARGET plus SMARTpool siRNA at a final concentration of 40 nM. As a negative control, a nontargeting scrambled siRNA pool (siControl) was used at the same concentration. siRNA transfection was performed using Lipofectamine 3000 or RNAiMAX transfection reagents according to the manufacturer’s protocols. The medium containing the transfection mix was replaced 6 hours after transfection. For PLA analysis in **Figure 3A**, siRNA knockdown of G3BP1 was induced for five days using siGENOME SMARTpool siRNA at a final concentration of 15 nM. Here siRNA against *Renilla* luciferase (Control) was used as a control.

Doxycyclin-inducible shRNA knockdown cell lines for G3BP1 were generated using the pTRIPZ system using the Trans-Lentiviral shRNA Packaging Mix (Horizon Discovery). Viral particles were produced using shRNA constructs targeting G3BP1 (shG3BP1 #1 or shG3BP1 #2) or a non-targeting scrambled control sequence (shControl) according to the manufacturer’s protocol. MCF-7-LC3 and MDA-MB-231 cells were transduced in three rounds. The cells were incubated with the viral supernatant containing 8 μg/mL polybrene for 16 hours, followed by 6 hours of fresh full medium. Antibiotic selection was carried out 48 hours posttransduction with 2 μg/mL puromycin for 7 days. Expression of the shRNA was induced with 2 μg/mL doxycycline for 4 days. Monoclonal cell populations were obtained by limiting dilutions. Knockdown efficiency was tested at protein level by immunoblotting.

#### Knockout cell lines

CRISPR/Cas9 knockout cell lines for G3BP1 and TSC2 were generated using a two-vector system as previously described (Sanjana et al., 2014). First, doxycyclin-inducible Cas9 expressing MDA-MB-231 and MCF-7 cell lines were generated by lentiviral transduction using the pCW-Cas9-Blast vector (Addgene plasmid # 83481) and thereafter selected with 5 μg/mL blasticidin for 48 hours. Next, the Cas9 expressing cells were transduced with the lentiGuide-Puro vector (Addgene plasmid # 52963) containing either no sgRNA (control), or sgRNA targeting G3BP1 (G3BP1 KO) or TSC2 (TSC2 KO). These cells were selected with 2 μg/mL puromycin for 48 hours. Monoclonal cell populations were obtained by limiting dilutions. Cas9 expression was induced with 2 μg/mL doxycycline for 48 hours. Knockout efficiency was tested at protein level by immunoblotting.

#### Cloning

The coding sequences (CDS) of G3BP1, G3BP2, LAMP1 and LAMP2 were obtained from the clone repository of the DKFZ Genomics and Proteomics Core Facility (GPCF) as Gateway® compatible clones in pENTR221 or pENTR223. The CDS of MTOR and TSC2 were gifts from Dominic Esposito (Addgene plasmids # 70422 and # 70640) and obtained as Gateway® compatible clones in pDonor-255. After sequence verification, the CDS were cloned into the BiFC destination vectors pGW-MYC-LC151 for G3BP1 and G3BP2, and pGW-HA-LN151 for LAMP1, LAMP2, MTOR and TSC2 by Gateway®-specific LR-reaction following the manufacturer’s protocol (Invitrogen). Previously, the vectors bFos-MYC-LC151 and bJun-HA-LN151 (Chu et al., 2009) were adapted for Gateway cloning. MYC-LC151 and HA-LN151 PCR-fragments were generated and cloned into modified pDEST26 vectors resulting in pGW-MYC-LC151 and pGW-HA-LN151, as previously described (Weiler et al., 2014). Using the Gateway®-specific LR reaction, TSC2 was also cloned into pEGFP-C. Three G3BP1 truncation constructs in pGW-MYC-LC151 were generated with primers placed at the end or start positions of each construct, respectively: G3BP1_1-182_-MYC, G3BP1_183-332_-MYC and G3BP1_333-466_-MYC. AttB sites were added to the CDS by a two-step PCR. The first PCR was performed with hybrid primers, consisting of half of the AttB sites and the other half being gene specific. The second PCR was done with primers covering the complete AttB sites (see key resources table for more details).

#### Cell lysis and immunoblotting

For lysis, cells were washed with PBS and lysed with radio immunoprecipitation assay (RIPA) buffer (1% IGEPAL CA-630, 0.1% SDS, and 0.5% sodium deoxycholate in PBS) supplemented with Complete Protease Inhibitor Cocktail, Phosphatase Inhibitor Cocktail 2 and Cocktail 3. Protein concentration was measured using Protein Assay Dye Reagent Concentrate and adjusted to the lowest value. Cell lysates were mixed with sample buffer (10% glycerol, 1% beta-mercaptoethanol, 1.7% SDS, 62.5 mM TRIS base [pH 6.8], and bromophenol blue), and heated for 5 minutes at 95 °C. Cell lysates were then loaded on SDS polyacrylamide gel electrophoresis (PAGE) gels with a concentration of 8%, 10%, 12% or 14% polyacrylamide. Polyacrylamide gels were prepared consisting of two distinct layers: a stacking and a separation gel. For the lower separation gel, polyacrylamide was diluted with 375 mM TRIS base [pH 8.8] to the respective percentage. For the upper stacking gel, polyacrylamide was mixed with 0.125 M TRIS base [pH 6.8] to a final concentration of 13%. Electrophoresis was carried out with a Mini-PROTEAN Tetra Vertical Electrophoresis Cell system that was filled with electrophoresis buffer (0.2 M glycine, 25 mM TRIS base, and 0.1% SDS), and an applied voltage of 90 to 150 V. Subsequently, proteins were transferred to polyvinylidene difluoride (PVDF) membranes by using a Mini-PROTEAN Tetra Vertical Electrophoresis Cell system filled with blotting buffer (0.1 M glycine, 50 mM TRIS base, 0.01% SDS, [pH 8.3], and 10% methanol) and an applied voltage of 45 V for 2 hours. Afterwards, membranes were blocked in 5% bovine serum albumin (BSA) – TRIS-buffered saline tween (TBST) buffer (0.15 M NaCl, 60 mM TRIS base, 3 mM KCl, and 0.1% Tween-20, [pH 7.4]). Membranes were incubated overnight with primary antibodies at 4 °C, following the manufacturer’s instructions for the respective antibodies. The next day, membranes were washed in TBST buffer and incubated for at least one hour with the corresponding horseradish peroxidase (HRP) coupled secondary antibodies. For detection, Pierce ECL western blotting substrate or SuperSignal West FEMTO were used to detect chemiluminescence using a LAS-4000 camera system, a ChemiDoc XRS+ camera or a Fusion Fx camera. For graphical presentation, raw images taken with the LAS-4000 or Fusion camera were exported as RGB color TIFF files using ImageJ version 1.50b, and further processed with Adobe Photoshop version CS5.1. Raw images taken with a ChemiDoc XRS+ camera were processed with Image Lab version 5.2.1 and exported for publication as TIFF files with 600 dpi resolution.

#### Immunoprecipitation (IP)

For IP experiments, cells were washed three times in ice-cold PBS and then harvested in CHAPS based IP lysis buffer (40 mM HEPES, 120 mM NaCl, and 0.3% CHAPS, [pH 7.5]) supplemented with Complete Protease Inhibitor Cocktail, Phosphatase Inhibitor Cocktail 2 and Cocktail 3. The lysate volume was adjusted to 1 – 2.5 mL per 15 cm cell culture plate, depending on the cell density. The lysate was incubated under gentle agitation for 20 minutes at 4 °C, centrifuged for 3 minutes at 600 g at 4 °C, the pellet was discarded and the supernatant was transferred to fresh tubes. In case of multiple samples, the protein concentration was measured using Protein Assay Dye Reagent Concentrate and all samples were adjusted to the lowest value. The lysates were pre-incubated with 10 μL pre-washed Protein G covered Dynabeads per mL of lysate for 30 minutes at 4 °C under gentle agitation. A fraction of each lysate was mixed with 5 x sample buffer, referred to as ‘lysate’ input in the figure panels. For IP, the pre-cleaned lysates were subdivided, and specific antibodies or isotype control IgG antibodies (mock condition) were added using 7.5 μg antibody per mL of pre-cleaned lysate. Isotype control IgG antibodies (mock antibodies) were used in the same concentration as the protein-specific antibodies. After 30 minutes at 4 °C under gentle agitation, 37.5 μL prewashed Protein G covered Dynabeads / mL lysate were added, and the incubation was continued for 90 minutes at 4 °C under gentle agitation. Finally, beads were washed with CHAPS lysis buffer three times shortly and three times for 10 minutes at 4 °C under gentle agitation, and taken up in 1 x sample buffer. Samples were heated for 5 minutes at 95 °C and separated by SDS PAGE. For IP experiments with TSC2 and respective mock antibodies, the samples were heated for 10 minutes at 70 °C.

For TSC1-IPs with NaCl and SDS washes, the IP was performed as detailed above but with a CHAPS-based IP lysis buffer without NaCl (40 mM HEPES, and 0.3% CHAPS, [pH 7.5]). Before the final washing steps, the TSC1-IP was subdivided into six tubes. Each IP was washed with CHAPS-based lysis buffer supplemented with the indicated NaCl or SDS concentrations three times shortly and three times for 10 minutes at 4 °C under gentle agitation, and taken up in 1 x sample buffer. Samples were heated for 10 minutes at 70 °C and separated by SDS PAGE.

For GFP-IP experiments, 1.7×10^6^ HEK293-β_2_AR cells per dish were seeded on 10 cm dishes (2 dishes per condition). 24 hours after seeding, the cells were co-transfected with 2 μg TSC2-GFP (full length) and 1 μg G3BP1-myc constructs (full-length or truncated versions) using Transfectin (ratio 2:1) in FBS-free DMEM, following the manufacturer’s protocol. After 48 hours of transient overexpression, cells were washed once in ice-cold PBS and pooled into one tube per condition. Cells were centrifuged at 16000 g for 1 minute at room temperature and resuspended in 1 mL of CHAPS-based IP lysis buffer, supplemented with protease inhibitors (100 μM Leupeptin, 100 μM Aprotinin, 1 μg / mL Pepstatin A) and phosphatase inhibitors (1 mM Sodium orthovanadate, 1 mM Sodium pyrophosphate, 1 mM sodium fluoride). The cells were disrupted and the DNA was sheared through the repeated use of a syringe with a 21G x 0.80 mm needle. Afterwards, the lysate was incubated on ice for 15 minutes at 4 °C, centrifuged for 45 minutes at 16000 g at 4 °C, the pellet was discarded and the supernatant was transferred to fresh tubes. If the supernatants appeared viscous the DNA shearing was repeated. Otherwise, the lysates were pre-incubated with 12 μL Protein G sepharose beads per mL of lysate for 60 minutes at 4 °C under gentle agitation. A fraction of each lysate was mixed with 5 x sample buffer (25 mM Tris-HCl pH 6.8; 4% (w/v) SDS; 3% (w/v) DTT; 0.02% (v/v) bromophenol blue), referred to as ‘lysate’ in the figure panels. For IP, the pre-cleared lysates were subdivided, and 1 μg/mL of anti-GFP antibody or anti-Flag antibody were added. After 3 hours at 4 °C under gentle agitation, 12 μL Protein G sepharose beads per mL lysate were added, and the incubation was continued for 60 minutes at 4 °C under gentle agitation. Finally, beads were washed with CHAPS-based lysis buffer five times shortly and once for 5 minutes at 4 °C under gentle agitation. In-between the samples were centrifuged for 1 minute at 9600 g to remove the supernatant. Finally, the IP samples were dissolved in 30 μL 1 x sample buffer. Samples were heated for 5 minutes at 95 °C and separated by SDS PAGE.

The animals that were used to obtain brain tissue for IP of endogenous TSC1 were sacrificed according to protocol, which complied with European Community Council Directive 2010/63/EU. The cerebral cortex from one hemisphere of a rat brain was homogenized in 4 mL lysis buffer (40 mM Tris-HCl pH 7.5, 120 mM NaCl) containing 0.3 % CHAPS, supplemented with protease and phosphatase inhibitors, using a glass teflon homogenizer. The homogenate was diluted 1:1 with lysis buffer containing 0.1 % CHAPS and incubated under gentle agitation for 90 minutes at room temperature. The brain lysate was centrifuged at 1000 g, 4 °C for 10 minutes, the pellet was discarded and the supernatant was transferred to fresh tubes. A fraction of each lysate was mixed with 4 x sample buffer, referred to as ‘lysate’ input in the figure panels. 30 μL of Protein G covered Dynabeads were pre-conjugated in lysis buffer containing 0.1 % CHAPS with 4 μg of TSC1 antibody or isotype control rabbit IgG (mock condition) for 2 hours at 4 °C. For IP, the pre-conjugated beads were incubated with the lysate at 4 °C overnight under gentle agitation. Finally, beads were washed with lysis buffer containing 0.1 % CHAPS four times for 3 minutes at 4 °C under gentle agitation, and taken up in 1 x sample buffer. Samples were heated for 10 minutes at 95 °C and separated by SDS PAGE.

#### Sucrose gradients

Cells were lysed in homogenization buffer (50 mM Tris-HCl, pH 7.4, 250 mM sucrose, 25 mM KCl, 5 mM MgCl2, 3 mM imidazole), supplemented with Complete Protease Inhibitor Cocktail and Phosphatase Inhibitor Cocktail 2 and Cocktail 3 on a rocking platform for 30 minutes at 4 C. Subsequently, cells were scraped and centrifuged at 12,000 g for 10 minutes at 4°C. The pellet was discarded, the supernatant was transferred to a fresh tube and the absolute protein concentration was determined with Protein Assay Dye Reagent Concentrate by calculating a BSA adjustment curve ranging from 0.5 mg / mL to 7.5 mg / mL BSA. 1.5 mg protein was loaded on 4 mL of a continuous sucrose gradient (10% to 40% sucrose) and centrifuged 194,000 x g for 16 hours. Each sample was divided into 26 fractions and 5 x sample buffer was added to a final concentration of 1 x. Every second fraction was analyzed by immunoblot.

#### Immunofluorescence (IF)

In order to analyze SG assembly, cells were grown on coverslips and treated as indicated in the respective figure captions. Cells were washed with PBS and fixed with ice-cold methanol for 5 minutes on ice. After fixation, cells were washed three times with PBS, and permeabilized with 0.1% Triton X-100 in PBS for 60 seconds. Cells were washed with PBS and blocked with 3% FCS in PBS for 30 minutes at room temperature, and incubated with primary antibodies against G3BP1 and EIF3A at 4 °C overnight. The cells were washed three times with PBS and incubated with Alexa Fluor 568 and Alexa Fluor 488 labeled secondary antibodies and Hoechst 33342 at room temperature for 30 minutes in the dark. Afterwards, cells were washed three times with PBS and twice in deionized water. The cells were mounted with Mowiol 4-88, including DABCO (1,4-diazabicyclo[2.2.2]octane) and supplemented with 10% NPG (n-propyl-gallate). Cells were analyzed by fluorescence microscopy. Images were taken using a wide-field AxioObserver Z1 microscope equipped with an Apotome, a 63x / 1.4 oil objective, and an AxioCamMRm CCD camera. For each experimental setup, the magnification and exposure times were adjusted to the condition with the brightest signal, and the settings were retained throughout for all conditions. For presentation in figures, single layers of Z-stacks were exported as TIFF with no compression using Zen2012 blue edition software, and brightness and contrast were adjusted for better visibility.

The number of SG / cell was analyzed on unprocessed image raw files without any adjustment using Fiji software version 1.49v, creating maximum intensity projections of all Z-stacks. We used a background subtraction of 1, threshold adjustment with the intermodes function, and the ‘Analyze Particles’ function with a particle size from 0.2-infinity and a circularity from 0.5-1. SG were counted using the EIF3A channel. The number of SG / image was then normalized to the number of cells by counting the nuclei in the Hoechst channel and analyzed using a one-way ANOVA followed by a Sidak’s multiple comparisons test.

TSC2-LAMP1 co-staining was performed as described previously (Carroll et al., 2016). Briefly, cells were grown on coverslips and treated as indicated in the figure. The medium was removed and cells were fixed with 4% formaldehyde in PBS for 10 minutes at room temperature. After fixation, cells were permeabilized with 0.5 % Triton X-100 in PBS for 10 minutes at room temperature. Cells were blocked with 5 % normal goat serum in PBS and 0.05 % Tween-20 for 1 hour at room temperature, and incubated with primary antibodies against TSC2 and LAMP1 at 4 °C overnight. The following day, cells were washed and incubated with the appropriate secondary antibodies for 1 hour at room temperature. Afterwards, the cells were washed and coverslips were mounted on slides with Prolong Gold antifade reagent with 4’,6-Diamidin-2-phenylindol (DAPI).

Cells were analyzed by fluorescence microscopy. Z-stack images were taken using a Leica SP8 microscope, a 63x objective, 1.5x digital zoom and filters suitable for the used fluorophores. Identical settings were used to capture images across five to six separate fields (20 to 40 cells) of view. For presentation in figures, pictures were opened in Fiji (version 1.52p) and Z-stacks were projected (max). Channels were split and brightness and contrast were adjusted for better visibility. Afterwards channels were converted to RGB colour. Regions of interest (ROI) were selected and coordinates were copied to maintain the same ROI in the different channels. For single channel images, channels were pseudo-coloured grey, for merge images, the Alexa 488 channel was left green and the Alexa 555 channel was pseudocoloured magenta. All images were exported as TIFF with no compression.

For TSC2-LAMP1 co-staining, the Manders’ coefficient was calculated using the Coloc2 plugin of the ImageJ software (v1.51r). Prior to running the plug-in, a mask was made of the DAPI channel and subtracted from the other channels. A constant threshold was applied to all the images in the Z-stack, and for every image within each experiment and the Manders’ colocalization coefficient was calculated. Differences in the tested conditions were analyzed using a one-way ANOVA followed by a Sidak’s multiple comparisons test across n = 5-6 fields of view from one dataset representative of at least three independent experiments.

#### Bimolecular fluorescence complementation (BiFC)

For BiFC analysis, we made use of the red fluorophore mLumin (Chu et al., 2009; Weiler et al., 2014). 24 hours prior to transfection HEK293T cells were seeded in a 24 well plate at 100,000 cells / well in full medium. The cells were transiently transfected with Lipofectamine 3000 following the manufacturer’s protocol in the following combinations: pGW-MYC-LC151-G3BP1 (G3BP1 fused to a C-terminal mLumin fragment) with empty pGW-HA-LN151 as a negative control (a N-terminal mLumin fragment only), and pGW-MYC-LC151-G3BP1 with either pGW-HA-LN151-LAMP1, pGW-HA-LN151-LAMP2, pGW-HA-LN151-MTOR, or pGW-HA-LN151-TSC2, respectively (a N-terminal mLumin fragment fused to LAMP1, LAMP2, MTOR or TSC2, respectively) (**Figure S3A**). For G3BP2, cells were transfected with pGW-MYC-LC151-G3BP2 and either empty pGW-HA-LN151 as a negative control, or pGW-HA-LN151-LAMP1, pGW-HA-LN151-LAMP2, pGW-HA-LN151-MTOR, pGW-HA-LN151-TSC2, respectively (**Figure S4D**). In order to achieve equal expression of all plasmids, 3 times the amount of DNA was used for the MTOR, TSC2 and empty control plasmids in comparison to the G3BP1, G3BP2, LAMP1, and LAMP2 plasmids. Cells were analyzed 48 hours after transfection using a wide-field AxioObserver Z1, equipped with a 10x / 0.3 Plan-NEO objective, an AxioCamMRm CCD camera and an mPlum (64 HE) filter. mLumin fluorescence was analyzed with Fiji version 1.49 using a background subtraction of 50, threshold adjustment from 20-max, a Gaussian Blur filter of 1 and the ‘Analyze Particles’ function with a particle size from 20-infinity. The mLumin fluorophore signal was measured in percent / image and compared between the different combinations by a one-way ANOVA followed by a Sidak’s multiple comparisons test across at least 3 independent fields of view from at least three independent datasets, respectively. In total at least 22 independent fields of view for G3BP1 and 15 independent fields of view for G3BP2 were analyzed. All pictures were taken from regions with a comparable cell density.

#### Proximity Ligation Assay (PLA)

For TSC2-LAMP2 PLAs, 72 h after siRNA transfection, MCF-7 cells were trypsinized and seeded in a 16-well chamber slide at a density of 4×10^4^ cells per well. The following day, cells were washed twice with HBSS, starved in HBSS for 16 hours, and then stimulated for 15 minutes with high-glucose DMEM containing 4 mM glutamine and 1 μM insulin. Afterwards, cells were washed once with PBS, fixed with 4 % formaldehyde for 15 minutes and permeabilized with 0.1 % Tween-20 in PBS for 5 minutes. The PLA was performed using the Duolink In Situ Red Starter Kit Mouse/ Rabbit according to the manufacturer’s instructions. Briefly, after permeabilization, the samples were blocked, and then incubated overnight with antibodies against LAMP2 and TSC2. The following day, the samples were incubated with the MINUS and PLUS PLA probes corresponding to the primary antibodies used, followed by ligation and rolling-circle amplification in the presence of Texas-Red labeled oligos to generate the PLA signal. Finally, the samples were mounted with DAPI-containing mounting medium. All incubations were performed in a humidity chamber using a volume of 40 μL per well. The experiment was imaged with a confocal microscope (SP8, Leica); twelve stacks (7-8 μm thick with 0.3 μm spacing between consecutive layers) per condition were acquired.

For G3BP1-TSC2 and G3BP1-LAMP1 PLAs, MCF-7 CRISPR control and G3BP1 KO cells were seeded in a 16-well chamber slide at a density of 2×10^4^ cells per well. The following day, cells were washed twice with PBS, and incubated with HBSS for 16 hours. Afterwards, cells were washed once with PBS and fixed with 4% formaldehyde for 5 minutes and permeabilized with 0.1 % Trition X100 in PBS for 5 minutes. The PLA was performed as described above with antibodies against G3BP1 and TSC2 or LAMP1. The slides were analyzed using an AxioObserver Z1 compound microscope equipped with an ApoTome.2 (6 pictures per slide), 63x objective, and Axiocam 702mono and Axiocam 298 color cameras. Six stacks (0.5 μm thick) per condition were acquired.

For quantitation of all PLAs, the number of PLA puncta was counted across maximum intensity projections of raw files of each stack using CellProfiler and then normalized to the number of DAPI-positive nuclei on that field. For presentation in figures, maximum intensity projections were exported as TIFF, and brightness or contrast were adjusted for better visibility.

#### Migration assay

2-well ibidi culture-inserts were placed into 24 well plates, generating a cell-free gap of 500 μM. After knockdown induction for 4 days, 15,000 MCF-7 shControl and shG3BP1 #1 cells/ well were seeded in 100 μL full DMEM medium. 4 replicates were seeded per condition and cell line (MCF-7 shControl and shG3BP1 #1) in the presence of 2 μg/mL doxycycline to induce shRNA expression. Where indicated, rapamycin was added during seeding to a final concentration of 20 nM. After 24 hours, ibidi culture-inserts were removed and the medium was replaced with 1 mL full DMEM medium, supplemented with 20 nM rapamycin where indicated. Pictures were taken after 0, 24, and 48 hours with a Nikon ECLIPSE Ti-E/B inverted microscope, equipped with a 4x objective, using the NIS Elements version 4.13.04 software (settings: optimal frame size 1280 x 1024, no binning, 12 bit). Pictures were taken from two different regions in an automated manner by selecting the x- and y-coordinates of the 24 well plate, assuring that the same region of the scratch was monitored across all conditions. Pictures were exported as TIFF files converting the 12 bit to 16 bit and analyzed using the TScratch software and a consistent threshold of 250. For quantitation, the width of the open wound area of the 48 hours time point was normalized to the initial scratch size and expressed as the percentage of wound closure. Data was compared using a one-way ANOVA followed by a Sidak’s multiple comparisons test across n = 12 scratches from 3 independent experiments.

#### Proliferation assays

Cell proliferation was monitored using an xCELLigence real-time cell analysis (RTCA) system, allowing real-time, label free cellular analysis. After knockdown induction for 4 days, MCF-7 cells (MCF-7 shControl and shG3BP1 #1) were seeded in duplicates at a total of 2,000 cells per E-plate 16 chamber following the manufacturer’s protocol, in the presence of 2 μg/mL doxycycline. Proliferation was measured as the relative change in electrical impedance every 30 minutes for 5 days until the cells reached the stationary growth phase. Proliferation was analyzed using the RTCA software 1.2. For the presentation of the growth curves, the measured impedance was normalized to the maximum value. Data was compared using a paired two-tailed Student’s t-test across n = 6 independent experiments.

#### G3BP1 expression and survival analyses

Clinical and RNAseq data of invasive breast cancer (TCGA, provisional) were downloaded from cBio Cancer Genomics Portal (www.cbioportal.org) using the CGDS-R package (Gao et al., 2013). For 522 patients, information on the breast cancer subtype was available, of which 514 had RNAseq V2 data for *G3BP1.* A Kruskal-Wallis ANOVA by ranks was applied to evaluate subtype-dependent differences in *G3BP1* transcription.

The Kaplan Meier Plotter database (www.kmplot.com) (Gyorffy et al., 2010; Szasz et al., 2016) was used for survival analysis. Relapse free survival (RFS) was assessed in breast cancer patients based on gene expression of *G3BP1* (probeID: 225007_at), *TSC1* (probeID: 209390_at), and *TSC2* (probeID: 215735_at). Outlier gene arrays were excluded leaving 1764 patients for analysis of *G3BP1* and 3571 patients for analyses of *TSC1/TSC2.* RFS analysis in relation to G3BP1 protein expression also was based on data available in the Kaplan-Meier Plotter database, which included 126 patients. For all calculations, patients were split based on the best performing expression threshold and log-rank p-values were calculated.

#### Zebrafish maintenance and breeding

Adult zebrafish of the AB strain (Zebrafish International Resource Center) were maintained under standard aquaculture conditions in UV-sterilized water at 28.5 °C on a 14 hour light / 10 hour dark cycle. Fertilized eggs were collected via natural spawning. Embryos and larvae were raised in embryo medium, containing 1.5 mM HEPES, pH 7.6, 17.4 mM NaCl, 0.21 mM KCl, 0.12 mM MgSO4 and 0.18 mM Ca(NO_3_)_2_ in an incubator on a 14 hour light / 10 hour dark cycle at 28.5°C. For all experiments described, larvae at 0-4 days post fertilisation (dpf) were used. All zebrafish experiments were approved by the Ethics Committee of the University of Leuven (Ethische Commissie van de KU Leuven, approval number 150/2015) and by the Belgian Federal Department of Public Health, Food Safety and Environment (Federale Overheidsdienst Volksgezondheid, Veiligheid van de Voedselketen en Leefmilieu, approval number LA1210199).

For pharmacological assessment, 3 dpf larvae were individually placed into the wells of a 96 well-plate, with each well containing 100 μL of a freshly prepared 10 μM rapamycin solution in embryo medium. The untreated larvae were treated similarly with 100 μL of embryo medium.

#### Antisense morpholino knockdown

To achieve knockdown of *g3bp1* in zebrafish embryos, we used morpholino antisense oligonucleotides designed to target the Exon 2 – Intron 2 boundary of the *g3bp1* mRNA (G3BP1 MO, **Figure S5B**). The morpholino sequence, as synthesized by GeneTools was: 5’-TAACAAAGGGCAAGTCACCTGTGCA-3’. A randomized 25-nucleotide morpholino was used as a negative control (control MO). Embryos were microinjected at the one-or two-cell stage with 1 nL of either *g3bp1 or control* morpholino, corresponding to 8 ng of morpholino per injection. The morpholino concentration used was defined by titration as the highest at which the larvae displayed no morphological abnormalities. The level of knockdown in the MO injected zebrafish embryos and larvae was evaluated by PCR. For each condition, 10 embryos or larvae were pooled. RNA was extracted using Trizol and treated with DNAse I. 1 μg of total RNA was reverse transcribed using the “High Capacity cDNA Reverse Transcription” kit and random primers. The generated cDNA was then amplified with gene-specific primers for *g3bp1* and *β-actin.*

#### Zebrafish larvae lysis and immunoblotting

For RPS6-pS235/236 analysis 10 zebrafish larvae (2-3 dpf) were pooled per condition and homogenized in RIPA buffer supplemented with Complete Mini Protease Inhibitor cocktail. A Pierce BCA protein assay kit was used to determine the protein concentration of the lysates. 40 μg of protein were separated on a NuPage Novex 10% Bis-Tris gel, using SDS-PAGE with NuPage MES SDS running buffer, followed by dry transfer to an iBlot gel transfer stacks nitrocellulose membrane with an iBlot Dry Blotting System, which was then blocked for 1 hour at room temperature in Odyssey blocking buffer. Overnight incubation at 4 °C with a primary antibody against RPS6-pS235/236 was followed by incubation with Dylight secondary goat antibody to rabbit IgG for 1 hour at room temperature. A rabbit antibody against GAPDH was used as a loading control. For detection, fluorescence signal was detected using an Odyssey 2.1 imaging system (Li-Cor, USA). For graphical presentation, raw images were further processed with Adobe Photoshop version CS5.1.

#### Non-invasive local field potential (LFP) recordings

Brain activity of 4 dpf zebrafish larvae was assessed by performing non-invasive local field potential recordings, reading the electrical signal from the skin of the larva’s head (Zdebik et al., 2013). A glass pipet (containing the recording electrode), filled with artificial cerebrospinal fluid (124 mM NaCl, 2 mM KCl, 2 mM MgSO4, 2 mM CaCl2, 1.25 mM KH2PO4, 26 mM NaHCO3 and 10 mM glucose), was positioned on the skin above the optic tectum using a stereomicroscope. The differential signal between the recording electrode and the reference electrode was amplified 10,000 times by DAGAN 2400 amplifier (Minnesota, USA), band pass filtered at 0.3-300 Hz and digitized at 2 kHz via a PCI-6251 interface (National Instruments, UK) with WinEDR (John Dempster, University of Strathclyde, UK). Recordings lasted for 10 minutes and were analyzed with Clampfit 10.2 software (Molecular Devices Corporation, USA). A polyspiking discharge was scored positive when its amplitude exceeded three times the amplitude of the baseline and it had a duration of at least 50 ms.

In addition, power spectral density (PSD) analysis of the recordings was performed using MatLab R2018 (MATrix LABoratory, USA) software (Hunyadi et al., 2017). In brief, the power spectral density of the signals were estimated using Welch’s method of averaging modified periodograms with 512-point fast fourier transform of 80% overlapping 100 sample (100 ms) long segments and a Hamming window. Next, the PSD estimate of each LFP recording was summed over each 10 Hz frequency band, ranging from 0 to 100 Hz. PSD estimates were normalized against the untreated control MO injected larvae and the data were plotted as mean (±SEM) PSD per condition over the 20-80 Hz range. Outliers were identified via the Iterative Grubbs test (alpha = 0.1).

### Quantitation and Statistical Analysis

#### Immunoblot quantitation

Quantitation of raw images taken with a LAS-4000 camera system or FUSION FX7 with DarQ-9 camera was performed using ImageQuant TL Version 8.1. Background subtraction was performed using the rolling ball method with a defined radius of 200 for all images. Quantitation of raw images taken with a ChemiDoc XRS+ camera system was performed using Image Lab version 5.2.1. For all images, pixel values of a single lane were normalized to the average value of all lanes, and then normalized to the loading control Tubulin if not indicated otherwise in the figure legends. Quantitation of raw images taken with an Odyssey 2.1 imaging system (Li-Cor) was performed using Image Studio Lite Version 5.2. For images from immunoblot analysis of zebrafish samples, pixel values of a single lane were normalized to the single value of the loading control GAPDH and then to the control MO within each experiment.

#### Protein sequence analysis

To analyse the sequence similarities between human G3BP1 (Uniprot ID: Q13283) and human G3BP2 (Q9UN86) and their domains, or between human and zebrafish G3BP1 (Q6P124) EMBOSS Needle (https://www.ebi.ac.uk/Tools/psa/emboss_needle/) with the Blosum62 matrix was used. Visualization of sequence alignments was done using Texshade based on a ClustalW alignment of the whole protein sequences. The domains indicated for G3BP1 were based on Reineke and Lloyd (2015).

#### Phylogenetic analysis

To identify possible orthologues in other species, the human proteins G3BP1 (Uniprot ID: Q13283), G3BP2 (Q9UN86), TSC1 (Q92574), TSC2 (P49815), TBC1D7 (Q9P0N9), RHEB (Q15382), and MTOR (P42345) were used as query proteins for a blastp+ search (Camacho et al., 2009) against the NCBI non-redundant protein sequence database (nr; version 2017-11). The e-value cut-off for identified proteins was 1e-30 with a maximum of 20,000 target sequences, disabled low-complexity filtering, using the BLOSUM62 matrix, a word size of 6 and gap opening and extension costs of 11 and 1, respectively. The results were parsed and filtered using custom Python scripts (https://github.com/MolecularBioinformatics/Phylogenetic-analysis) and manually curated as described earlier (Bockwoldt et al., 2019).

#### Statistical analysis

GraphPad Prism version 7.04 or 8.03 was used for statistical analysis and statistical presentation of quantitations. Where two conditions were compared, a paired two-tailed Student’s t-test was performed. If more than two conditions were compared, a one-way ANOVA followed by a Sidak’s multiple comparisons test was applied. In the case of immunoblot time courses or PSD analysis with equal intervals more than two conditions were compared using a two-way ANOVA. For bar graphs, the corresponding dot plots were overlaid. For *G3BP1* expression analysis a Kruskal-Wallis ANOVA by ranks was performed using Dell Statistica version 13. For the analysis of relapse-free survival the Kaplan-Meier Plotter was used and a log-rank test was applied. For each experiment, the number of replicates and the statistical test applied is indicated in the figure legend.

#### Data availability

All data are available from the corresponding authors upon reasonable request.

## Supplemental figure legends

**Figure S1. G3BP1 does not alter mTORC1 activity upon arsenite stress, related to Figure 1.**

**(A)** Amino acid sequence of G3BP1. G3BP1’s five protein domains are indicated according to Reineke and Lloyd (2015) and highlighted in blue, green, brown, yellow and pink. G3BP1 peptides identified in MTOR IPs by mass spectrometry (Schwarz et al., 2015) are shown in red. In total, 20 unique peptides were identified with a sequence coverage of 58.4%.

**(B)** Representative annotated MS2 spectrum of the identified G3BP1 peptide DFFQSYGNVVELR. The respective peptide sequence is highlighted with a red frame in the full-length amino acid sequence of G3BP1 depicted in (**A**).

**(C)** IPs from MCF-7 cells with antibodies against MTOR or mock (rat IgG). Data shown are representative of n = 6 biological replicates.

**(D)** IPs from MCF-7 cells with antibodies against RPTOR (RPTOR#1 or #2) or mock (rat IgG). Data shown are representative of n = 3 biological replicates.

**(E)** Nucleotide sequence of G3BP1. The targeting sequences of the four different siRNAs from the G3BP1 siRNA pool (light green), two shRNA sequences against G3BP1 (dark green), and the sgRNA used for CRIPSR/Cas9 mediated knockout (orange) are highlighted.

**(F)** Time course analysis of shG3BP1 #1 or shControl MCF-7 cells that were serum starved and exposed to 500 μM arsenite for up to 60 minutes. Data shown are representative of n = 3 biological replicates.

**(G)** Quantitation of G3BP1 immunoblot data shown in (**F**). Data are shown as the mean ± SEM. G3BP1 levels (black and green curve), were compared between shControl and shG3BP1 #1 cells, using a two-way ANOVA across n = 3 biological replicates. p-values are presented at the bottom of the graph.

**(H)** Quantitation of RPS6KB1-pT389 immunoblot data shown in (**F**). RPS6KB1-pT389 levels (black and blue curve) are represented and compared between shControl and shG3BP1 #1 cells as described in (**G**).

**(I)** Time course analysis of siG3BP1 and siControl transfected MCF-7 cells exposed to 500 μM arsenite for up to 60 minutes. Data shown are representative of n = 3 biological replicates.

**(J)** Quantitation of G3BP1 immunoblot data shown in (**I**). Data are shown as the mean ± SEM. Protein levels were normalized to the average intensity of all lanes, and then to the loading control GAPDH. G3BP1 levels (black and green curve) were compared between siControl and siG3BP1 cells, using a two-way ANOVA across n = 3 biological replicates. p-values are presented at the bottom of the graphs.

**(K)** Quantitation of RPS6KB1-pST389 immunoblot data shown in (I). RPS6KB1-pT389 levels (black and blue bars) are represented and compared between siControl and siG3BP1 cells as described in (J).

**Figure S2. G3BP1 suppresses mTORC1 activation by insulin and amino acids in the absence of stress granules, related to Figure 1.**

**(A)** G3BP1 knockdown was induced in MCF-7 cells harboring a second shRNA sequence (shG3BP1 #2, **see Figure S1E)** targeting another exon than shG3BP1 #1. Cells were serum and amino acid starved, and stimulated with 100 nM insulin / aa for the indicated time periods. Data shown are representative of n = 5 biological replicates.

**(B)** Quantitation of G3BP1 immunoblot data shown in (**A**). Data are shown as the mean ± SEM and overlaid with the single data points represented as dot plots. G3BP1 levels (black and green bars) were compared between shControl and shG3BP1 #2 cells, using a one-way ANOVA followed by a Sidak’s multiple comparisons test across n = 5 biological replicates. p-values are presented above the corresponding bar graphs.

**(C)** Quantitation of RPS6KB1-pT389 immunoblot data shown in (**A**). RPS6KB1-pT389 levels (black and blue bars) are represented and compared between shControl and shG3BP1 #2 cells as described in (**B**).

**(D)** Quantitation of RPS6-pS235/236 immunoblot data shown in (**A**). RPS6-pS235/236 levels (black and blue bars) are represented and compared between shControl and shG3BP1 #2 cells as described in (**B**).

**(E)** G3BP1 knockdown was induced in MDA-MB-231 cells harboring a second shRNA sequence (shG3BP1 #2) targeting another exon than shG3BP1 #1. Cells were serum and amino acid starved, and stimulated with 100 nM insulin / aa for the indicated time periods. Data shown are representative of n = 4 biological replicates.

**(F)** Quantitation of G3BP1 immunoblot data shown in (**E**). Data are shown as the mean ± SEM and overlaid with the single data points represented as dot plots. G3BP1 levels (black and green bars) were compared between shControl and shG3BP1 #2 cells, using a one-way ANOVA followed by a Sidak’s multiple comparisons test across n = 4 biological replicates. p-values are presented above the corresponding bar graphs.

**(G)** Quantitation of RPS6KB1-pT389 immunoblot data shown in (**E**). RPS6KB1-pT389 levels (black and blue bars) are represented and compared between shControl and shG3BP1 #2 cells as described in (**F**).

**(H)** Quantitation of RPS6-pS235/236 immunoblot data shown in (**E**). RPS6-pS235/236 levels (black and blue bars) are represented and compared between shControl and shG3BP1 #2 cells as described in (**F**).

**(I)** siG3BP1 and siControl transfected MCF-7 cells were serum and amino acid starved, and stimulated with 100 nM insulin / aa for 15 minutes. Data shown are representative of n = 6 biological replicates.

**(J)** Quantitation of G3BP1 immunoblot data shown in (**I**). Data are shown as the mean ± SEM and overlaid with the single data points represented as dot plots. G3BP1 levels (black and green bars) were compared between siControl and siG3BP1 cells, using a one-way ANOVA followed by a Sidak’s multiple comparisons test across n = 6 biological replicates. p-values are presented above the corresponding bar graphs.

**(K)** Quantitation of RPS6KB1-pT389 immunoblot data shown in (**I**). RPS6KB1-pT389 levels (black and blue bars) are represented and compared between siControl and siG3BP1 cells as described in (**J**).

**(L)** Quantitation of RPS6-pS235/236 immunoblot data shown in (**I**). RPS6-pS235/236 levels (black and blue bars) are represented and compared between siControl and siG3BP1 cells as described in (**J**).

**(M)** Schematic diagram of sgRNA designed for the G3BP1 locus. The sequence of the sgRNA is indicated in green. The locations of the nuclease-specific protospacer adjacent motif (PAM) sequence is indicated in orange.

**(N)** IF analysis of shControl and shG3BP1 #1 MCF-7 cells. Cells were either serum and amino acid starved and stimulated with 100 nM insulin / aa for 15 minutes; or serum starved and treated with 500 μM arsenite for 30 minutes. Scale bar 10 μm. White regions in merged images, co-localization of EIF3A and G3BP1. Inserts, magnifications of the area in the yellow squares in the merged pictures. Nuclei were stained with Hoechst. Representative images are shown for n = 3 biological replicates.

**(O)** Quantitation of data shown in (N). Data are shown as the mean ± SEM and overlaid with the single data points represented as dot plots. The number of SG / cell was analyzed using the eIF3A channel. shControl and shG3BP1 #1 cells were compared using a one-way ANOVA followed by a Sidak’s multiple comparisons test across n = 9 pictures from n = 3 biological replicates. p-values are presented above the corresponding bar graphs.

**Figure S3: BiFC constructs and their expression, related to Figure 4.**

**(A)** Scheme of the plasmids used for BiFC analyses in **Figure 4E**. If the two BiFC interaction partners A and B are in close proximity, the C-terminal and N-terminal fragments of mLumin bind and enable the fluorophore to reconstitute, which can be detected by fluorescence microscopy. A = pGW-myc-LC151 (G3BP1 fused to C-terminal mLumin). B = pGW-HA-LN151 (N-terminal mLumin, control; or N-terminal mLumin fused to MTOR, LAMP1, LAMP2, or TSC2).

**(B)** Immunoblot analysis of the expression of BiFC fusion proteins used i4 Figure 4E. HEK293T cells were transfected with plasmids carrying G3BP1 fused to the C-terminal mLumin fragment, together with an N-terminal mLumin fragment fused to TSC2, LAMP1, LAMP2 or MTOR, or an N-terminal mLumin fragment only (control). Data shown are representative of n = 3 biological replicates.

**Figure S4. G3BP2 phenocopies G3BP1 effects in the TSC complex-mTORC1 axis, related to Figure 5.**

**(A)** Sequence alignment of human G3BP1 and G3BP2. Protein domains are indicated according to Reineke and Lloyd (2015). High similarity is highlighted in blue. PxxP motifs are indicated in green.

**(B)** Sequence similarities of human G3BP1 (Q13283) and G3BP2 (Q9UN86). Sequence alignments of the domains were done based on the domain regions (AS in G3BP1) defined for G3BP1 in Reineke and Lloyd (2015).

**(C)** IPs from MCF-7 cells with antibodies against MTOR or mock (rat IgG). Data shown are representative of n = 3 biological replicates.

**(D)** Scheme of the plasmids used for BiFC analyses in **Figure 5B**. If the two BiFC interaction partners A and B are in close proximity, the C-terminal and N-terminal fragments of mLumin bind and enable the fluorophore to reconstitute, which can be detected by fluorescence microscopy. A = pGW-myc-LC151 (G3BP2 fused to C-terminal mLumin). B = pGW-HA-LN151 (N-terminal mLumin, control; or N-terminal mLumin fused to MTOR, LAMP1, LAMP2, or TSC2).

**(E)** Expression of BiFC fusion proteins used in Figure 5B. HEK293T cells were transfected with plasmids carrying G3BP2 fused to the C-terminal mLumin fragment, together with an N-terminal mLumin fragment fused to TSC2, LAMP1, LAMP2 or MTOR, or an N-terminal mLumin fragment only (control). Data shown are representative of n = 3 biological replicates.

**Figure S5. Zebrafish G3BP1: sequence alignment, generation of G3BP1 morpholino, and epileptogenic events, related to Figure 6.**

**(A)** Sequence alignment of human and zebrafish G3BP1. Protein domains are indicated according to Reineke and Lloyd (2015). High similarity is highlighted in blue. PxxP motifs are indicated in green. The sequences share 67,8 % similarity and 77,4 % identity.

**(B)** Scheme of the generation of G3BP1 morpholino (G3BP1 MO). The G3BP1 MO was designed to target the Exon 2 – Intron 2 boundary of the *g3bp1* mRNA, interfering with normal splicing, leading to a knockdown (G3BP1 MO).

**(C)** Zebrafish larvae were injected with G3BP1 MO and non-invasive local field potentials were recorded from larval optic tecta at 3 dpf for 10 minutes. Representative 10 minutes recording of G3BP1 MO with magnification of a polyspiking event is shown.

**(D)** Zebrafish larvae were injected with Control MO and non-invasive local field potentials were recorded from larval optic tecta at 3 dpf for 10 minutes. Representative 10 minutes recording of Control MO with magnification of a polyspiking event is shown.

## References

Alam, U., and Kennedy, D. (2019). Rasputin a decade on and more promiscuous than ever? A review of G3BPs. Biochim Biophys Acta Mol Cell Res 1866, 360–370.

Anderson, P., and Kedersha, N. (2002). Stressful initiations. J Cell Sci 115, 3227–3234.

Anderson, P., and Kedersha, N. (2008). Stress granules: the Tao of RNA triage. Trends Biochem Sci 33, 141–150.

Anderson, P., Kedersha, N., and Ivanov, P. (2015). Stress granules, P-bodies and cancer. Biochim Biophys Acta 1849, 861–870.

Annibaldi, A., Dousse, A., Martin, S., Tazi, J., and Widmann, C. (2011). Revisiting G3BP1 as a RasGAP binding protein: sensitization of tumor cells to chemotherapy by the RasGAP 317-326 sequence does not involve G3BP1. PLoS One 6, e29024.

Appenzeller, S., Balling, R., Barisic, N., Baulac, S., Caglayan, H., Craiu, D., De Jonghe, P., Depienne, C., Dimova, P., Djémié, T., et al. (2014). De novo mutations in synaptic transmission genes including DNM1 cause epileptic encephalopathies. Am J Hum Genet 95, 360–370.

Astrinidis, A., Cash, T.P., Hunter, D.S., Walker, C.L., Chernoff, J., and Henske, E.P. (2002). Tuberin, the tuberous sclerosis complex 2 tumor suppressor gene product, regulates Rho activation, cell adhesion and migration. Oncogene 21, 8470–8476.

Avruch, J., Hara, K., Lin, Y., Liu, M., Long, X., Ortiz-Vega, S., and Yonezawa, K. (2006). Insulin and amino-acid regulation of mTOR signaling and kinase activity through the Rheb GTPase. Oncogene 25, 6361–6372.

Baldassari, S., Licchetta, L., Tinuper, P., Bisulli, F., and Pippucci, T. (2016). GATOR1 complex: the common genetic actor in focal epilepsies. J Med Genet 53, 503–510.

Baselga, J., Campone, M., Piccart, M., Burris, H.A., 3rd, Rugo, H.S., Sahmoud, T., Noguchi, S., Gnant, M., Pritchard, K.I., Lebrun, F., et al. (2012). Everolimus in postmenopausal hormone-receptor-positive advanced breast cancer. N Engl J Med 366, 520–529.

Baulac, S. (2016). mTOR signaling pathway genes in focal epilepsies. Prog Brain Res 226, 61–79.

Betz, C., and Hall, M.N. (2013). Where is mTOR and what is it doing there? J Cell Biol 203, 563–574.

Bikkavilli, R.K., and Malbon, C.C. (2011). Arginine methylation of G3BP1 in response to Wnt3a regulates beta-catenin mRNA. J Cell Sci 124, 2310–2320.

Bley, N., Lederer, M., Pfalz, B., Reinke, C., Fuchs, T., Glass, M., Moller, B., and Huttelmaier, S. (2015). Stress granules are dispensable for mRNA stabilization during cellular stress. Nucleic Acids Res 43, e26.

Bockwoldt, M., Houry, D., Niere, M., Gossmann, T.I., Reinartz, I., Schug, A., Ziegler, M., and Heiland, I. (2019). Identification of evolutionary and kinetic drivers of NAD-dependent signaling. Proc Natl Acad Sci U S A 116, 15957–15966.

Borkowska, J., Schwartz, R.A., Kotulska, K., and Jozwiak, S. (2011). Tuberous sclerosis complex: tumors and tumorigenesis. Int J Dermatol 50, 13–20.

Buchan, J.R., and Parker, R. (2009). Eukaryotic stress granules: the ins and outs of translation. Mol Cell 36, 932–941.

Camacho, C., Coulouris, G., Avagyan, V., Ma, N., Papadopoulos, J., Bealer, K., and Madden, T.L. (2009). BLAST+: architecture and applications. BMC Bioinformatics 10, 421.

Carroll, B., Maetzel, D., Maddocks, O.D., Otten, G., Ratcliff, M., Smith, G.R., Dunlop, E.A., Passos, J.F., Davies, O.R., Jaenisch, R., et al. (2016). Control of TSC2-Rheb signaling axis by arginine regulates mTORC1 activity. Elife 5.

Chu, J., Zhang, Z., Zheng, Y., Yang, J., Qin, L., Lu, J., Huang, Z.L., Zeng, S., and Luo, Q. (2009). A novel far-red bimolecular fluorescence complementation system that allows for efficient visualization of protein interactions under physiological conditions. Biosens Bioelectron 25, 234–239.

Condon, K.J., and Sabatini, D.M. (2019). Nutrient regulation of mTORC1 at a glance. J Cell Sci 132.

Crino, P.B. (2015). mTOR signaling in epilepsy: insights from malformations of cortical development. Cold Spring Harb Perspect Med 5.

Crino, P.B. (2016). The mTOR signalling cascade: paving new roads to cure neurological disease. Nat Rev Neurol 12, 379–392.

Curatolo, P., D’Argenzio, L., Cerminara, C., and Bombardieri, R. (2008). Management of epilepsy in tuberous sclerosis complex. Expert Rev Neurother 8, 457–467.

Curatolo, P., Moavero, R., and de Vries, P.J. (2015). Neurological and neuropsychiatric aspects of tuberous sclerosis complex. Lancet Neurol 14, 733–745.

Debaize, L., Jakobczyk, H., Rio, A.G., Gandemer, V., and Troadec, M.B. (2017). Optimization of proximity ligation assay (PLA) for detection of protein interactions and fusion proteins in non-adherent cells: application to pre-B lymphocytes. Mol Cytogenet 10, 27.

Demetriades, C., Doumpas, N., and Teleman, A.A. (2014). Regulation of TORC1 in response to amino acid starvation via lysosomal recruitment of TSC2. Cell 156, 786–799.

Demetriades, C., Plescher, M., and Teleman, A.A. (2016). Lysosomal recruitment of TSC2 is a universal response to cellular stress. Nat Commun 7, 10662.

Dibble, C.C., Elis, W., Menon, S., Qin, W., Klekota, J., Asara, J.M., Finan, P.M., Kwiatkowski, D.J., Murphy, L.O., and Manning, B.D. (2012). TBC1D7 is a third subunit of the TSC1-TSC2 complex upstream of mTORC1. Mol Cell 47, 535–546.

Dou, N., Chen, J., Yu, S., Gao, Y., and Li, Y. (2016). G3BP1 contributes to tumor metastasis via upregulation of Slug expression in hepatocellular carcinoma. Am J Cancer Res 6, 2641–2650.

Eskelinen, E.L. (2006). Roles of LAMP-1 and LAMP-2 in lysosome biogenesis and autophagy. Mol Aspects Med 27, 495–502.

Forman-Kay, J.D., and Mittag, T. (2013). From sequence and forces to structure, function, and evolution of intrinsically disordered proteins. Structure 21, 1492–1499.

Friend, S., and Royce, M. (2016). The Changing Landscape of Breast Cancer: How Biology Drives Therapy. Medicines 3, 2.

Gallouzi, I.E., Parker, F., Chebli, K., Maurier, F., Labourier, E., Barlat, I., Capony, J.P., Tocque, B., and Tazi, J. (1998). A novel phosphorylation-dependent RNase activity of GAP- SH3 binding protein: a potential link between signal transduction and RNA stability. Mol Cell Biol 18, 3956–3965.

Gao, J., Aksoy, B.A., Dogrusoz, U., Dresdner, G., Gross, B., Sumer, S.O., Sun, Y., Jacobsen, A., Sinha, R., Larsson, E., et al. (2013). Integrative analysis of complex cancer genomics and clinical profiles using the cBioPortal. Sci Signal 6, pl1.

Garami, A., Zwartkruis, F.J., Nobukuni, T., Joaquin, M., Roccio, M., Stocker, H., Kozma, S.C., Hafen, E., Bos, J.L., and Thomas, G. (2003). Insulin activation of Rheb, a mediator of mTOR/S6K/4E-BP signaling, is inhibited by TSC1 and 2. Mol Cell 11, 1457–1466.

Geback, T., Schulz, M.M., Koumoutsakos, P., and Detmar, M. (2009). TScratch: a novel and simple software tool for automated analysis of monolayer wound healing assays. Biotechniques 46, 265–274.

Ghosh, S., Tergaonkar, V., Rothlin, C.V., Correa, R.G., Bottero, V., Bist, P., Verma, I.M., and Hunter, T. (2006). Essential role of tuberous sclerosis genes TSC1 and TSC2 in NF-kappaB activation and cell survival. Cancer Cell 10, 215–226.

Goncharova, E.A., Goncharov, D.A., Lim, P.N., Noonan, D., and Krymskaya, V.P. (2006). Modulation of cell migration and invasiveness by tumor suppressor TSC2 in lymphangioleiomyomatosis. Am J Respir Cell Mol Biol 34, 473–480.

Grabiner, B.C., Nardi, V., Birsoy, K., Possemato, R., Shen, K., Sinha, S., Jordan, A., Beck, A.H., and Sabatini, D.M. (2014). A diverse array of cancer-associated MTOR mutations are hyperactivating and can predict rapamycin sensitivity. Cancer Discov 4, 554–563.

Gyorffy, B., Lanczky, A., Eklund, A.C., Denkert, C., Budczies, J., Li, Q., and Szallasi, Z. (2010). An online survival analysis tool to rapidly assess the effect of 22,277 genes on breast cancer prognosis using microarray data of 1,809 patients. Breast Cancer Res Treat 123, 725–731.

Hao, F., Kondo, K., Itoh, T., Ikari, S., Nada, S., Okada, M., and Noda, T. (2018). Rheb localized on the Golgi membrane activates lysosome-localized mTORC1 at the Golgilysosome contact site. J Cell Sci 131.

Heberle, A.M., Razquin Navas, P., Langelaar-Makkinje, M., Kasack, K., Sadik, A., Faessler, E., Hahn, U., Marx-Stoelting, P., Opitz, C.A., Sers, C., et al. (2019). The PI3K and MAPK/p38 pathways control stress granule assembly in a hierarchical manner. Life Sci Alliance 2.

Heyne, H.O., Singh, T., Stamberger, H., Abou Jamra, R., Caglayan, H., Craiu, D., De Jonghe, P., Guerrini, R., Helbig, K.L., Koeleman, B.P.C., et al. (2018). De novo variants in neurodevelopmental disorders with epilepsy. Nat Genet 50, 1048–1053.

Hojgaard, C., Sorensen, H.V., Pedersen, J.S., Winther, J.R., and Otzen, D.E. (2018). Can a Charged Surfactant Unfold an Uncharged Protein? Biophys J 115, 2081–2086.

Holz, M.K., and Blenis, J. (2005). Identification of S6 kinase 1 as a novel mammalian target of rapamycin (mTOR)-phosphorylating kinase. J Biol Chem 280, 26089–26093.

Hoxhaj, G., and Manning, B.D. (2019). The PI3K-AKT network at the interface of oncogenic signalling and cancer metabolism. Nat Rev Cancer.

Hoyle, N.P., Castelli, L.M., Campbell, S.G., Holmes, L.E., and Ashe, M.P. (2007). Stressdependent relocalization of translationally primed mRNPs to cytoplasmic granules that are kinetically and spatially distinct from P-bodies. J Cell Biol 179, 65–74.

Hu, C.D., Chinenov, Y., and Kerppola, T.K. (2002). Visualization of interactions among bZIP and Rel family proteins in living cells using bimolecular fluorescence complementation. Mol Cell 9, 789–798.

Huang, J., and Manning, B.D. (2008). The TSC1-TSC2 complex: a molecular switchboard controlling cell growth. Biochem J 412, 179–190.

Huang, J., Yang, J., Lei, Y., Gao, H., Wei, T., Luo, L., Zhang, F., Chen, H., Zeng, Q., and Guo, L. (2016). An ANCCA/PRO2000-miR-520a-E2F2 regulatory loop as a driving force for the development of hepatocellular carcinoma. Oncogenesis 5, e229.

Hunyadi, B., Siekierska, A., Sourbron, J., Copmans, D., and de Witte, P.A.M. (2017). Automated analysis of brain activity for seizure detection in zebrafish models of epilepsy. J Neurosci Methods 287, 13–24.

Inoki, K., Li, Y., Xu, T., and Guan, K.L. (2003). Rheb GTPase is a direct target of TSC2 GAP activity and regulates mTOR signaling. Genes Dev 17, 1829–1834.

Inoki, K., Ouyang, H., Zhu, T., Lindvall, C., Wang, Y., Zhang, X., Yang, Q., Bennett, C., Harada, Y., Stankunas, K., et al. (2006). TSC2 integrates Wnt and energy signals via a coordinated phosphorylation by AMPK and GSK3 to regulate cell growth. Cell 126, 955–968.

Irvine, K., Stirling, R., Hume, D., and Kennedy, D. (2004). Rasputin, more promiscuous than ever: a review of G3BP. Int J Dev Biol 48, 1065–1077.

Jedrusik-Bode, M., Studencka, M., Smolka, C., Baumann, T., Schmidt, H., Kampf, J., Paap, F., Martin, S., Tazi, J., Muller, K.M., et al. (2013). The sirtuin SIRT6 regulates stress granule formation in C. elegans and mammals. J Cell Sci 126, 5166–5177.

Jozwiak, S., Kotulska, K., Wong, M., and Bebin, M. (2019). Modifying genetic epilepsies - Results from studies on tuberous sclerosis complex. Neuropharmacology 166, 107908.

Kedersha, N., and Anderson, P. (2007). Mammalian stress granules and processing bodies. Methods Enzymol 431, 61–81.

Kedersha, N., Panas, M.D., Achorn, C.A., Lyons, S., Tisdale, S., Hickman, T., Thomas, M., Lieberman, J., McInerney, G.M., Ivanov, P., et al. (2016). G3BP-Caprin1-USP10 complexes mediate stress granule condensation and associate with 40S subunits. J Cell Biol 212, 845–860.

Kedersha, N., Tisdale, S., Hickman, T., and Anderson, P. (2008). Real-time and quantitative imaging of mammalian stress granules and processing bodies. Methods Enzymol 448, 521–552.

Kennedy, D., French, J., Guitard, E., Ru, K., Tocque, B., and Mattick, J. (2001). Characterization of G3BPs: tissue specific expression, chromosomal localisation and rasGAP(120) binding studies. J Cell Biochem 84, 173–187.

Kim, J., and Guan, K.L. (2019). mTOR as a central hub of nutrient signalling and cell growth. Nat Cell Biol 21, 63–71.

Koboldt, D.C., Fulton, R.S., McLellan, M.D., Schmidt, H., Kalicki-Veizer, J., McMichael, J.F., Fulton, L.L., Dooling, D.J., Ding, L., Mardis, E.R., et al. (2012). Comprehensive molecular portraits of human breast tumours. Nature 490, 61–70.

Kwiatkowski, D.J. (2003). Tuberous sclerosis: from tubers to mTOR. Ann Hum Genet 67, 87–96.

Kwiatkowski, D.J., and Wagle, N. (2015). mTOR Inhibitors in Cancer: What Can We Learn from Exceptional Responses? EBioMedicine 2, 2–4.

Liao, Y.C., Fernandopulle, M.S., Wang, G., Choi, H., Hao, L., Drerup, C.M., Patel, R., Qamar, S., Nixon-Abell, J., Shen, Y., et al. (2019). RNA Granules Hitchhike on Lysosomes for Long-Distance Transport, Using Annexin A11 as a Molecular Tether. Cell 179, 147–164 e120.

LiCausi, F., and Hartman, N.W. (2018). Role of mTOR Complexes in Neurogenesis. Int J Mol Sci 19.

Liu, G.Y., and Sabatini, D.M. (2020). mTOR at the nexus of nutrition, growth, ageing and disease. Nat Rev Mol Cell Biol.

Long, X., Lin, Y., Ortiz-Vega, S., Yonezawa, K., and Avruch, J. (2005). Rheb binds and regulates the mTOR kinase. Curr Biol 15, 702–713.

Ma, L., Chen, Z., Erdjument-Bromage, H., Tempst, P., and Pandolfi, P.P. (2005). Phosphorylation and functional inactivation of TSC2 by Erk implications for tuberous sclerosis and cancer pathogenesis. Cell 121, 179–193.

Mahboubi, H., and Stochaj, U. (2017). Cytoplasmic stress granules: Dynamic modulators of cell signaling and disease. Biochim Biophys Acta 1863, 884–895.

Marcotte, L., and Crino, P.B. (2006). The neurobiology of the tuberous sclerosis complex. Neuromolecular Med 8, 531–546.

Martin, S., Zekri, L., Metz, A., Maurice, T., Chebli, K., Vignes, M., and Tazi, J. (2013). Deficiency of G3BP1, the stress granules assembly factor, results in abnormal synaptic plasticity and calcium homeostasis in neurons. J Neurochem 125, 175–184.

Matsuki, H., Takahashi, M., Higuchi, M., Makokha, G.N., Oie, M., and Fujii, M. (2013). Both G3BP1 and G3BP2 contribute to stress granule formation. Genes Cells 18, 135–146.

Menon, S., Dibble, C.C., Talbott, G., Hoxhaj, G., Valvezan, A.J., Takahashi, H., Cantley, L.C., and Manning, B.D. (2014). Spatial control of the TSC complex integrates insulin and nutrient regulation of mTORC1 at the lysosome. Cell 156, 771–785.

Meric-Bernstam, F., Akcakanat, A., Chen, H., Do, K.A., Sangai, T., Adkins, F., Gonzalez-Angulo, A.M., Rashid, A., Crosby, K., Dong, M., et al. (2012). PIK3CA/PTEN mutations and Akt activation as markers of sensitivity to allosteric mTOR inhibitors. Clin Cancer Res 18, 1777–1789.

Min, L., Ruan, Y., Shen, Z., Jia, D., Wang, X., Zhao, J., Sun, Y., and Gu, J. (2015). Overexpression of Ras-GTPase-activating protein SH3 domain-binding protein 1 correlates with poor prognosis in gastric cancer patients. Histopathology 67, 677–688.

Molle, K.-D. (2006). Regulation of the mammalian target of rapamycin complex 2 (mTORC2). In Department Biozentrum (Basel: University of Basel), pp. 92.

Moon, S.L., Morisaki, T., Khong, A., Lyon, K., Parker, R., and Stasevich, T.J. (2019). Multicolour single-molecule tracking of mRNA interactions with RNP granules. Nat Cell Biol 21, 162–168.

Mossmann, D., Park, S., and Hall, M.N. (2018). mTOR signalling and cellular metabolism are mutual determinants in cancer. Nat Rev Cancer 18, 744–757.

Nellist, M., van Slegtenhorst, M.A., Goedbloed, M., van den Ouweland, A.M., Halley, D.J., and van der Sluijs, P. (1999). Characterization of the cytosolic tuberin-hamartin complex. Tuberin is a cytosolic chaperone for hamartin. J Biol Chem 274, 35647–35652.

Neve, R.M., Chin, K., Fridlyand, J., Yeh, J., Baehner, F.L., Fevr, T., Clark, L., Bayani, N., Coppe, J.P., Tong, F., et al. (2006). A collection of breast cancer cell lines for the study of functionally distinct cancer subtypes. Cancer Cell 10, 515–527.

Orlova, K.A., and Crino, P.B. (2010). The tuberous sclerosis complex. Ann N Y Acad Sci 1184, 87–105.

Panas, M.D., Kedersha, N., Schulte, T., Branca, R.M., Ivanov, P., and Anderson, P. (2019). Phosphorylation of G3BP1-S149 does not influence stress granule assembly. J Cell Biol 218, 2425–2432.

Paplomata, E., and O’Regan, R. (2014). The PI3K/AKT/mTOR pathway in breast cancer: targets, trials and biomarkers. Ther Adv Med Oncol 6, 154–166.

Parker, F., Maurier, F., Delumeau, I., Duchesne, M., Faucher, D., Debussche, L., Dugue, A., Schweighoffer, F., and Tocque, B. (1996). A Ras-GTPase-activating protein SH3-domain-binding protein. Molecular and cellular biology 16, 2561–2569.

Pende, M., Um, S.H., Mieulet, V., Sticker, M., Goss, V.L., Mestan, J., Mueller, M., Fumagalli, S., Kozma, S.C., and Thomas, G. (2004). S6K1(-/-)/S6K2(-/-) mice exhibit perinatal lethality and rapamycin-sensitive 5’-terminal oligopyrimidine mRNA translation and reveal a mitogen-activated protein kinase-dependent S6 kinase pathway. Mol Cell Biol 24, 3112–3124.

Peng, M., Yin, N., and Li, M.O. (2017). SZT2 dictates GATOR control of mTORC1 signalling. Nature 543, 433–437.

Prigent, M., Barlat, I., Langen, H., and Dargemont, C. (2000). IkappaBalpha and IkappaBalpha /NF-kappa B complexes are retained in the cytoplasm through interaction with a novel partner, RasGAP SH3-binding protein 2. J Biol Chem 275, 36441–36449.

Rabanal-Ruiz, Y., and Korolchuk, V.I. (2018). mTORC1 and Nutrient Homeostasis: The Central Role of the Lysosome. Int J Mol Sci 19.

Reineke, L.C., and Lloyd, R.E. (2015). The stress granule protein G3BP1 recruits protein kinase R to promote multiple innate immune antiviral responses. J Virol 89, 2575–2589.

Reineke, L.C., and Neilson, J.R. (2019). Differences between acute and chronic stress granules, and how these differences may impact function in human disease. Biochem Pharmacol 162, 123–131.

Roach, E.S., and Kwiatkowski, D.J. (2016). Seizures in tuberous sclerosis complex: hitting the target. Lancet 388, 2062–2064.

Sadowski, K., Kotulska-Jozwiak, K., and Jozwiak, S. (2015). Role of mTOR inhibitors in epilepsy treatment. Pharmacol Rep 67, 636–646.

Sancak, Y., Bar-Peled, L., Zoncu, R., Markhard, A.L., Nada, S., and Sabatini, D.M. (2010). Ragulator-Rag complex targets mTORC1 to the lysosomal surface and is necessary for its activation by amino acids. Cell 141, 290–303.

Sancak, Y., Thoreen, C.C., Peterson, T.R., Lindquist, R.A., Kang, S.A., Spooner, E., Carr, S.A., and Sabatini, D.M. (2007). PRAS40 is an insulin-regulated inhibitor of the mTORC1 protein kinase. Mol Cell 25, 903–915.

Sanjana, N.E., Shalem, O., and Zhang, F. (2014). Improved vectors and genome-wide libraries for CRISPR screening. Nat Methods 11, 783–784.

Scheldeman, C., Mills, J.D., Siekierska, A., Serra, I., Copmans, D., Iyer, A.M., Whalley, B.J., Maes, J., Jansen, A.C., Lagae, L., et al. (2017). mTOR-related neuropathology in mutant tsc2 zebrafish: Phenotypic, transcriptomic and pharmacological analysis. Neurobiol Dis 108, 225–237.

Schwarz, J.J., Wiese, H., Tolle, R.C., Zarei, M., Dengjel, J., Warscheid, B., and Thedieck, K. (2015). Functional Proteomics Identifies Acinus L as a Direct Insulin- and Amino Acid-Dependent Mammalian Target of Rapamycin Complex 1 (mTORC1) Substrate. Mol Cell Proteomics 14, 2042–2055.

Somasekharan, S.P., El-Naggar, A., Leprivier, G., Cheng, H., Hajee, S., Grunewald, T.G., Zhang, F., Ng, T., Delattre, O., Evdokimova, V., et al. (2015). YB-1 regulates stress granule formation and tumor progression by translationally activating G3BP1. J Cell Biol 208, 913–929.

Szasz, A.M., Lanczky, A., Nagy, A., Forster, S., Hark, K., Green, J.E., Boussioutas, A., Busuttil, R., Szabo, A., and Gyorffy, B. (2016). Cross-validation of survival associated biomarkers in gastric cancer using transcriptomic data of 1,065 patients. Oncotarget 7, 49322–49333.

Takahashi, M., Higuchi, M., Matsuki, H., Yoshita, M., Ohsawa, T., Oie, M., and Fujii, M. (2013). Stress granules inhibit apoptosis by reducing reactive oxygen species production. Mol Cell Biol 33, 815–829.

Tee, A.R. (2018). The Target of Rapamycin and Mechanisms of Cell Growth. Int J Mol Sci 19.

Tee, A.R., Manning, B.D., Roux, P.P., Cantley, L.C., and Blenis, J. (2003). Tuberous sclerosis complex gene products, Tuberin and Hamartin, control mTOR signaling by acting as a GTPase-activating protein complex toward Rheb. Curr Biol 13, 1259–1268.

Tee, A.R., Sampson, J.R., Pal, D.K., and Bateman, J.M. (2016). The role of mTOR signalling in neurogenesis, insights from tuberous sclerosis complex. Semin Cell Dev Biol 52, 12–20.

Thedieck, K., Holzwarth, B., Prentzell, M.T., Boehlke, C., Klasener, K., Ruf, S., Sonntag, A.G., Maerz, L., Grellscheid, S.N., Kremmer, E., et al. (2013). Inhibition of mTORC1 by astrin and stress granules prevents apoptosis in cancer cells. Cell 154, 859–874.

Thien, A., Prentzell, M.T., Holzwarth, B., Klasener, K., Kuper, I., Boehlke, C., Sonntag, A.G., Ruf, S., Maerz, L., Nitschke, R., et al. (2015). TSC1 activates TGF-beta-Smad2/3 signaling in growth arrest and epithelial-to-mesenchymal transition. Dev Cell 32, 617–630.

Tourriere, H., Chebli, K., Zekri, L., Courselaud, B., Blanchard, J.M., Bertrand, E., and Tazi, J. (2003). The RasGAP-associated endoribonuclease G3BP assembles stress granules. J Cell Biol 160, 823–831.

van der Poest Clement, E., Jansen, F.E., Braun, K.P.J., and Peters, J.M. (2020). Update on Drug Management of Refractory Epilepsy in Tuberous Sclerosis Complex. Paediatr Drugs 22, 73–84.

Wagle, N., Grabiner, B.C., Van Allen, E.M., Hodis, E., Jacobus, S., Supko, J.G., Stewart, M., Choueiri, T.K., Gandhi, L., Cleary, J.M., et al. (2014). Activating mTOR mutations in a patient with an extraordinary response on a phase I trial of everolimus and pazopanib. Cancer Discov 4, 546–553.

Wang, X., and Proud, C.G. (1997). p70 S6 kinase is activated by sodium arsenite in adult rat cardiomyocytes: roles for phosphatidylinositol 3-kinase and p38 MAP kinase. Biochem Biophys Res Commun 238, 207–212.

Wang, Y., Fu, D., Chen, Y., Su, J., Wang, Y., Li, X., Zhai, W., Niu, Y., Yue, D., and Geng, H. (2018). G3BP1 promotes tumor progression and metastasis through IL-6/G3BP1/STAT3 signaling axis in renal cell carcinomas. Cell Death Dis 9, 501.

Weiler, M., Blaes, J., Pusch, S., Sahm, F., Czabanka, M., Luger, S., Bunse, L., Solecki, G., Eichwald, V., Jugold, M., et al. (2014). mTOR target NDRG1 confers MGMT-dependent resistance to alkylating chemotherapy. Proc Natl Acad Sci U S A 111, 409–414.

Winslow, S., Leandersson, K., and Larsson, C. (2013). Regulation of PMP22 mRNA by G3BP1 affects cell proliferation in breast cancer cells. Mol Cancer 12, 156.

Wippich, F., Bodenmiller, B., Trajkovska, M.G., Wanka, S., Aebersold, R., and Pelkmans, L. (2013). Dual specificity kinase DYRK3 couples stress granule condensation/dissolution to mTORC1 signaling. Cell 152, 791–805.

Wolfson, R.L., Chantranupong, L., Wyant, G.A., Gu, X., Orozco, J.M., Shen, K., Condon, K.J., Petri, S., Kedir, J., Scaria, S.M., et al. (2017). KICSTOR recruits GATOR1 to the lysosome and is necessary for nutrients to regulate mTORC1. Nature 543, 438–442.

Wong, M., and Crino, P.B. (2012). mTOR and Epileptogenesis in Developmental Brain Malformations. In Jasper’s Basic Mechanisms of the Epilepsies, th, J.L. Noebels, M. Avoli, M.A. Rogawski, R.W. Olsen, and A.V. Delgado-Escueta, eds. (Bethesda (MD)).

Wyant, G.A., Abu-Remaileh, M., Frenkel, E.M., Laqtom, N.N., Dharamdasani, V., Lewis, C.A., Chan, S.H., Heinze, I., Ori, A., and Sabatini, D.M. (2018). NUFIP1 is a ribosome receptor for starvation-induced ribophagy. Science 360, 751–758.

Yang, X., Shen, Y., Garre, E., Hao, X., Krumlinde, D., Cvijovic, M., Arens, C., Nystrom, T., Liu, B., and Sunnerhagen, P. (2014). Stress granule-defective mutants deregulate stress responsive transcripts. PLoS Genet 10, e1004763.

Zdebik, A.A., Mahmood, F., Stanescu, H.C., Kleta, R., Bockenhauer, D., and Russell, C. (2013). Epilepsy in kcnj10 morphant zebrafish assessed with a novel method for long-term EEG recordings. PLoS One 8, e79765.

Zekri, L., Chebli, K., Tourriere, H., Nielsen, F.C., Hansen, T.V., Rami, A., and Tazi, J. (2005). Control of fetal growth and neonatal survival by the RasGAP-associated endoribonuclease G3BP. Mol Cell Biol 25, 8703–8716.

Zhang, H., Ma, Y., Zhang, S., Liu, H., He, H., Li, N., Gong, Y., Zhao, S., Jiang, J.D., and Shao, R.G. (2015). Involvement of Ras GTPase-activating protein SH3 domain-binding protein 1 in the epithelial-to-mesenchymal transition-induced metastasis of breast cancer cells via the Smad signaling pathway. Oncotarget 6, 17039–17053.

Zhang, H., Zhang, S., He, H., Zhao, W., Chen, J., and Shao, R.G. (2012). GAP161 targets and downregulates G3BP to suppress cell growth and potentiate cisplaitin-mediated cytotoxicity to colon carcinoma HCT116 cells. Cancer Sci 103, 1848–1856.

Zhang, J., Kim, J., Alexander, A., Cai, S., Tripathi, D.N., Dere, R., Tee, A.R., Tait-Mulder, J., Di Nardo, A., Han, J.M., et al. (2013). A tuberous sclerosis complex signalling node at the peroxisome regulates mTORC1 and autophagy in response to ROS. Nat Cell Biol 15, 1186–1196.

Zhang, P., Fan, B., Yang, P., Temirov, J., Messing, J., Kim, H.J., and Taylor, J.P. (2019). Chronic optogenetic induction of stress granules is cytotoxic and reveals the evolution of ALS-FTD pathology. Elife 8.

Zhang, Y., Gao, X., Saucedo, L.J., Ru, B., Edgar, B.A., and Pan, D. (2003). Rheb is a direct target of the tuberous sclerosis tumour suppressor proteins. Nat Cell Biol 5, 578–581.

