## Supplemental Figures and Key Resource Table for "G3BP1 tethers the TSC complex to lysosomes and suppresses mTORC1 in the absence of stress granules"

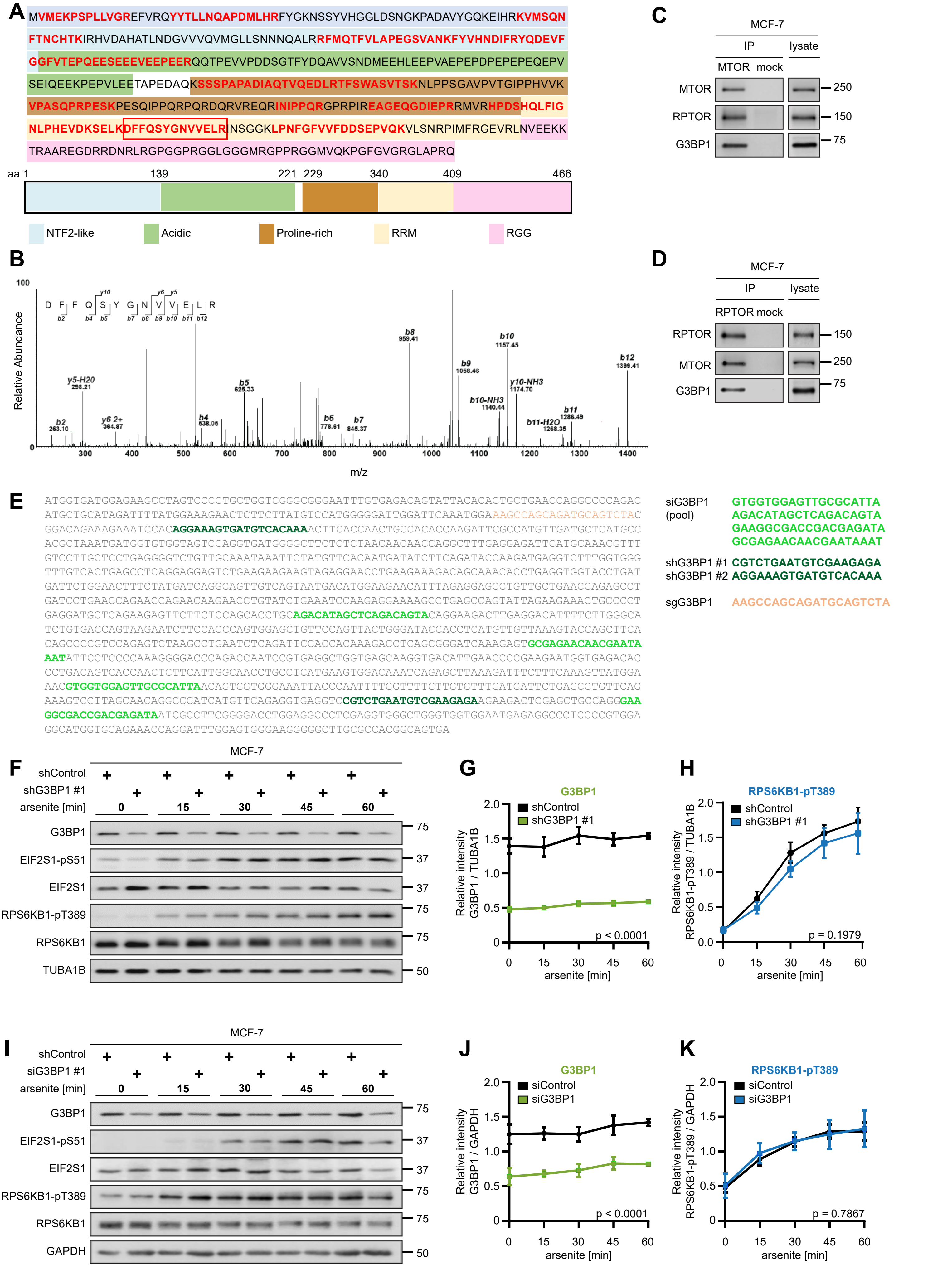
A
MVMEKPSPLLVGREFVRQYYTLLNQAPDMLHRFYGKNSSYVHGGLDSNGKPADAVYGQKEIHRKVMSQN
FTNCHTKIRHVDAHATLNDGVVVQVMGLLSNNNQALRRFMQTFVLAPEGSVANKFYVHNDIFRYQDEVF
GGFVTEPQEESEEEVEEPEERQQTPEVVPDDSGTFYDQAVVSNDMEEHLEEPVAEPEPDPEPEPEQEPV
SEIQEEKPEPVLEETAPEDAQKSSSPAPADIAQTVQEDLRTFSWASVTSKNLPPSGAVPVTGIPPHVVK
VPASQPRPESKPESQIPPQRPQRDQRVREQRINIPPQRGPRPIREAGEQGDIEPRRMVRHPDSHQLFIG
NLPHEVDKSELKDFFQSYGNVVELRINSGGKLPNFGFVVFDDSEPVQKVLSNRPIMFRGEVRLNVEEKK
TRAAREGDRRDNRLRGPGGPRGGLGGGMRGPPRGGMVQKPGFGVGRGLAPRQ
aa 1 139 221 229 340 409 466
NTF2-like
Acidic
Proline-rich
RRM
RGG
B
DFFQSYGNVVELR
Relative Abundance
100
0
y5-H2O 298.21
b2 263.10
y6 2+ 364.87
b4 538.06
b5 625.33
b6 778.61
b7 845.37
b8 959.41
b9 1058.46
b10-NH3 1140.44
b10 1157.45
y10-NH3 1174.70
b11-H2O 1268.35
b11 1286.49
b12 1399.41
300 400 500 600 700 800 900 1000 1100 1200 1300 1400
m/z
C
MCF-7
IP lysate
MTOR mock
MTOR
RPTOR
G3BP1
250
150
75
D
MCF-7
IP lysate
RPTOR mock
RPTOR
MTOR
G3BP1
150
250
75
E
ATGGTGATGGAGAAGCCTAGTCCCCTGCTGGTCGGGCGGGAATTTGTGAGACAGTATTACACACTGCTGAACCAGGCCCCAGAC
ATGCTGCATAGATTTTATGGAAAGAACTCTTCTTATGTCCATGGGGGATTGGATTCAAATGGAAAGCCAGCAGATGCAGTCTAC
GGACAGAAAGAAATCCACAGGAAAGTGATGTCACAAAACTTCACCAACTGCCACACCAAGATTCGCCATGTTGATGCTCATGCC
ACGCTAAATGATGGTGTGGTAGTCCAGGTGATGGGGCTTCTCTCTAACAACAACCAGGCTTTGAGGAGATTCATGCAAACGTTT
GTCCTTGCTCCTGAGGGGTCTGTTGCAAATAAATTCTATGTTCACAATGATATCTTCAGATACCAAGATGAGGTCTTTGGTGGG
TTTGTCACTGAGCCTCAGGAGGAGTCTGAAGAAGAAGTAGAGGAACCTGAAGAAAGACAGCAAACACCTGAGGTGGTACCTGAT
GATTCTGGAACTTTCTATGATCAGGCAGTTGTCAGTAATGACATGGAAGAACATTTAGAGGAGCCTGTTGCTGAACCAGAGCCT
GATCCTGAACCAGAACCAGAACAAGAACCTGTATCTGAAATCCAAGAGGAAAAGCCTGAGCCAGTATTAGAAGAAACTGCCCCT
GAGGATGCTCAGAAGAGTTCTTCTCCAGCACCTGCAGACATAGCTCAGACAGTACAGGAAGACTTGAGGACATTTTCTTGGGCA
TCTGTGACCAGTAAGAATCTTCCACCCAGTGGAGCTGTTCCAGTTACTGGGATACCACCTCATGTTGTTAAAGTACCAGCTTCA
CAGCCCCGTCCAGAGTCTAAGCCTGAATCTCAGATTCCACCACAAAGACCTCAGCGGGATCAAAGAGTGCGAGAACAACGAATA
AATATTCCTCCCCAAAGGGGACCCAGACCAATCCGTGAGGCTGGTGAGCAAGGTGACATTGAACCCCGAAGAATGGTGAGACAC
CCTGACAGTCACCAACTCTTCATTGGCAACCTGCCTCATGAAGTGGACAAATCAGAGCTTAAAGATTTCTTTCAAAGTTATGGA
AACGTGGTGGAGTTGCGCATTAACAGTGGTGGGAAATTACCCAATTTTGGTTTTGTTGTGTTTGATGATTCTGAGCCTGTTCAG
AAAGTCCTTAGCAACAGGCCCATCATGTTCAGAGGTGAGGTCCGTCTGAATGTCGAAGAGAAGAAGACTCGAGCTGCCAGGGAA
GGCGACCGACGAGATAATCGCCTTCGGGGACCTGGAGGCCCTCGAGGTGGGCTGGGTGGTGGAATGAGAGGCCCTCCCCGTGGA
GGCATGGTGCAGAAACCAGGATTTGGAGTGGGAAGGGGGCTTGCGCCACGGCAGTGA
siG3BP1 (pool) GTGGTGGAGTTGCGCATTA
AGACATAGCTCAGACAGTA
GAAGGCGACCGACGAGATA
GCGAGAACAACGAATAAAT
shG3BP1 #1 CGTCTGAATGTCGAAGAGA
shG3BP1 #2 AGGAAAGTGATGTCACAAA
sgG3BP1 AAGCCAGCAGATGCAGTCTA
F
MCF-7
shControl
shG3BP1 #1
arsenite [min]
0 15 30 45 60
G3BP1
EIF2S1-pS51
EIF2S1
RPS6KB1-pT389
RPS6KB1
TUBA1B
75
37
37
75
75
50
G
G3BP1
shControl
shG3BP1 #1
Relative intensity G3BP1 / TUBA1B
2.0 1.5 1.0 0.5 0
p < 0.0001
0 15 30 45 60
arsenite [min]
H
RPS6KB1-pT389
shControl
shG3BP1 #1
Relative intensity RPS6KB1-pT389 / TUBA1B
2.0 1.5 1.0 0.5 0.0
p = 0.1979
0 15 30 45 60
arsenite [min]
I
MCF-7
shControl
siG3BP1 #1
arsenite [min]
0 15 30 45 60
G3BP1
EIF2S1-pS51
EIF2S1
RPS6KB1-pT389
RPS6KB1
GAPDH
75
37
37
75
75
50
J
G3BP1
siControl
siG3BP1
Relative intensity G3BP1 / GAPDH
2.0 1.5 1.0 0.5 0
p < 0.0001
0 15 30 45 60
arsenite [min]
K
RPS6KB1-pT389
siControl
siG3BP1
Relative intensity RPS6KB1-pT389 / GAPDH
2.0 1.5 1.0 0.5 0
p = 0.7867
0 15 30 45 60
arsenite [min]

### Figure S1

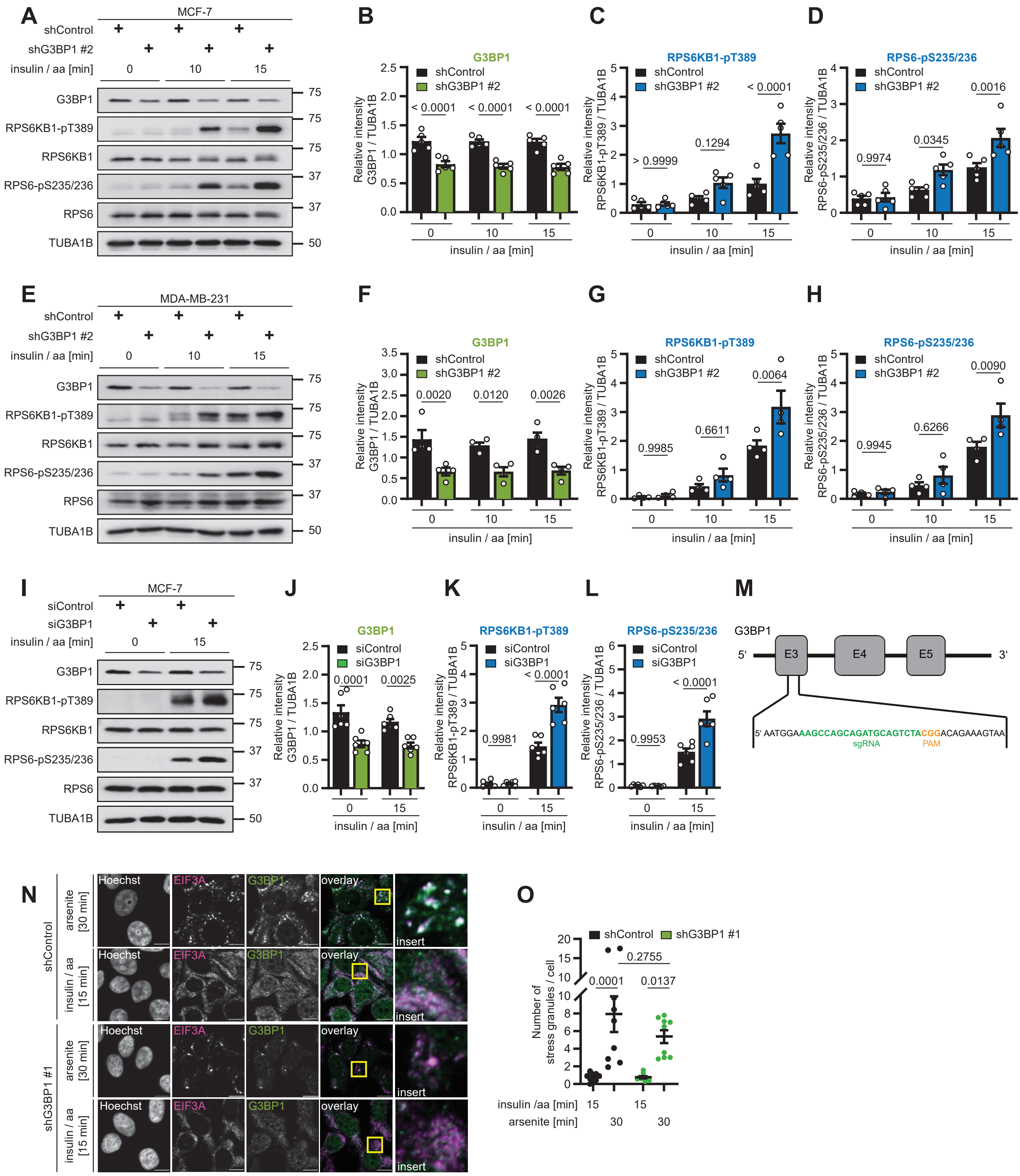
A
MCF-7
shControl
shG3BP1 #2
insulin / aa [min]
0
10
15
G3BP1
RPS6KB1-pT389
RPS6KB1
RPS6-pS235/236
RPS6
TUBA1B
B
G3BP1
C
RPS6KB1-pT389
D
RPS6-pS235/236
Relative intensity
insulin / aa [min]
E
MDA-MB-231
F
G
H
I
siControl
siG3BP1
J
K
L
M
5' AATGGAAAGCCAGCAGATGCAGTCTACGGACAGAAAGTAA
sgRNA
PAM
N
Hoechst
EIF3A
overlay
insert
arsenite [30 min]
insulin / aa [15 min]
shG3BP1 #1
O
Number of stress granules / cell
insulin /aa [min]
arsenite [min]

### Figure S2

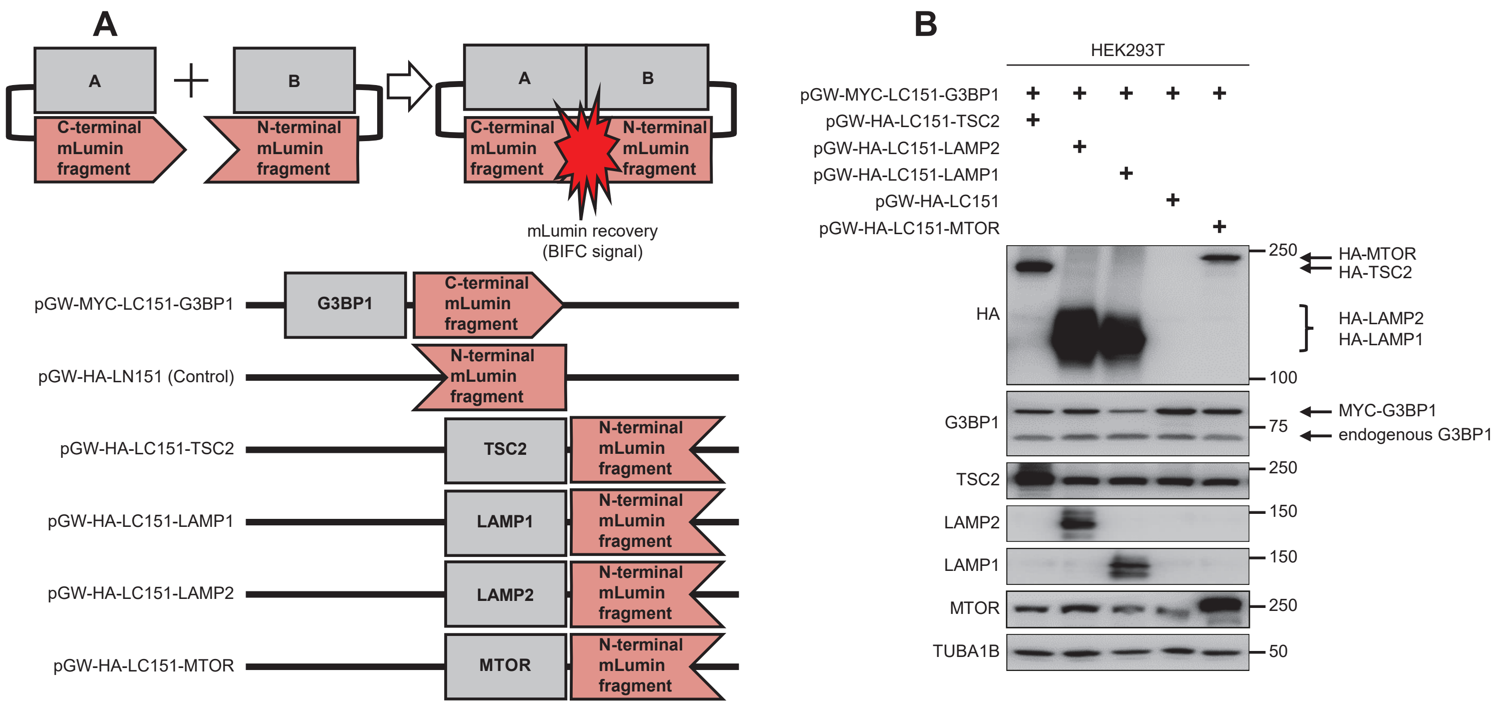
A
A
B
C-terminal mLumin fragment
N-terminal mLumin fragment
mLumin recovery (BIFC signal)
pGW-MYC-LC151-G3BP1
G3BP1
pGW-HA-LN151 (Control)
pGW-HA-LC151-TSC2
TSC2
pGW-HA-LC151-LAMP1
LAMP1
pGW-HA-LC151-LAMP2
LAMP2
pGW-HA-LC151-MTOR
MTOR
B
HEK293T
pGW-MYC-LC151-G3BP1
pGW-HA-LC151-TSC2
pGW-HA-LC151-LAMP2
pGW-HA-LC151-LAMP1
pGW-HA-LC151
pGW-HA-LC151-MTOR
HA
G3BP1
TSC2
LAMP2
LAMP1
MTOR
TUBA1B
250
100
75
150
50
HA-MTOR
HA-TSC2
HA-LAMP2
HA-LAMP1
MYC-G3BP1
endogenous G3BP1

**Figure S3**

**A**

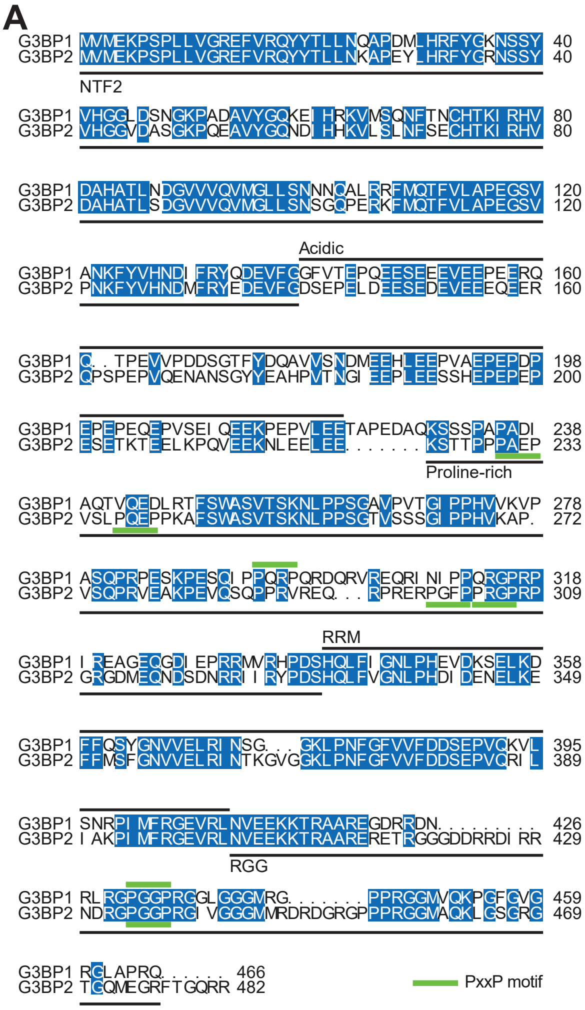


**B**

| Domain | G3BP1 (amino acids) | G3BP2 (amino acids) | Identity | Similarity | Gaps |
|---|---|---|---|---|---|
| Full length | 1-466 | 1-482 | 63.9 % | 74.4 % | 7.7 % |
| NTF2 | 1-139 | 1-139 | 82.0 % | 92.8 % | 0.0 % |
| Acidic | 140-221 | 140-223 | 45.3 % | 62.8 % | 7.0 % |
| Proline-rich | 229-339 | 224-330 | 58.6 % | 65.5 % | 3.6 % |
| RRM | 340-408 | 331-402 | 76.4 % | 91.7 % | 4.2 % |
| RGG | 409-466 | 403-482 | 58.1 % | 60.8 % | 21.6 % |

**C**

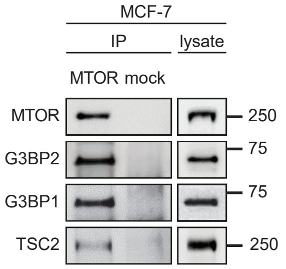


**D**

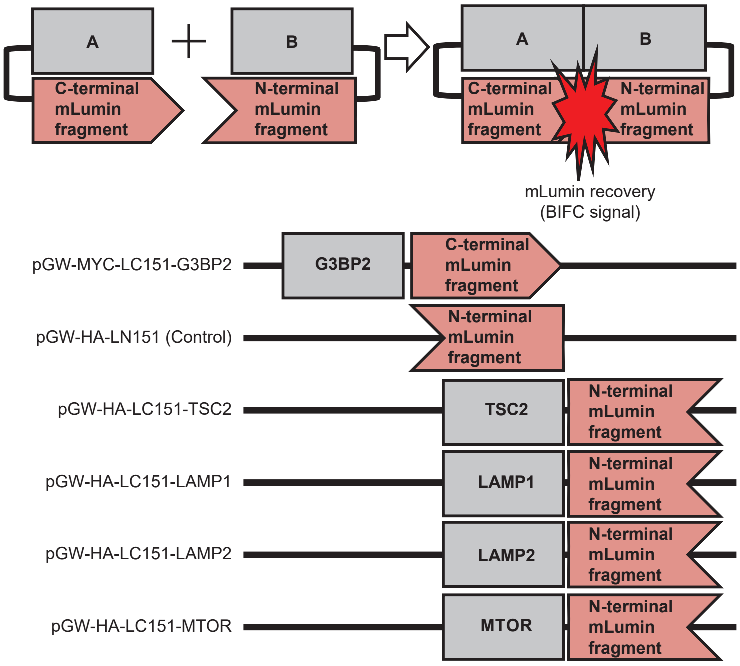


**E**

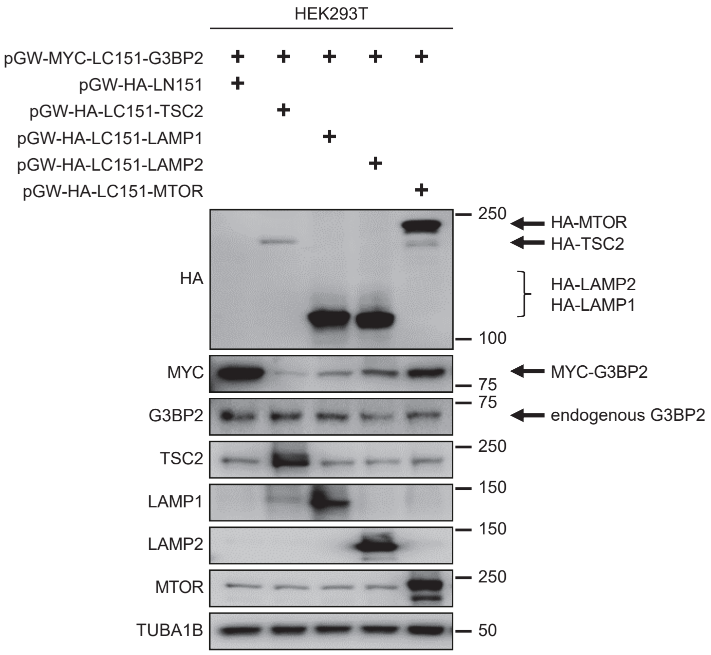


**Figure S4**

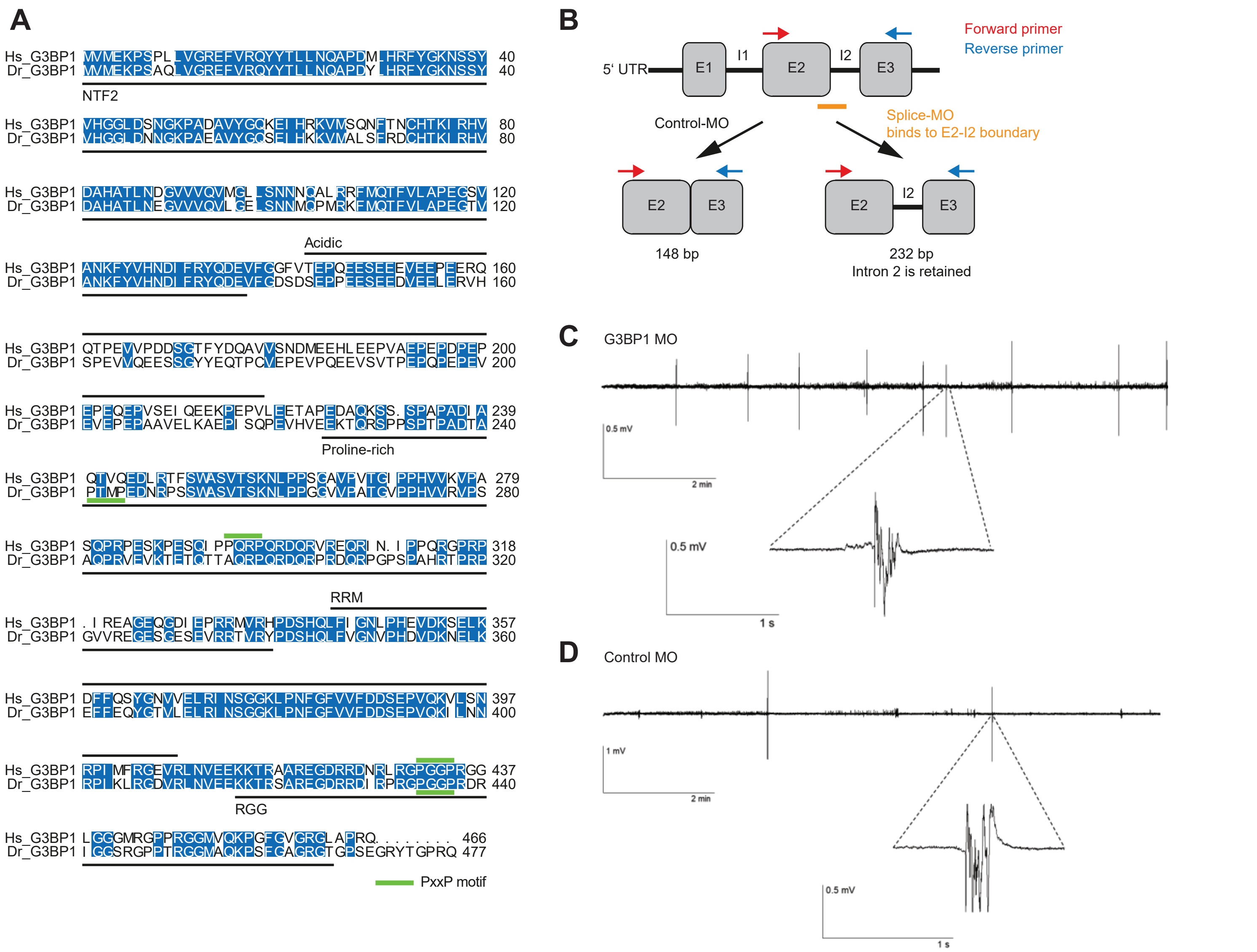
A
Hs_G3BP1 MVMEKPSPLLVGREFVRQYYTLLNQAPDMLHRFYGKNSSY 40
Dr_G3BP1 MVMEKPSAQLVGREFVRQYYTLLNQAPDYLHRFYGKNSSY 40
NTF2
Hs_G3BP1 VHGGLDSNGKPADAVYGQKEIHRKVMSQNFTNCHTKIRHV 80
Dr_G3BP1 VHGGLDNNGKPAEAVYGQSELHKKVMALSFRDCHTKIRHV 80
Hs_G3BP1 DAHATLNDGVVVQVMGLLSNNNQALRRFMQTFVLAPEGSV 120
Dr_G3BP1 DAHATLNEGVVVQVLGELSNNMQPMRKFMQTFVLAPEGTV 120
Acidic
Hs_G3BP1 ANKFYVHNDIFRYQDEVFGGFVTEPQEESEEEVEEPEERQ 160
Dr_G3BP1 ANKFYVHNDIFRYQDEVFGDSDSEPPEESEEDVEELERVH 160
Hs_G3BP1 QTPEVVPDDSGTFYDQAVVSNDMEEHLEEPVAEPEPDPEP 200
Dr_G3BP1 SPEVVQEESSGYYEQTPCVEPEVPQEEVSVTPEPQPEPEV 200
Hs_G3BP1 EPEQEPVSEIQEEKPEPVLEETAPEDAQKSS.SPAPADIA 239
Dr_G3BP1 EVEPEPAAVELKAEPISQPEVHVEEKTQRSPPSPTPADTA 240
Proline-rich
Hs_G3BP1 QTVQEDLRTFSWASVTSKNLPPSGAVPVTGIPPHVVKVPA 279
Dr_G3BP1 PTMPEDNRPSSWASVTSKNLPPGGVVPAITGVPPHVVRVPS 280
Hs_G3BP1 SQPRPESKPESQIPPQRPQRDQRVREQRIN.IPPQRGPRP 318
Dr_G3BP1 AQPRVEVKTETQTTAQRPQRDQRPRDQRPGPSPAHRTPRP 320
RRM
Hs_G3BP1 .IREAGEQGDIEPRRMVRHPDSHQLFIGNLPHEVDKSELK 357
Dr_G3BP1 GVVREGESGESEVRRTVRYPDSHQLFVGNVPHDVDKNELK 360
Hs_G3BP1 DFFQSYGNVVELRINSGGKLPNFGFVVFDDSEPVQKVLSN 397
Dr_G3BP1 EFFEQYGTVLELRINSGGKLPNFGFVVFDDSEPVQKILNN 400
Hs_G3BP1 RPIMFRGEVRLNVEEKKTRAAREGDRRDNRLRGPGGPRGG 437
Dr_G3BP1 RPIKLRGDVRLNVEEKKTRSAREGDRRDIRPRGPGGPRDR 440
RGG
Hs_G3BP1 LGGGMRGPPRGGMVQKPGFGVGRGLAPRQ........ 466
Dr_G3BP1 IGGSRGPPTRGGMAQKPSFGAGRGTGPSEGRYTGPRQ 477
PxxP motif
B
Forward primer
Reverse primer
5‘ UTR
E1
I1
E2
I2
E3
Control-MO
Splice-MO
binds to E2-I2 boundary
E2
E3
148 bp
E2
I2
E3
232 bp
Intron 2 is retained
C
G3BP1 MO
0.5 mV
2 min
0.5 mV
1 s
D
Control MO
1 mV
2 min
0.5 mV
1 s

### Figure S5

#### STAR METHODS

##### Key Resources Table

| REAGENT or RESOURCE | | SOURCE | IDENTIFIER |
|---|---|---|---|
| Antibodies | | | |
| EIF2S1 | dilution in IB 1:1000 | Cell Signaling | Cat# 9722; RRID: AB_2230924 |
| EIF2S1-pS51 | dilution in IB 1:1000 | Cell Signaling | Cat# 9721; RRID: AB_330951 |
| EIF3A | dilution in IB 1:10000; dilution in IF 1:1000 | Cell Signaling | Cat# 3411; RRID: AB_2096523 |
| FLAG | used in IP at 1ug/mL | Sigma-Aldrich | Cat# F3165; RRID: AB_259529 |
| G3BP1 | dilution in IB 1:1000 | Santa Cruz | Cat# sc-365338; RRID: AB_10846950 |
| G3BP1 | dilution in IB 1:1000; dilution in IF 1:200; dilution in PLA 1:2000 | Santa Cruz | Cat# sc-81940; RRID: AB_1123055 |
| G3BP2 | dilution in IB 1:1000 | Bethyl | Cat# A302-040A; RRID: AB_1576545 |
| GAPDH | dilution in IB 1:10000 | Abcam | Cat# ab37187; RRID: AB_732651 |
| GAPDH (zebrafish) | dilution in IB 1:1000 | Sigma-Aldrich | Cat# SAB2701826 |
| GFP | dilution in IB 1:1000; used in IP at 1 µg/mL | Roche | Cat# 11814460001; RRID: AB_390913 |
| Goat anti-Mouse IgG (H+L) Cross-Adsorbed Secondary Antibody, Alexa Fluor 488 | dilution in IF 1:500 | Invitrogen | Cat# A-11001; RRID: AB_2534069 |
| Goat anti-Rabbit IgG (H+L) Cross-Adsorbed Secondary Antibody, Alexa Fluor 568 | dilution in IF 1:500 | Invitrogen | Cat# A-11011; RRID: AB_143157 |
| Goat anti-Mouse IgG (H+L) cross-adsorbed secondary, Alexa Fluor 555 | dilution in IF 1:1000 | Thermo Fisher Scientific | Cat# A-21422; RRID: AB_2535844 |

| | | | |
|---|---|---|---|
| Goat anti-Rabbit IgG (H+L) cross-adsorbed secondary, Alexa Fluor 488 | dilution in IF 1:1000 | Thermo Fisher Scientific | Cat# A-11008; RRID: AB_143165 |
| Goat anti-Mouse IgG (H+L) Secondary Antibody, HRP-coupled | dilution in IB 1:4000 | Thermo Fisher Scientific | Cat# 31430; RRID: AB_228307 |
| Goat anti-Rabbit IgG (H+L) Secondary Antibody, HRP-coupled | dilution in IB 1:4000 | Thermo Fisher Scientific | Cat# 31460; RRID: AB_228341 |
| Goat anti-Rabbit IgG (H+L) Secondary Antibody, Dylight 800 (zebrafish) | dilution in IB 1:10000 | Thermo Fisher Scientific | Cat# SA5-35571; RRID: AB_2556775 |
| HA | dilution in IB 1:1000 | Roche | Cat# 11867423001; RRID: AB_390918 |
| Histone H3 (H3C1) | dilution in IB 1:1000 | Bethyl | Cat# A300-822A; RRID: AB_597872 |
| HSP90 (CDC37) | dilution in IB 1:10000 | Cell Signaling | Cat# 4877; RRID: AB_2233307 |
| LMNA A/C | dilution in IB 1:10000 | Cell Signaling | Cat# 2032; RRID: AB_2136278 |
| LAMP1 | dilution for IB 1:1000; dilution for PLA 1:200 | Cell Signaling | Cat# 9091; RRID: AB_2687579 |
| LAMP1 | dilution in IF 1:1000 | Developmental Studies Hybridoma Bank | Cat# H4A3; RRID: AB_2296838 |
| LAMP2 | dilution for IB 1:1000; dilution in IF 1:200 | Santa Cruz | Cat# sc-18822; RRID: AB_626858 |
| LAMP2 | dilution for PLA 1:200 | Developmental Studies Hybridoma Bank | Cat# H4B4, RRID: AB_2134755 |
| MTOR | dilution in IB 1:1000 | Cell Signaling | Cat# 2983; RRID: AB_2105622 |
| MTOR epitope maps to residues 221 and 261 of human mTOR | used in IP at 7.5 µg/mL | Monoclonal Antibody Core Unit. Helmholtz Zentrum München, Germany | TQREP-3G6 |
| Mock antibody mouse | used in IP at 7.5 µg/mL | Santa Cruz | Cat# sc-2025; RRID: AB_737182 |

| | | | |
|---|---|---|---|
| Mock antibody rabbit | used in IP at 7.5 µg/mL | Bethyl | Cat# P120-101; RRID: AB_479829 |
| | used in IP from rat brains at 4 µg/mL | Sigma-Aldrich | Cat# I5006; RRID: AB_1163659 |
| Mock antibody rat | used in IP at 7.5 µg/mL | Monoclonal Antibody Core Unit. Helmholtz Zentrum München, Germany | RmC3-7H8 |
| MYC-tag | dilution in IB 1:1000 | Cell Signaling | Cat# 2276; RRID: AB_331783 |
| RPS6KB1 | dilution in IB 1:1000 | Cell Signaling | Cat# 2708; RRID: AB_390722 |
| RPS6KB1-pT389 | dilution in IB 1:1000 | Cell Signaling | Cat# 9206; RRID: AB_2285392 |
| RPS6KB1-pT389 | dilution in IB 1:1000 | Cell Signaling | Cat# 9205; RRID: AB_330944 |
| RAB5A | dilution in IB 1:1000 | Cell Signaling | Cat# 3547; RRID: AB_2300649 |
| RAB7A | dilution in IB 1:1000 | Cell Signaling | Cat# 9367; RRID: AB_1904103 |
| RPTOR | dilution in IB 1:1000 | Cell Signaling | Cat# 2280; RRID: AB_561245 |
| RPTOR #1 epitope maps to residues 686 and 704 of human Raptor | used in IP at 7.5 µg/mL | Monoclonal Antibody Core Unit. Helmholtz Zentrum München, Germany | RAP1-20C4 |
| RPTOR #2 | used in IP at 7.5 µg/mL | Bethyl | Cat# A300-553A; RRID: AB_2130793 |
| RPS6 | dilution for IB 1:1000 | Cell Signaling | Cat# 2317; RRID: AB_2238583 |
| RPS6-pS235/236 | dilution for IB 1:1000 | Cell Signaling | Cat# 4856; RRID: AB_2181037 |
| S6-pS235/236 (zebrafish) | dilution for IB 1:1000 | Cell Signaling | Cat# 2211; RRID: AB_331679 |
| 4x Sample buffer for TSC1 IP in rat brain tissue | 40% glycerol, 2% β-mercaptoethanol, 8% SDS, 240 mM Tris-HCl pH 6.8, and bromophenol blue | N/A | N/A |

| | | | |
|---|---|---|---|
| 5x Sample buffer | 10% glycerol, 1% β-mercaptoethanol, 1.7% SDS, 62.5 mM Tris-HCl pH 6.8, and bromophenol blue | N/A | N/A |
| Sample buffer for GFP-IPs | 25 mM Tris-HCl pH 6.8; 4% (w/v) SDS; 3% (w/v) DTT; 0.02% (v/v) bromophenol blue | N/A | N/A |
| TSC1 | dilution for IB 1:1000 | Cell Signaling | Cat# 4906; RRID: AB_2209790 |
| TSC1 #1 | used in IP at 7.5 µg / mL | Gift from Michael N. Hall, Basel, Switzerland (Molle, 2006). Generated according to van Slegtenhorst et al. (1998). | N/A |
| TSC1 #2 | used in IP at 7.5 µg / mL | Thermo Fisher Scientific (Invitrogen) | Cat# 37-0400; RRID: AB_2533292 |
| TSC1 #3 | dilution for IB 1:1000; used in IP at 4 µg/mL | Cell Signaling | Cat# 6935; RRID: AB_10860420 |
| TSC2 | dilution for IB 1:1000; dilution in IF 1:800; dilution for PLA 1:1600 | Cell Signaling | Cat# 4308; RRID: AB_10547134 |
| TSC2 #1 | used in IP at 7.5 µg / mL | Thermo Fisher Scientific (Invitrogen) | Cat# 37-0500; RRID: AB_2533293 |
| TSC2 #2 epitope maps to residues 1535 and 1784 of human TSC2 | used in IP at 7.5 µg / mL | Gift from Michael N. Hall, Basel, Switzerland (Molle, 2006). Generated according to van Slegtenhorst et al. (1998). | N/A |
| TSC2 #3 | used in IP at 7.5 µg / mL | Abcam | Cat# ab52936; RRID: AB_883283 |
| TUBA1B | dilution for IB 1:10000 | Abcam | Cat# ab108629; RRID: AB_10866252 |
| Bacterial and Virus Strains | | | |
| DB3.1 | | Thermo Fisher Scientific | Cat# 11782018 (discontinued) |
| DH5-alpha | | New England Biolabs | Cat# C2987H |

| Chemicals, Peptides, and Recombinant Proteins | | |
|---|---|---|
| Aprotin | Sigma-Aldrich | Cat# A1153 |
| Beta-Mercaptoethanol | Gibco | Cat# 21-985-023 |
| Bromophenol Blue | Sigma-Aldrich | Cat# B5525 |
| BSA (bovine serum albumin) | Carl Roth | Cat# 8076.5 |
| CHAPS (3-[(3-Cholamidopropyl)dimethylammonio]-1-propanesulfonate hydrate) | Sigma-Aldrich | Cat# 3023 |
| CHAPS (3-[(3-Cholamidopropyl)dimethylammonio]-1-propanesulfonate hydrate)<br>(for IPs in rat brain tissue) | Roth | Cat# 1479.3 |
| Complete Protease Inhibitor Cocktail | Sigma-Aldrich | Cat# D27802 |
| DABCO (1,4-diazabicyclo[2.2.2]octane) | Merck | Cat# 11836153001 |
| DMEM (Dulbecco's Modified Eagle's Medium) w: 4.5 g/L Glucose, w/o: L-Glutamine, w: Sodium pyruvate, w: 3.7 g/L NaHCO3 | PAN | Cat# P04-03600 |
| DMEM (Dulbecco's Modified Eagle's Medium) used for PLA experiments in Figure 3A | Thermo Fisher Scientific | Cat# 41965-039 |
| DMSO (dimethyl sulfoxide) | Sigma-Aldrich | Cat# D2650 |
| Doxycycline | Sigma-Aldrich | Cat# D3447 |
| Duolink™ In Situ Mounting Medium with DAPI | Sigma-Aldrich | Cat# DUO82040 |
| Dynabeads Protein G for Immunoprecipitation | Thermo Fisher Scientific | Cat# 10009D |
| FBS (fetal bovine serum) | Gibco | Cat# 10270106 |
| FBS (fetal bovine serum) | Sigma-Aldrich | Cat# F9665 |
| Glycerol | Sigma-Aldrich | Cat# G5516 |
| Glycine | Sigma-Aldrich | Cat# G7126 |
| HEPES ((4-(2-hydroxyethyl)-1-piperazineethanesulfonic acid) | Life technologies | Cat# 15630080 |
| HBSS (Hank's Balanced Salt Solution) w/o: Phenol red, w: Ca and Mg, w: 0.35 g/L NaHCO3 | PAN | Cat# P04-32505 |
| Hoechst 33342 (dilution in IF: 1:100.000) | Invitrogen | Cat# H3570 |
| Insulin | Sigma-Aldrich | Cat# I1882 |
| IGEPAL CA-630 (NP40) | Sigma-Aldrich | Cat# I8896 |
| Imidazole | Sigma-Aldrich | Cat# I0250 |

| | | |
|---|---|---|
| KCl (potassium chloride) | Sigma-Aldrich | Cat# P9541 |
| Leupeptin | Sigma-Aldrich | Cat# 103476-89-7 |
| L-glutamine | Gibco | Cat# 25030024 |
| L-analyl-glutamine | Gibco | Cat# 25030081 |
| Methanol | Klinipath | Cat# 4063-9005 |
| MgCl2 (magnesium chloride) | Sigma-Aldrich | Cat# M2670 |
| Mowiol 4-88 | Carl Roth | Cat# 07131 |
| NaCl (sodium chloride) | Sigma-Aldrich | Cat# S7653 |
| NPG (n-propyl-gallate) | VWR | Cat# EM8.20599.0500 |
| PBS (phosphate-buffered saline) | PAN | Cat# P04-36500 |
| PBS (phosphate-buffered saline) for non-sterile washing | Biochrom | Cat# L182-50 |
| Penicillin/Streptomycin | Gibco | Cat# 15140122 |
| Pepstatin A | Sigma-Aldrich | Cat# 26305-03-3 |
| PhosSTOP™ for IPs in brain tissue | Merck | Cat# 4906837001 |
| Phosphatase Inhibitor Cocktail 2 | Sigma-Aldrich | Cat# P5726 |
| Phosphatase Inhibitor Cocktail 3 | Sigma-Aldrich | Cat# P0044 |
| Pierce™ 16% Formaldehyde (w/v), Methanol-free | Thermo Fisher Scientific | Cat# 28908 |
| Polyacrylamide | Bio-Rad | Cat# 161-0159 |
| Polybrene | Sigma-Aldrich | Cat# H9268 |
| Prolong Gold antifade reagent with 4′,6-Diamidin-2-phenylindol (DAPI) | Thermo Fisher Scientific | Cat# P36935 |
| Protein G sepharose beads | GE | Cat# 17061801 |
| Puromycin | Sigma-Aldrich | Cat# P8833 |
| PVDF (polyvinylidene difluoride) membrane | Millipore | Cat# IPVH00010 |
| Rapamycin | Calbiochem | Cat# 553210 |
| RIPA buffer (zebrafish lysis) | Merck | Cat# R0278 |
| SDS (sodium dodecyl sulfate) | Sigma-Aldrich | Cat# 71725 |
| Sodium deoxycholate | Sigma-Aldrich | Cat# 30970 |
| Sodium fluoride | Sigma-Aldrich | Cat# 7681-49-4 |

| | | |
|---|---|---|
| Sodium orthovanadate | Sigma-Aldrich | Cat# 13721-39-6 |
| Sodium pyrophosphate | Sigma-Aldrich | Cat# 13472-36-1 |
| Sucrose | Sigma-Aldrich | Cat# S2395 |
| Transfectin | Biorad | Cat# 1703350 |
| TRIS base (tris(hydroxymethyl)aminomethane) | VWR | Cat# A1086.5000 |
| Triton X-100 | Sigma-Aldrich | Cat# 93443 |
| Tryphan Blue | Gibco | Cat# 15250061 |
| Trypsin | Gibco | Cat# 15400054 |
| Tween-20 | MP Biomedicals | Cat# 11TWEEN201 |
| Critical Commercial Assays | | |
| Bio-Rad Protein Assay Dye Reagent Concentrate | Bio-Rad | Cat# 500-0006 |
| Duolink In Situ Red Starter Kit Mouse/Rabbit | Sigma-Aldrich | Cat# DUO92008 |
| Duolink® In Situ PLA® Probe Anti-Rabbit PLUS Affinity purified Donkey anti-Rabbit IgG (H+L) | Sigma-Aldrich | Cat# DUO92002 |
| Duolink® In Situ PLA® Probe Anti-Mouse MINUS Affinity purified Donkey anti-Mouse IgG (H+L) | Sigma-Aldrich | Cat# DUO92004 |
| JetPEI | Poly-Plus | Cat# 101-40N |
| Lipofectamine 3000 Transfection Reagent | Thermo Fisher Scientific | Cat# L3000015 |
| Lipofectamine RNAiMAX Transfection Reagent | Thermo Fisher Scientific | Cat# 13778150 |
| MidiPrepKit NUCLEOBOND XTRA MIDI | Macherey-Nagel | Cat# 740410.50 |
| Pierce BCA protein assay kit | Thermo Fisher Scientific | Cat# 23225 |
| Pierce ECL Western Blotting Substrate | Thermo Fisher Scientific | Cat# 32209 |
| SuperSignal West FEMTO Maximum Sensitivity Substrate | Thermo Fisher Scientific | Cat# 34095 |
| Trans-Lentiviral shRNA Packaging Mix | Dharmacon | Cat# TLP5912 |
| Deposited Data | | |
| Invasive breast cancer (The Cancer Genome Atlas, TCGA, provisional) | www.cbioportal.org | N/A |
| *TSC1 RNA* expression data | www.kmplot.com | probeID: 209390_at |

| | | |
|---|---|---|
| *TSC2 RNA* expression data | www.kmplot.com | probeID: 215735_s_at |
| *G3BP1* RNA expression data | www.kmplot.com | probeID: 225007_at |
| G3BP1 protein expression data | www.kmplot.com | probeID: Q13283 |
| Experimental Models: Cell Lines | | |
| HEK293T | Thien et al. (2015) | N/A |
| HEK293-$\beta_2$AR | Lavoie et al. (2002) | N/A |
| HeLa alpha Kyoto | Thedieck et al. (2007) | N/A |
| MCF7 ACC115 | DSMZ | Cat# ACC115; RRID: CVCL_0031 |
| MCF7 Control | This Paper | N/A |
| MCF7 G3BP1 KO | This Paper | N/A |
| MCF7 GFP-LC3 | Gift from Joern Dengjel, Fribourg, Switzerland | N/A |
| MCF7 shControl | This paper | N/A |
| MCF7 shG3BP1 #1 | This paper | N/A |
| MCF7 shG3BP1 #2 | This paper | N/A |
| MDA-MB-231 | ATCC | Cat# HTB-26; RRID: CVCL_0062 |
| MDA-MB-231 TSC Control | This Paper | N/A |
| MDA-MB-231 TSC2 KO | This Paper | N/A |
| MDA-MB-231 shControl | This paper | N/A |
| MDA-MB-231 shG3BP1 #1 | This paper | N/A |
| MDA-MB-231 shG3BP1 #2 | This paper | N/A |
| Experimental Models: Organisms/Strains | | |
| AB Danio rerio | Zebrafish International Resource Center | Cat# ZL1; RRID: ZIRC_ZL1 |
| Wistar Cmd:(WI)WU rats | Mossakowski Medical Research Centre Polish Academy of Sciences | N/A |

| Recombinant DNA | | |
|---|---|---|
| bFos-myc-LC151 | Gift from Qingming Luo, Wuhan, China(Chu et al., 2009) | N/A |
| bJun-HA-LN151 | Gift from Qingming Luo, Wuhan, China(Chu et al., 2009) | N/A |
| lentiGuide-Puro | Sanjana et al. (2014) | RRID: Addgene_52963 |
| pCW-Cas9-Blast | Sanjana et al. (2014) | RRID: Addgene_83481 |
| pGW-myc-LC151 | Stefan Pusch (Weiler et al., 2014) | N/A |
| pGW-HA-LN151 | Stefan Pusch (Weiler et al., 2014) | N/A |
| pGW-myc-LC151-G3BP1 | This paper | N/A |
| pGW-myc-LC151-G3BP1 1-182 | This paper | N/A |
| pGW-myc-LC151-G3BP1 183-332 | This paper | N/A |
| pGW-myc-LC151-G3BP1 333-466 | This paper | N/A |
| pGW-myc-LC151-G3BP2 | This paper | N/A |
| pGW-HA-LN151-LAMP1 | This paper | N/A |
| pGW-HA-LN151-LAMP2 | This paper | N/A |
| pGW-HA-LN151-mTOR | This paper | N/A |
| pGW-HA-LN151-TSC2 | This paper | N/A |
| pEGFP-C-TSC2 | This paper | N/A |
| pEGFP-C (derivate of pDEST with a C-terminal EGFP tag) | Stefan Pusch | NA |
| pDEST | Stefan Pusch; Clone repository of the DKFZ Genomics and Proteomics Core Facility (GPCF) | N/A |

| | | |
|---|---|---|
| pENTR221-G3BP1 | Clone repository of the DKFZ Genomics and Proteomics Core Facility (GPCF) | CloneId: 182373397 |
| pENTR223-G3BP2 | Clone repository of the DKFZ Genomics and Proteomics Core Facility (GPCF) | CloneId: 192451551 |
| pENTR221-LAMP1 | Clone repository of the DKFZ Genomics and Proteomics Core Facility (GPCF) | CloneId: 193137117 |
| pENTR221-LAMP2 | Clone repository of the DKFZ Genomics and Proteomics Core Facility (GPCF) | CloneId: 115072391 |
| psPAX2 | Shalem et al. (2014) | RRID: Addgene_12260 |
| pMD2.G | Shalem et al. (2014) | RRID: Addgene_12259 |
| R777-E138 Hs.MTOR-nostop | Gift from Dominic Esposito, Addgene | Cat# 70422; RRID: Addgene_70422 |
| R777-E356 Hs.TSC2-nostop | Gift from Dominic Esposito, Addgene | Cat# 70640; RRID: Addgene_70640 |
| siControl (ON-TARGET plus Non-targeting Pool) | Dharmacon | Cat# D-001810-10-05 |
| siG3BP1 pool (ON-TARGET plus Human G3BP1 siRNA – SMART pool) | Dharmacon | Cat# L-012099-00-0020 |
| siG3BP2 pool (ON-TARGET plus Human G3BP2 siRNA – SMART pool) | Dharmacon | Cat# L-015329-01-0020 |
| siG3BP1 (siGENOME) (used for PLA experiments in Figure 3A) | Dharmacon | Cat# M-012099-02-0005 |
| anti-Luc siRNA-1 | Dharmacon | Cat# D-002050-01-20 |
| TRIPZ Inducible Lentiviral Human G3BP1 shRNA (F6 = shG3BP1 #1) | Dharmacon | Cat# RHS4696-200750396; CloneId: V3THS_329105 |

| | | |
|---|---|---|
| TRIPZ Inducible Lentiviral Human G3BP1 shRNA (H11 = shG3BP1 #2) | Dharmacon | Cat# RHS4696-200753099; CloneId: V3THS_329104 |
| TRIPZ Inducible Lentiviral Non-silencing shRNA Control | Dharmacon | Cat# RHS4743 |
| Software and Algorithms | | |
| Adobe Photoshop version CS5.1 | Adobe Systems Incorporated | RRID: SCR_014199 |
| Cell Profiler version 3.1.5 | Broad Institute of Harvard and MIT (www.cellprofiler.org) | RRID: SCR_007358 |
| CGDS-R package version 1.2.6 | https://github.com/cBioPortal/cgdsr | N/A |
| Dell Statistica version 13 | Dell Inc. | N/A |
| Fiji version 1.49v and 1.52p | ImageJ | RRID: SCR_002285 |
| GraphPad Prism version 7.04 and 8.0 | GraphPad Software | RRID: SCR_002798 |
| ImageJ version 1.50b | ImageJ | RRID: SCR_003070 |
| Image Lab version 5.2.1 | Bio-Rad | RRID: SCR_014210 |
| ImageQuant TL version 8.1 | GE Healtcare | RRID: SCR_014246 |
| Image Studio Lite Version 5.2 | Li-Cor | RRID: SCR_013715 |
| NIS Elements version 4.13.04 | Nikon | RRID: SCR_014329 |
| TScratch | www.cse-lab.ethz.ch/software.html (ETH Zürich;(Geback et al., 2009) | RRID: SCR_014282 |
| ZEN2012 blue edition | Zeiss | N/A |
| Oligonucleotides | | |
| AttB1 GGGGACAAGTTTGTACAAAAAAGCAGGCTCCACC | Stefan Pusch | N/A |
| AttB2 GGGGACCACTTTGTACAAGAAAGCTGGGTT | Stefan Pusch | N/A |
| G3BP1_B1 CAAAAAAGCAGGCTCCACCATGGTGATGGAGAAGCCTAGTC | Stefan Pusch | N/A |

| | | |
|---|---|---|
| G3BP1_182_B2o CAAGAAAGCTGGGTTGTCATTACTGACAACTGCCTGATC | Stefan Pusch | N/A |
| G3BP1_183_B1 CAAAAAAGCAGGCTCCACCATGGAAGAACATTTAGAGGAGCCTG | Stefan Pusch | N/A |
| G3BP1_332_B2o CAAGAAAGCTGGGTTTCTTCGGGGTTCAATGTCAC | Stefan Pusch | N/A |
| G3BP1_333_B1 CAAAAAAGCAGGCTCCACCATGGTGAGACACCCTGACAG | Stefan Pusch | N/A |
| G3BP1_B2o CAAGAAAGCTGGGTTCTGCCGTGGCGCAAG | Stefan Pusch | N/A |
| sgTSC2-Exon 2-F 5'-CACCGACGGAGTTTATCATCACCG | Invitrogen | N/A |
| sgTSC2-Exon 2-R 5'-AAACCGGTGATGATAAACTCCGTC | Invitrogen | N/A |
| sgG3BP1-Exon 3-F 5'-CACCGAAGCCAGCAGATGCAGTCTA | Invitrogen | N/A |
| sgG3BP1-Exon 3-R 5'-AAACTAGACTGCATCTGCTGGCTTC | Invitrogen | N/A |
| Zebrafish g3bp1 (fwd): 5'-ATGGTGATGGAGAAGCCAAG-3' | Invitrogen | N/A |
| Zebrafish g3bp1(rev): 5'- 5'-TTCCATTGTTGTCCAGTCCA-3' | Invitrogen | N/A |
| Zebrafish β-actin (fwd): 5'-CGAGCAGGAGATGGGAACC-3' | Invitrogen | N/A |
| Zebrafish β-actin (rev): 5'-CAACGGAAACGCTCATTGC-3' | Invitrogen | N/A |
| Other | | |
| 24 well plates | TPP | Cat# 92424 |
| 6 cm cell culture dish | Greiner bio-one | Cat# 628160 |
| 10 cm cell culture dish | TPP | Cat# 93100 |
| 15 cm cell culture dish | TPP | Cat# 93150 |
| 70 Ti Rotor for ultracentrifuge | Beckman Coulter | Cat# 337922 |
| AxioObserver Z1 | Zeiss | N/A |

| | | |
|---|---|---|
| Beckman Optima L-70K Ultracentrifuge | Beckman Coulter | Cat# 8043-30-1187 |
| ChemiDoc XRS+ | Bio Rad | Cat# 1708265 |
| Cover Glass | VWR international | Cat# 631-0130 |
| E-plate 16 for RTCA | ACEA Biosciences, Inc. | Cat# 05469813001 |
| FUSION FX7 with the DarQ-9 camera | Vilber | N/A |
| ibidi culture-insert 2 well | ibidi | Cat# 80209 |
| iBlot gel transfer stacks nitrocellulose membrane | Thermo Fisher Scientific | Cat# IB301002 |
| LAS-4000 mini camera system | GE Healthcare | N/A |
| LAS-4000 camera system | GE Healthcare | N/A |
| Microscope slides | Thermo Fisher Scientific | Cat# 4951PLUS4 |
| Mini-PROTEAN® Tetra Vertical Electrophoresis Cell system | Bio Rad | Cat# 1658029FC |
| Nikon ECLIPSE Ti-E/B | Nikon | N/A |
| NuPage MES SDS running buffer | Thermo Fisher Scientific | Cat# NP0002 |
| NuPage Novex 10% Bis-Tris gel | Thermo Fisher Scientific | Cat# NP0302BOX |
| Odyssey blocking buffer | Li-cor | Cat# 927-40000 |
| Odyssey 2.1 imaging system | Li-Cor, USA | N/A |
| RTCA Control Unit with RTCA Software | ACEA Biosciences, Inc. | Cat# 05454417001 |
| RTCA DP Analyzer | ACEA Biosciences, Inc. | Cat# 05469759001 |

#### Data availability

All data are available from the corresponding authors upon reasonable request.

#### References


Chu, J., Zhang, Z., Zheng, Y., Yang, J., Qin, L., Lu, J., Huang, Z.L., Zeng, S., and Luo, Q. (2009). A novel far-red bimolecular fluorescence complementation system that allows for efficient visualization of protein interactions under physiological conditions. Biosens Bioelectron *25*, 234-239.

Geback, T., Schulz, M.M., Koumoutsakos, P., and Detmar, M. (2009). TScratch: a novel and simple software tool for automated analysis of monolayer wound healing assays. Biotechniques *46*, 265-274.

Lavoie, C., Mercier, J.F., Salahpour, A., Umapathy, D., Breit, A., Villeneuve, L.R., Zhu, W.Z., Xiao, R.P., Lakatta, E.G., Bouvier, M., *et al.* (2002). Beta 1/beta 2-adrenergic receptor heterodimerization regulates beta 2-adrenergic receptor internalization and ERK signaling efficacy. J Biol Chem *277*, 35402-35410.

Molle, K.-D. (2006). Regulation of the mammalian target of rapamycin complex 2 (mTORC2). In Department Biozentrum (Basel: University of Basel), pp. 92.

Sanjana, N.E., Shalem, O., and Zhang, F. (2014). Improved vectors and genome-wide libraries for CRISPR screening. Nat Methods *11*, 783-784.

Shalem, O., Sanjana, N.E., Hartenian, E., Shi, X., Scott, D.A., Mikkelson, T., Heckl, D., Ebert, B.L., Root, D.E., Doench, J.G., *et al.* (2014). Genome-scale CRISPR-Cas9 knockout screening in human cells. Science *343*, 84-87.

Thedieck, K., Polak, P., Kim, M.L., Molle, K.D., Cohen, A., Jeno, P., Arrieumerlou, C., and Hall, M.N. (2007). PRAS40 and PRR5-like protein are new mTOR interactors that regulate apoptosis. PLoS One *2*, e1217.

Thien, A., Prentzell, M.T., Holzwarth, B., Klasener, K., Kuper, I., Boehlke, C., Sonntag, A.G., Ruf, S., Maerz, L., Nitschke, R., *et al.* (2015). TSC1 activates TGF-beta-Smad2/3 signaling in growth arrest and epithelial-to-mesenchymal transition. Dev Cell *32*, 617-630.

van Slegtenhorst, M., Nellist, M., Nagelkerken, B., Cheadle, J., Snell, R., van den Ouweland, A., Reuser, A., Sampson, J., Halley, D., and van der Sluijs, P. (1998). Interaction between hamartin and tuberin, the TSC1 and TSC2 gene products. Hum Mol Genet *7*, 1053-1057.

Weiler, M., Blaes, J., Pusch, S., Sahm, F., Czabanka, M., Luger, S., Bunse, L., Solecki, G., Eichwald, V., Jugold, M., *et al.* (2014). mTOR target NDRG1 confers MGMT-dependent resistance to alkylating chemotherapy. Proc Natl Acad Sci U S A *111*, 409-414.
